# Structural basis of DNA N^6^-adenine methylation in eukaryotes

**DOI:** 10.1101/2025.07.08.663716

**Authors:** Bei Nan, Wentao Yang, Lanheng Nie, Dantong Wang, Mengyang Xu, Junhua Niu, Yuanyuan Wang, Wenjia Guan, Zhao Chen, Guangchao Zhuang, Shan Gao, Yifan Liu

**Author notes:** These authors contributed equally (Bei Nan, Wentao Yang, Lanheng Nie, Dantong Wang, Mengyang Xu, Junhua Niu, Yuanyuan Wang). Corresponding author: Shan Gao; Yifan Liu.

## Abstract

As a transcription-associated epigenetic mark abundantly present in many unicellular eukaryotes, DNA N^6^-adenine methylation (6mA) is maintained by the AMT1 complex. However, the structure-function relationship of the AMT1 complex remains to be fully elucidated. Here, we employ AlphaFold3 (AF3) for de novo structural modeling of the AMT1 complex, complemented by molecular dynamics simulations. Functional relevance of key structural features, especially those of the AMT1 ternary complex with the double-stranded DNA (dsDNA) substrate and the cofactor, are validated by extensive mutagenesis, both *in vivo and in vitro*. Our analysis reveals the structural basis for DNA substrate recognition, base flipping, and catalysis in this prototypical eukaryotic DNA 6mA-methyltransferase. It also allows us to delineate the reaction pathway for processive methylation involving translocation of the AMT1 complex along the dsDNA. As the active site is conserved across the MT-A70 family of eukaryotic N^6^-adenine methyltransferases, the structural insight will facilitate rational design of specific inhibitors.

## Introduction

As a base modification, N^6^-adenine methylation occurs in both DNA and RNA, where it is referred to as 6mA and m6A, respectively (Figure 1A). While 6mA is well-established in prokaryotes, its widespread presence as an epigenetic mark in eukaryotes has only recently been recognized^1, 2, 3, 4^. Currently, 6mA is best characterized in the unicellular eukaryote *Tetrahymena thermophila*^4, 5, 6, 7, 8^. In eukaryotes, N^6^-adenine methylation is catalyzed by MT-A70 family methyltransferases (MTases), which are classified into several clades with distinct structures and functions^9, 10, 11^. METTL3 and METTL14, the founding members of the family, form a heterodimer that deposits m6A in mRNA—a well-known epi-transcriptomic mark^12, 13^. Another MT-A70 MTase, AMT1, is required for semiconservative transmission of 6mA in double-stranded DNA (dsDNA)^5, 7, 14, 15^. Two AMT1-containing complexes, defined by their mutually exclusive subunits, AMT6 and AMT7, coordinate in 6mA transmission in *Tetrahymena*: the AMT7 complex, with higher activity and processivity, is important for transcription-associated methylation, while the less active and processive AMT6 complex is primarily involved in replication-coupled methylatioN^6^ (Figure 1B).

**Figure 1.**
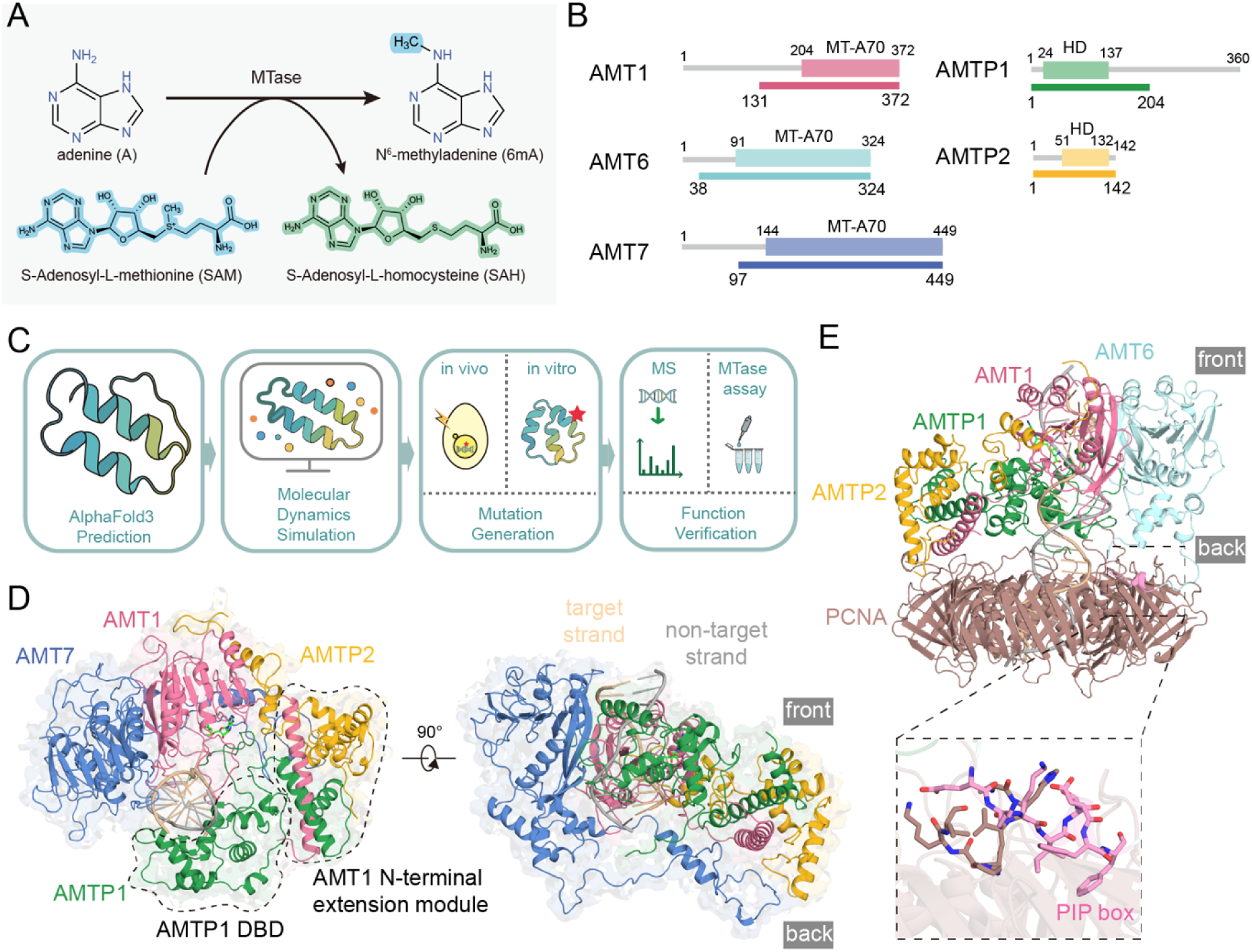
Overall structural organization of the AMT1 holo-complex. A. Methyl transfer from S-adenosyl-L-methionine (SAM) to adenine (A), generating N^6^-methyladenine (6mA) and S-adenosyl-L-homocysteine (SAH). B. Domain architecture of the AMT1 complex subunits: AMT1, AMT6, AMT7, AMTP1, and AMTP2. HD, homeobox-like domain. The colored bars below represent sequences used in structural modeling. C. Workflow for structural modeling and functional verification for the AMT1 complex. After the initial AlphaFold3 (AF3) prediction, critical models were further optimized by molecular dynamics (MD) simulations. Key residues underpinning the predicted structural features were systematically mutated, and their functional relevance was verified by various *in vitro* and *in vivo* assays. D. Front (left) and side (right) views of the AMT7 ternary complex (AMT7-hemi-SAM). The representative AF3 model was optimized by extended MD simulations. The AMTP1 DNA-binding domain (DBD) and the AMT1 N-terminal extension module are circled by dotted lines. E. Side view of the AMT6 ternary complex (AMT6-hemi-SAM) with tandemly loaded PCNA. Expanded view: interactions between PCNA and the PCNA-interaction protein (PIP) box (pink) of AMT6.

Extensive structural studies of prokaryotic DNA MTases—including both 5mC (5-methyl cytosine)^16, 17^ and 6mA-MTases^18^—reveal a conserved MTase fold at the active site, which can bind both the substrate (cytosine or adenine) and the cofactor (S-adenosyl methionine [SAM] or S-adenosyl homocysteine [SAH])^19, 20, 21^. Flipping of the target base out of the dsDNA helix and into the active site is a universal mechanism for DNA MTase catalysis^22^. The MT-A70 family of eukaryotic 6mA/m6A-MTases are classified as hetero-dimeric beta class amino MTases^20, 21^. The structure of the METTL3-METTL14 apo-complex demonstrates a division of labor between the cofactor-binding catalytic METTL3 subunit and the non-catalytic METTL14 subunit^23, 24, 25^, analogous to a homo-dimeric prokaryotic 6mA MTase EcoP15I^26, 27^. Similarly, the structure of the hetero-tetrameric AMT1 apo-complex features AMT1 as the catalytic subunit and AMT7 as the non-catalytic subunit, alongside AMTP1 and AMTP2 as auxiliary subunits^28, 29^. Although recent structural analysis of the AMT1 complex associated with a DNA mimic—the Overcoming Classical Restriction (OCR) protein from bacteriophage T7—has provided important clues for substrate binding^30^, no structure of the AMT1 holo-complex engaging its cognate substrate has been resolved. A gap remains in understanding the molecular basis for its substrate recognition and catalysis.

The past few years have witnessed dramatic advances in AI-based protein structure prediction, marked by breakthroughs such as AlphaFold2 and RoseTTAFold^31, 32^. The advent of AlphaFold3 (AF3) and RoseTTAFold all-atom extends accurate modeling to other biomolecules, including nucleic acids and small-molecule ligands (with covalent modifications such as 6mA treated as a special case)^33, 34^. In this study, we integrate AF3 predictions with molecular dynamics (MD) simulations to model the AMT1 complex, especially the ternary holo-complex bound to the dsDNA and cofactor (Figure 1B). Our structural modeling, supported by extensive in vivo and in vitro mutagenesis, provides mechanistic insight into substrate recognition, base flipping, and catalysis in this prototypical eukaryotic DNA 6mA-MTase. Notably, we uncover the structural basis of its substrate specificity for the ApT dinucleotide and its strong preference for the hemi-methylated state. Our AF3 predictions also demonstrate substantial conformational heterogeneity in the AMT1 complex, underscoring the structural plasticity intrinsic to the dynamic process of DNA methylation. By systematic analysis of the conformational space of the AMT1 complex, we delineate a reaction pathway for processive DNA methylation that involves continuous translocation of the AMT1 complex along the dsDNA. Given the conservation of the active site across the MT-A70 family of eukaryotic 6mA/m6A MTases, our findings offer structural insights that will facilitate the rational design of small molecule inhibitors, including those targeting METTL3-METTL14 in cancer and leukemia^35, 36^.

## Results

### Structural modeling of the AMT1 complex

In *Tetrahymena*, the AMT1 complex is primarily involved in 6mA maintenance methylation (hemi-methylated ApT → full-methylation), but also capable of *de novo* methylation (unmodified ApT → hemi-methylation), albeit with reduced activity^5, 6, 7, 14^. We used AF3 for structural modeling of the AMT1 apo-complex (without DNA), the binary holo-complex (with only DNA), and the ternary holo-complex (with both DNA and SAM/SAH) (Figure 1B, C, Figure S1A, B, Supplemental Table S1). For the holo-complex, we focused on dsDNA substrates featuring a palindromic sequence and a central ApT duplex in one of the three methylation states, which correspond to the reactants and products of *de novo* and maintenance methylation (Figure S1B). We also modeled the complex with AMT6 or AMT7 as the alternative subunit (Figure S1A). By evaluating 28 distinct input settings with 1,000 seeds each (generating 5 models per seed), we produced a total of 140,000 models (Figure 1C, Figure S1A, Supplemental Table S1). This comprehensive approach allowed us to identify high-confidence structures while exploring conformational heterogeneity of the complex.

We subsequently used extensive molecular dynamics (MD) simulations to optimize and validate critical AF3 models, assess their stability, and explore their local conformational landscape (Figure 1C). Functional implications for key structural features were then experimentally verified, via *in vitro* and *in vivo* mutagenesis. This was followed by MTase assays of the reconstituted AMT1 complex, alongside mass spectrum (MS) and PacBio sequencing-based quantification of 6mA levels in *Tetrahymena* mutants (Figure 1C). This integrated approach allowed us to identify physically and biologically relevant interactions and establish critical structure-function relationships.

### Overall structural organization of the AMT1 complex

Published studies of the AMT1 apo-complex have shed light on its structural core, which comprises the heterodimeric AMT1 and AMT7 MT-A70 domains flanked by the AMT1 N-terminal extension module^28, 29, 30^. However, the DNA-binding domain (DBD) of AMTP1, long hypothesized to interact with the substrate, is not resolved in these structures^28, 29, 30^. Our AF3 modeling of the AMT1 apo-complex predicted a high-confidence and precise local structure for the AMTP1 DBD (Figure S1C). However, the AMTP1 DBD exhibited dispersed global arrangements: it was positioned at variable distances and orietations relative to the structural core, to which it was tethered only by a long linker (Figure S1C). By contrast, a definite global arrangement emerged in AF3 models of the AMT1 holo-complex, in which the protein tetramer links into a torus with the dsDNA passing through the central hole (Figure 1D, Figure S1D, S2A-F). The AMTP1 DBD is confined to the DNA major groove, making multiple well-defined contacts with the structural core (Figure 1D).

Within the AMT1 holo-complex, the catalytic subunit AMT1 engages the target strand of the dsDNA at the ApT site. Based on the target strand directionality, we define two faces of the AMT1 holo-complex: the front facing the 3’ end and the back facing the 5’ end (Figure 1D, side view). Looking down at the front face, the AMTP1 DBD, AMT6/AMT7 MT-A70, AMT1 MT-A70, and AMT1 N-terminal extension module are arranged clockwise (Figure 1D, front view). The AMT1 N-terminal extension module— comprising the AMT1 N-terminal helix, the AMTP1 C-terminal helix-turn-helix-like domain, and AMTP2—is not directly involved in DNA binding (Figure 1D, front view). The remaining three interconnected domains—the AMTP1 DBD, AMT6/AMT7 MT-A70, and AMT1 MT-A70—wrap tightly around the dsDNA (Figure 1D, front view). Both the adenine-binding and cofactor-binding pockets, formed solely by AMT1, open at the front (Figure 1D, front view). By contrast, sequences divergent in AMT6 and AMT7— especially their long N-terminal tails and internal loops—protrude from the back (Figure 1D, side view). These AMT6- and AMT7-specific regions are implicated in regulating and targeting their MTase activities, as replication-coupled and transcription-associated methylation are preferentially performed by the AMT6 and AMT7 complexes, respectively^6^. Indeed, the Proliferating Cell Nuclear Antigen (PCNA)-interacting protein (PIP) box of AMT6 resides within a long loop divergent from AMT7 at the back (Figure 1E). This spatial arrangement positions the AMT6 PIP box for direct insertion into a conserved pocket at the front of the tandemly loaded replication sliding clamp (Figure 1E).

The interface between the AMT1 and AMT6/AMT7 MT-A70 domains is defined in the AMT1 apo-complex structures^28, 29^. We tracked two interatomic distances across this interface in our AF3 models of all AMT1 apo- and holo-complexes. These values tightly clustered in scatter plots, indicating a stable and homogeneous local conformation (Figure S1E). Nonetheless, substantial global conformational heterogeneity still existed in AF3 models of the AMT1 complex, reflecting its intrinsic structural dynamics. Systematic examinations revealed that the conformational heterogeneity was often attributable to variable interfaces between largely invariable structural domains (Figure S1F, Supplemental Table S3). Building on this insight, we developed several sets of parameters to monitor these variable interfaces within the AMT1 complex (Figure S1F, Supplemental Table S3). Each set of parameters tracked key distances across a specific interface, and the distributions of these values were mapped across all AF3 models. Conformational analyses based on each parameter set are detailed in the subsequent Results sections focusing on individual structural features. In the Discussion, the conformational heterogeneity of the AMT1 complex is recapitulated by analyzing a collection of relevant parameters using Uniform Manifold Approximation and Projection (UMAP).

### DNA binding by AMT1

DNA and RNA substrates are not present in previously solved structures of the AMT1 complex^28, 29^ and the METTL3-METTL14 complex^23, 24, 25^. Therefore, the molecular mechanisms for eukaryotic 6mA/m6A MTases remain to be fully elucidated. We first analyzed an AMT1 ternary complex containing hemi-methylated dsDNA and cofactor SAM (AMT7-hemi-SAM). The top 10 AF3 models, ranked by their ipTM scores (>0.9, indicating high-confidence predictions), were exclusively in the base flipping mode. These AF3 models were largely superimposable after pruning their flexible regions (Figure S2). We focused on one representative model, in which the unmodified adenine on the target strand was flipped out of the dsDNA helix and into the active site, corresponding to the enzyme-substrate (ES) complex for maintenance methylation^5, 6^. Starting from this structure, we performed extended MD simulations (1 μs), implemented in GROMACS using the latest CHARMM36 force field^37, 38, 39^. The system reached convergence, as most MD models from the late trajectories were also largely superimposable (Figure S3).

Here, we highlight three AMT1 loops crucial for substrate binding: the *Major Groove Insertion Loop*, the *Minor Groove Insertion Loop*, and the *Regulation Loop* (Figure 2A). In the presence of dsDNA and SAM, they form well-defined structures that grip the target strand and enclose the extrahelical adenine (Figure 2A, Figure S4A). AMT1 and METTL3 closely align in both their primary sequences and 3D structures, exhibiting one-to-one correspondence across these substrate-binding loops (Figure S4B, C). Their sequence conservation and divergence illuminate the shared and unique features of adenine methylation in DNA and RNA. The AMT1 *Major Groove Insertion Loop*, which engages the extrahelical adenine as well as the major groove, corresponds to METTL3 *Gate Loop 1* but harbors a seven-amino-acid insertion (86-92: SSQPSRG) in its center (Figure 2B, Figure S4B, C). Notably, most of the AMT1 *Major Groove Insertion Loop* is unresolved in the published apo-complex structures^28, 29^. While it remained relatively homogenous in the AF3 models of the AMT1 ternary holo-complex, it exhibited highly variable conformations in the AMT1 binary holo-complex—where the target adenine mostly remained intrahelical—as well as in the AMT1 apo-complex (Figure S4A). These observations suggest that the AMT1 *Major Groove Insertion Loop* undergoes a disorder-to-order transition upon binding the dsDNA, especially the extrahelical adenine. The AMT1 *Minor Groove Insertion Loop* corresponds to the METTL3 *Interface Loop* but is divergent in its sequence (Figure 2C, Figure S4B, C). It is physically associated with the AMT7 *Multiple Interaction Loop*, which is partially conserved in AMT6 but divergent in METTL14 (Figure 2C, Figure S4B, D). The *Regulation Loop*, which controls the interaction between the extrahelical adenine and the cofactor, aligns with METTL3 *Gate Loop 2* (Figure 2D, Figure S4B, C).

**Figure 2.**
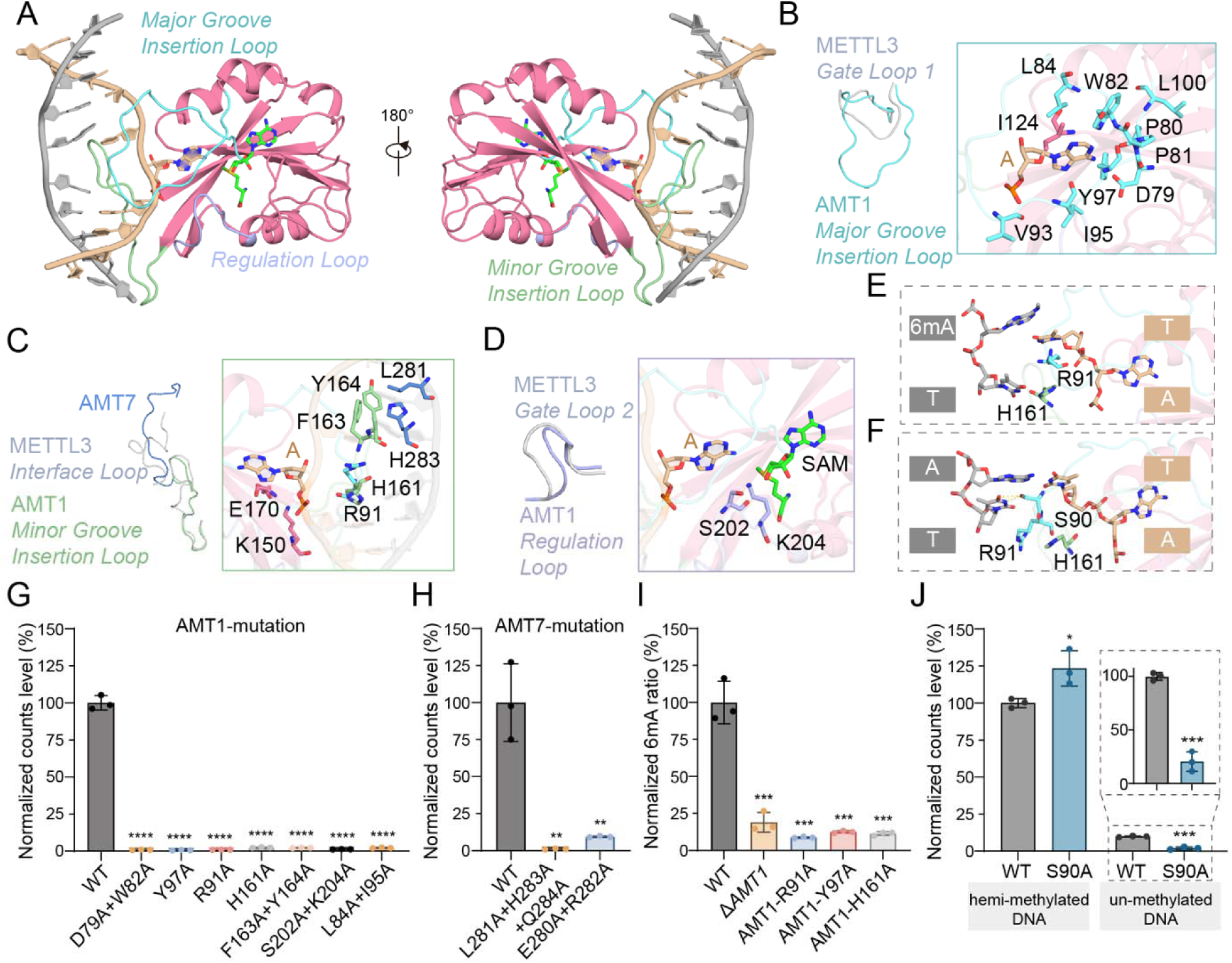
DNA binding by AMT1. A. Opposite views of DNA binding by AMT1. MD model of the AMT1 ternary complex (AMT7-hemi-SAM), base-flipping mode. Left/right: DNA major/minor groove facing outward. The three substrate-binding loops are colored and labeled. SAM is shown in the stick model. B. Close-up view of the AMT1 *Major Groove Insertion Loop*. Left: structural overlay of the AMT1 *Major Groove Insertion Loop* and METTL3 *Gate Loop 1*. Right: residues critical for forming the hydrophobic core and interacting with the extrahelical adenine. C. Close-up view of the AMT1 *Minor Groove Insertion Loop*. Left: structural overlay of the AMT1 *Minor Groove Insertion Loop* and the METTL3 *Interface Loop*. The AMT7 *Multiple Interaction Loop* and the METTL14 *Interface Loop* are also shown. Right: residues in the hydrophobic core (AMT1 F163, Y164 and AMT7 L281, H283) and critical for DNA binding (AMT1 R91, K150, and E170). D. Close-up view of the AMT1 *Regulation Loop*. Left: structural overlay of the AMT1 *Regulation Loop* and METTL3 *Gate Loop 2*. Right: residues critical for interacting with the extrahelical adenine and the cofactor SAM. E. Insertion of R91 and H161 into the base stack at the ApT duplex in the enzyme-substrate (ES) complex for maintenance methylation (AMT7-hemi-SAM). The non-target 6mA:T base pair is disrupted. F. Insertion of R91 and H161 into the base stack at the ApT duplex in the ES complex for *de novo* methylation (AMT7-un-SAM). The non-target 6mA:T pair is aligned for proper hydrogen bond formation. Yellow dashed line: hydrogen bond between AMT1 S90 and the orphan thymine on the non-target strand. G. In vitro MTase activity of the reconstituted AMT7 complex with the indicated point mutations of AMT1. The wildtype (WT) AMT7 complex was included as the positive control. Data are shown as mean±SD. Two-tailed Student’s t-test: \*\*\*\**P*< 0.0001. H. In vitro MTase activity of the reconstituted AMT7 complex with the indicated point mutations of AMT7. Two-tailed Student’s t-test: ** *P*< 0.01. I. Mass spectrometry quantification of the 6mA-to-A ratio (6mA/A, %) in *Tetrahymena* WT (positive control), Δ*AMT1* (negative control), and mutants with the indicated point mutations of AMT1. Two-tailed Student’s t-test: *** *P*< 0.001. J. In vitro MTase activity of the reconstituted AMT7 complex with the AMT1 S90A mutation. Both the hemi-methylated and un-methylated dsDNA substrates were tested. Two-tailed Student’s t-test: * *P*<0.05, *** *P*<0.001.

### Base flipping mechanisms

While base flipping is ubiquitous in DNA MTases, the angle can vary substantially^22^. In Dam and CcrM, the target base flips ∼90°^26, 40^; in M.HhaI and EcoP15I, the target base flips ∼180°^16, 17, 27^. To better understand the base flipping mechanism, we analyzed the MD trajectories of the AMT1 ternary complex. In the representative MD models, the target adenine flips out completely (∼180°) and inserts into a deep pocket of AMT1 (Figure 2A). Base flipping is stabilized by residues conserved in both AMT1 and METTL3 (Figure 2B-D, Figure S4C). The proximal portion of the AMT1 *Major Groove Insertion Loop*—containing DPPW (AMT1 79-82; METTL3 395-398) and Y97 (METTL3 Y406)—maintains close contact with the extrahelical adenine (Figure 2B). This structure is stabilized by a hydrophobic core comprising P80 (P396), P81 (P397), W82 (W398), L84 (I400), V93 (L402), I95 (L404), Y97 (Y406), and L100 (L409) from the AMT1 *Major Groove Insertion Loop*, as well as I124 (T433) connecting β3 and α3 (Figure 2B, Figure S4E). The extrahelical adenine also interacts with the AMT1 *Regulation Loop*, containing S202 (S511) and K204 (K513) (Figure 2D). Its Hoogsteen edge is buttressed against a chain of alternating acidic-basic residues within the AMT1 *Major Groove Insertion Loop* (D79 [D395]) and the *Regulation Loop* (K204 [K513]), and flanking the *Minor Groove Insertion Loop* (E170 [E481] and K150 [K459]) (Figure 2B-D). Interactions are extended to the sugar-phosphate moiety: AMT1 K150 (K459) contacts the 5’ phosphate, while I124 (T433) approaches the 2’ carbon of the sugar ring (Figure S4E). The switch from AMT1 I124 to METTL3 T433 accommodate the 2’-hydroxyl group present in RNA but not DNA.

The AMT1 *Major Groove Insertion Loop* and *Minor Groove Insertion Loop* engage the base stack around the ApT duplex, stretching the dsDNA along its helical axis and unwinding it (Figure 2A, E). The distal portion of the AMT1 *Major Groove Insertion Loop*—featuring seven amino acids (86-92: SSQPSRG) missing from METTL3 *Gate Loop 1* (Figure S4C)—inserts into the dsDNA from the major groove and, in concert with the AMTP1 DBD, expands the major groove substantially (Figure 2A, B, E). In parallel, the AMT1 *Minor Groove Insertion Loop* and the AMT7 *Multiple Interaction Loop*, stabilized by a hydrophobic core, insert into the dsDNA from the minor groove and expand it substantially (Figure 2A, C, E). Notably, the guanidino group of R91 within the *Major Groove Insertion Loop* and the imidazole group of H161 within the *Minor Groove Insertion Loop*—both planar—directly intercalate into the base stack (Figure 2E). Lastly, all these residues—critical for the interactions conducive to base flipping in dsDNA, but not in single-stranded RNA—are conserved in the AMT1 complex but divergent in the METTL3-METTL14 complex (Figure S4C, D, Supplemental File S1, S2).

### ApT recognition by AMT1

In *Tetrahymena*, 6mA occurs exclusively in the ApT dinucleotide; the AMT1 complex is critical for its maintenance methylation, but also capable of *de novo* methylation^5^. *In vitro* methylation by the reconstituted AMT1 complex also shows a strong bias for the ApT dinucleotide, with much higher activity for the hemi-methylated ApT than the unmodified substrate^5, 6, 7, 14^. To elucidate the structural features underpinning the substrate specificity, we closely examined the MD trajectories of the AMT1 ternary complex. The proximal portion of the AMT1 *Major Groove Insertion Loop* directly recognizes the extrahelical adenine (Figure 2B). Its distal portion, together with the AMT1 *Minor Groove Insertion Loop*, inserts into the base stack to mediate additional sequence-specific interactions with the ApT duplex (Figure 2E). R91 in the AMT1 *Major Groove Insertion Loop* is positioned in close proximity to the non-target 6mA and the target thymine (Figure 2E). Using AMT1 R91 as an anchor point, we tracked its distances to A/6mA in the ApT duplex (Figure S5A, B, Supplemental Table S3). For most AF3 models of the AMT6/AMT7 ternary complexes in the base flipping mode, the distance to the intrahelical A/6mA fluctuated around small values (5 Å), indicative of close contact and deep insertion (Figure S5A). In the corresponding MD trajectories, the distances also quickly converged on similar values (Figure S5B). Similarly, H161 in the AMT1 *Minor Groove Insertion Loop* can form hydrogen bonds with 6mA or thymine on the non-target strand (Figure 2E). We conclude that AMT1 R91 and H161 play critical roles in ApT duplex recognition by displacing the target adenine and stabilizing the remaining bases within the dsDNA.

Intriguingly, in the ES complex for maintenance methylation (AMT7-hemi-SAM), the non-target 6mA, while not flipped out, is also not properly paired with its complementary thymine base on the target strand (Figure 2E). Rather, these two intrahelical bases are buckled by the double insertion of AMT1 R91 and H161; their misalignment precludes hydrogen bond formation (Figure 2E). As 6mA can substantially weaken A:T base pairing^41, 42, 43^, its presence in the non-target A:T predisposes the duplex for disruption, thereby facilitating the major/minor groove insertion and base flipping by the AMT1 complex. By contrast, in the ES complex for *de novo* methylation (AMT7-un-SAM), the intrahelical adenine is generally aligned with its complementary thymine for standard base pairing (Figure 2F). The stable A:T pair compresses the dsDNA around the ApT duplex and disturbs the extrahelical adenine at the AMT1 active site (Figure 2F). These structure features of the AMT1 complex underpin its preferential methylation of hemi-6mApT over unmodified ApT as well as non-ApT sites.

We reconstituted the AMT1 complex incorporating mutations affecting these residues, including AMT1 D79A, W82A, D79A+W82A, Y97A, R91A, L84A+I95A (within the AMT1 *Major Groove Insertion Loop*); AMT1 H161A, F163A+Y164A (within the AMT1 *Minor Groove Insertion Loop*); AMT1 S202A+K204A (within the AMT1 *Regulation Loop*) (Figure 2G), and AMT7 L281A+H283A+Q284A, E280A+R282A (within the AMT7 *Multiple Interaction Loop*) (Figure 2H). *In vitro* MTase activity was abolished by all these mutations (Figure 2G, H). Some mutations were also introduced into *Tetrahymena*; MS analysis revealed diminished endogenous 6mA levels in these mutants (comparable to Δ*AMT1*) (Figure 2I). These results confirm the critical role of the AMT1 substrate-binding loops.

In the ES complex for maintenance methylation (AMT7-hemi-SAM), 6mA on the non-target strand, especially its N^6^-methyl group, makes contacts with the distal portion of the AMT1 *Major Groove Insertion Loop*, including S90 (Figure S5C). By contrast, in the ES complex for *de novo* methylation (AMT7-un-SAM), the orphan thymine on the non-target strand forms a stable hydrogen bond with the hydroxyl side chain of AMT1 S90 (Figure 2F). AMT1 S90, largely conserved in AMT1 (Supplemental File S1), can thus interact favorably with unmodified ApT, while accommodating hemi-6mApT. Indeed, the reconstituted AMT1 complex containing the S90A mutation exhibited drastically reduced *de novo* methylation activity; by contrast, the mutation resulted in a moderate increase in maintenance methylation (Figure 2J). Therefore, AMT1 S90 is crucial for the AMT1 complex to perform both *de novo* and maintenance methylation.

### AMT1 active site

The cofactor-binding pocket is conserved in AMT1 and METTL3^28, 29^. In the top ranked AF3 models for the AMT1 ternary complex, cofactor placement was highly consistent, with SAM and SAH similarly positioned (Figure S6A). SAM/SAH placement was not significantly altered during MD simulations (Figure 3A, Figure S6B). SAM/SAH placement in these predicted structures was all but identical to that in the solved structures of the AMT1 complex (7f4n)^28^ and the METTL3-METTL14 complex (5il2)^24^ (Figure 3A). However, the SAH conformation in another solved AMT1 complex structure (7yi8) is substantially different^29^ (Figure S6C). This discrepancy is surprising given that the cofactor-binding pocket is highly similar across all aforementioned structures (Figure 3A). We strongly favor our current cofactor placement, as it avoids the steric clashes encountered in the alternative model.

**Figure 3.**
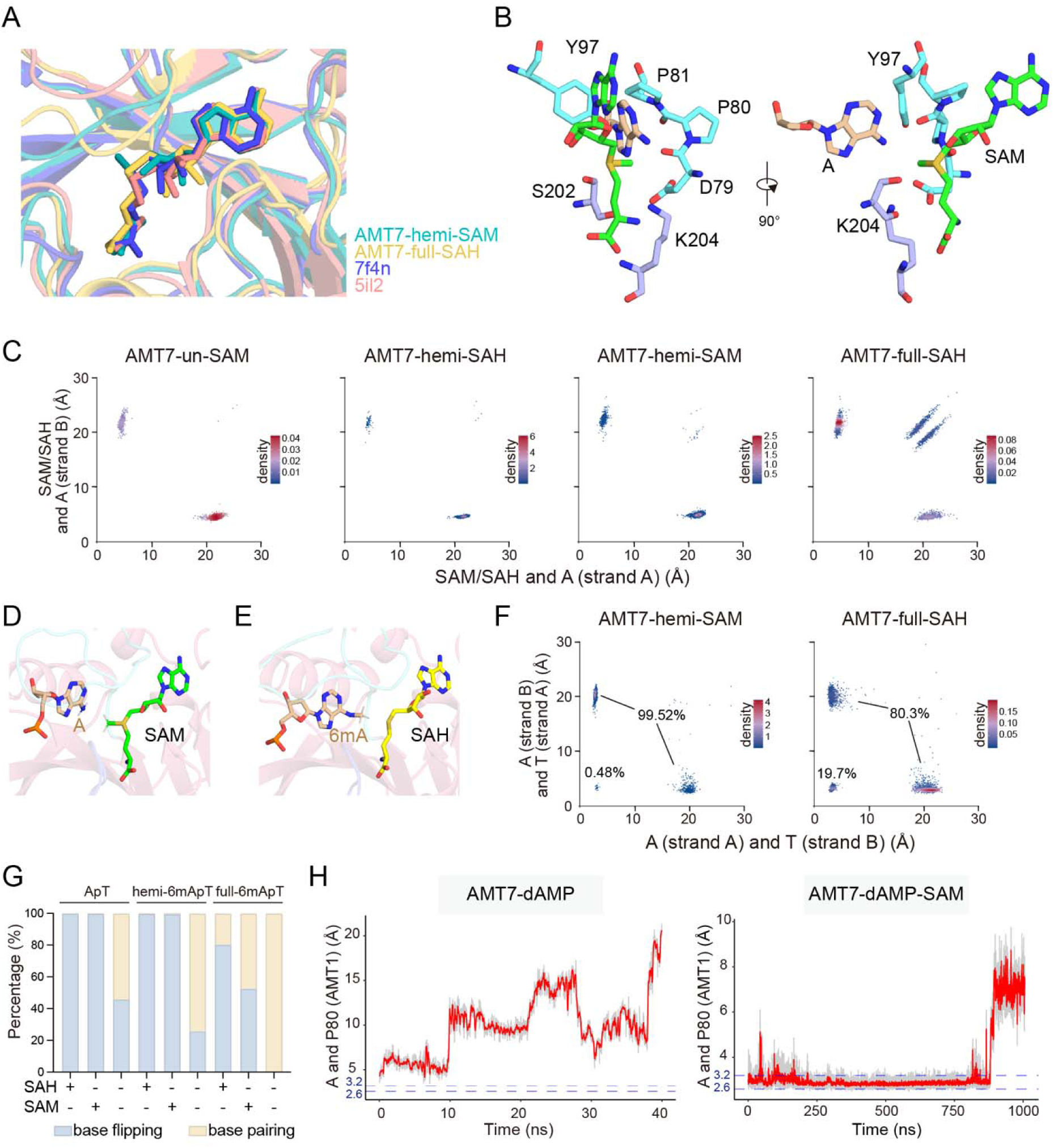
Structure-function relationship in the AMT1 active site. A. Structural overlay of SAM/SAH in the cofactor-binding pocket. The AMT1 ternary holo-complex (AMT7-hemi-SAM and AMT7-full-SAH, MD models) and the cofactor-containing AMT1/METTL3-METTL14 apo-complex (7f4n and 5il2, cryo-EM structures)^24, 29^ are aligned by AMT1/METTL3. B. Critical residues in the aperture separating the cofactor and the extrahelical adenine in the AMT1 ternary complex (AMT7-hemi-SAM, MD model). Residues from the *Major Groove Insertion Loop*: saxe blue; the *Regulation Loop*: lavender. C. Conformational analysis tracking base flipping into the active site. The AMT7 ternary complexes corresponding to the ES/EP state for *de novo* and maintenance methylation. Scatterplot: distance between cofactor (S_δ_) and A/6mA (N^6^) on DNA strand A/B (x/y-axis, Å); colored as per local density of the AF3 models. D. Active site of the ES complex for maintenance methylation (AMT7-hemi-SAM). E. Active site of the EP complex for maintenance methylation (AMT7-full-SAH). F. Conformational analysis tracking base flipping. The ES (AMT7-hemi-SAM) and EP (AMT7-full-SAH) complexes for maintenance methylation. Scatterplot: distance between A/6mA (N^6^) on DNA strand A/B and thymine (O^4^) on DNA strand B/A (x/y-axis, Å); colored as per local density of the AF3 models. G. Proportion of AF3 models in base flipping/pairing for the AMT7 holo-complex. H. Representative MD trajectories for AMT7-dAMP and AMT7-dAMP-SAM, tracking the distance between the adenine (N^6^) and AMT1 P80 (O). The gray traces represent raw frame-by-frame values; the red traces represent the 20-frame moving average. The blue dashed lines mark the 2.2-2.6 Å interval in which the dAMP remains properly positioned in the adenine-binding pocket.

In the AMT1 ternary complex, the extrahelical adenine and SAM are juxtaposed, separated only by a narrow aperture comprising the main/side chains of the AMT1 *Major Groove Insertion Loop* and *Regulation Loop* (Figure 3B, Figure S6D). Key residues within the aperture, including D79 (D395), P80 (P396), P81 (P397), Y97 (Y406), S202 (S511), and K204 (K513), are conserved in METTL3 (Figure 3B, Figure S4C). This allows both the substrate and cofactor to be precisely positioned and orientated, restricting access between them to the exocyclic amino group of the adenine and the sulfonium group of SAM. In most AF3 models of the AMT6/AMT7 ternary complexes (corresponding to the ES and EP states for both *de novo* and maintenance methylation), the distance between the extrahelical adenine and SAH/SAM is less than 5 Å; for AF3 models with high ipTM scores (>0.9), this distance fluctuates around 4.8 Å (Figure 3C, Figure S6E). These values are similar to the equivalent distance in the DNA adenine MTase M.TaqI (4.7 Å)^18^ and the cytosine MTase M.HhaI (5.0 Å)^17^.

Quantum mechanics and molecular mechanics (QM/MM) simulations of the METTL3-METTL14 active site indicate a classical S_N_2 nucleophilic mechanism for methyl transfer^44^. To validate whether the same catalytic mechanism applies here, we used MD simulations to scrutinize the AMT1 active site. We examined the ES complex for maintenance methylation (AMT6/AMT7-hemi-SAM), in which the unmodified adenine is flipped into the active site and juxtaposed with SAM (Figure 3D, Figure S6F). We also examined the EP complex (AMT6/AMT7-full-SAH), in which one of the 6mA is flipped into the active site and juxtaposed with SAH (Figure 3E, Figure S6F). For the EP complex, we focused on the N^6^-protonated extrahelical 6mA, a key intermediate featuring the sp^3^ N^6^, attached to two hydrogen atoms and a N^6^-methyl (Figure 3E, Figure S6F), in accordance with the METTL3-METTL14 QM/MM analysis^44^. Extensive MD simulations of the ES and EP complexes revealed features consistent with an S_N_2 in-line attack mechanism: ***1)*** the transferring methyl group almost perfectly aligns on the axis between the extrahelical A/6mA and the cofactor SAM/SAH, and ***2)*** the distance between the substrate/product and SAM/SAH is also optimal (Figure 3D, E, Figure S6G, H). Similar to the METTL3-METTL14 active site^44^, the AMT1 P80 (METTL3 P395) main chain oxygen is stably positioned next to the extrahelical adenine N^6^ for hydrogen bond formation; concurrently, the D79 (D396) side chain carboxyl group is positioned parallel to the methyl transfer path, neutralizing the positive charge (Figure 3B, Figure S6F). Thus, our structural modeling of the AMT1 ternary complex provides the molecular basis for substrate and cofactor binding, illuminating their optimal placement and alignment for methyl transfer.

### Cooperative adenine and cofactor binding

For AF3 models of the AMT1 holo-complex, we tracked the distance between the A:T pairs in the ApT duplex. Small values (<5 Å) correspond to base pairing, while large values (>15 Å) correspond to base flipping (Figure 3F). For the ES complex, single base flipping was predominantly (99.5%) (Figure 3F, G). Indeed, single base flipping was in the majority across the AMT1 ternary complex, especially among high-scoring AF3 models (>99% at ipTM≥0.9), regardless of their DNA methylation state (full/hemi/un), cofactor (SAM/SAH), or alternative subunit (AMT6/AMT7) (Figure 3G, Supplemental Table S2). However, a substantial drop in single base flipping, accompanied by a corresponding jump in double base pairing, was observed in the cofactor-free binary complex (alongside a substantial drop in ipTM scores). This suggests cooperativity between base flipping and cofactor binding (Figure 3G, Supplemental Table S2).

This cooperativity can be at least partially attributed to the juxtaposition of the adenine-binding and cofactor-binding pockets within AMT1. In the AMT1 ternary complex, several residues at the aperture separating the two pockets—including AMT1 D79, P81, Y97, and K204—simultaneously engage the extrahelical adenine and SAM (Figure 3B). In the absence of the cofactor, some contacts are lost, including the favorable edge-to-face aromatic interaction between AMT1 Y97 and the extrahelical adenine (Figure S7A). We tracked the distance between the extrahelical adenine and AMT1 P80 in the adenine-binding pocket (Figure S7B, C). In almost all AF3 models of the AMT1 ternary complex, the distance was under 5 Å, consistent with close contact and stable binding (Figure S7C). However, the distance was significantly longer and more diffusely distributed in the AF3 models of the AMT1 binary complex, consistent with reduced stability for the extrahelical adenine within its binding pocket (Figure S7C).

We also investigated this cooperativity with extensive MD simulations. For both the AMT1 binary and ternary complexes (AMT7-hemi and AMT7-hemi-SAM), significant conformational changes in the extrahelical adenine were not observed in our MD runs (1 μs) (Figure S7D), indicating slow kinetics for base flipping. This is due to the high binding affinity of the dsDNA^6^. To accelerate the extrahelical adenine dissociation, we used dAMP as a substitute for dsDNA, analogous to a previous MD study of METTL3-METTL14 binding kinetics^44^. Specifically, we replaced the single nucleotide containing the extrahelical adenine with dAMP and removed the rest of the dsDNA (Figure S7E). MD simulations starting from these new structures (AMT7-dAMP and AMT7-dAMP-SAM) revealed substantial differences in their binding behaviors, which we quantified by tracking the distance between the extrahelical adenine and AMT1 P80 in their trajectories (Figure 3H, Figure S7F). In the ternary complex (AMT7-dAMP-SAM), this distance remained at small values (<5 Å) over extended periods, occasionally interrupted by moderate fluctuations (Figure 3H, Figure S7F), supporting deep and stable insertion of the extrahelical adenine into its binding pocket. By contrast, in the binary complex (AMT7-dAMP), this distance started at a higher value and fluctuated erratically (Figure 3H, Figure S7F). It occasionally decreased to ∼5 Å, but only remained there for short intervals (Figure S7F). More often, it increased to >20 Å, signifying adenine dissociation (Figure 3H, Figure S7F). Therefore, without the cofactor, adenine placement in the binary complex is destabilized. Intriguingly, the extrahelical adenine exited the binding pocket via the major groove, facilitated by the hydrophobic core within the AMT1 *Major Groove Insertion Loop* (Figure S7G). Based on both AF3 modeling and MD simulations, we conclude that the extrahelical adenine and the cofactor bind cooperatively within the AMT1 complex.

### Cooperative DNA binding by AMT1 and AMTP1

Both AMTP1 and AMTP2 contain an N-terminal homeobox-like domain (HD) implicated in DNA binding^45^. In our AF3 models, the core of the AMTP1 DBD is superimposable to the solved structure of the AMTP2 HD (Figure S8B), which is exapted to stabilize the interaction between AMT1 and AMTP1^28, 29^. The AMTP1 DBD structure is further supported by XL-MS data (Figure S8C)^29^. The AMTP1 DBD features a slightly concave interface with the dsDNA (Figure 4A, B). DNA binding is facilitated by multiple basic residues at this interface, some of which are conserved across AMTP1 homologues (K44, R68*, K101*, K114, R117, and K118; *: conserved) (Figure 4B, Supplemental File S3). Furthermore, the DNA-recognition helix α3 and its C-terminal loop insert into the major groove (mediated by Y63, M64, Q67*, and Q73*) (Figure 4C, Figure S8D).

**Figure 4.**
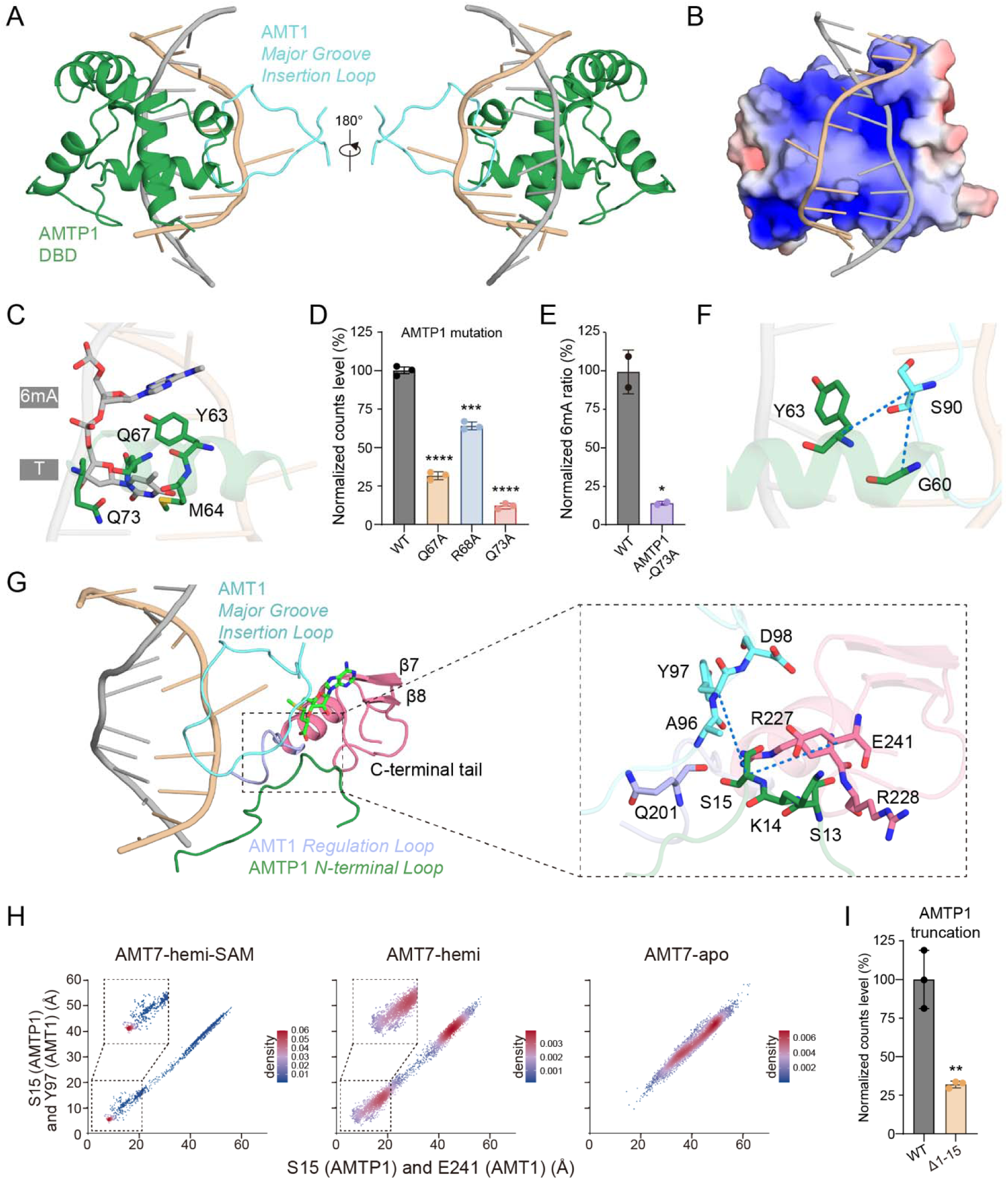
Cooperative DNA binding by AMT1 and AMTP1. A. Opposite views of cooperative binding by the AMT1 *Major Groove Insertion Loop* and the AMTP1 DBD. MD model of the AMT1 ternary complex (AMT7-hemi-SAM), base-flipping mode. Left/right: DNA major groove facing outward/inward. B. Electrostatic surface representation of the AMTP1 DBD. Blue: positive. Red: negative. DNA major groove facing inward. C. Interactions between the AMTP1 DBD and the non-target strand. Ribbon model: the AMTP1 α3-helix and its C-terminal loop, the dsDNA; stick model: critical residues as well as the non-target 6mApT. D. In vitro MTase activity of the reconstituted AMT7 complex with the indicated AMTP1 mutations. The WT complex was included as the positive control. Data are shown as mean±SD. Two-tailed Student’s t-test: \*\*\**P*< 0.001, \*\*\*\**P*< 0.0001. E. Mass spectrometry quantification of the 6mA-to-A ratio (6mA/A, %) in *Tetrahymena* WT and AMTP1 Q73A mutant. Two-tailed Student’s t-test: \**P*< 0.05. F. Interactions between the AMT1 *Major Groove Insertion Loop* and the AMTP1 DBD α3-helix. The distances between AMT1 S90 (C_α_) and AMTP1 G60/Y63 (C_α_) are marked by blue dashed lines. G. The AMTP1 *N-terminal Loop* docking at the AMT1 active site. Expanded view: critical residues at the interface. The distances between AMTP1 S15 (C_α_) and AMT1 E241/Y97 (C_α_) are marked by blue dashed lines. H. Conformational analysis tracking the AMTP1 *N-terminal Loop* docking at the AMT1 active site. AF3 models for the AMT7 ternary holo-complex (AMT7-hemi-SAM), binary holo-complex (AMT7-hemi), and apo-complex (AMT7). Scatterplot: distance between AMTP1 S15 (C_α_) and AMT1 E241/Y97 (C_α_, x/y-axis, Å); colored as per local density of AF3 models. I. In vitro MTase activity of the reconstituted AMT7 complex with the AMTP1 N-terminal truncation (Δ1-15). The WT complex was included as the positive control. Two-tailed Student’s t-test: \*\**P*< 0.01.

Underscoring the critical role of AMTP1, the AMT1-AMT7-AMTP2 sub-complex lacks the high DNA-binding affinity and MTase activity characteristic of the intact AMT1 complex^14, 28^. Furthermore, in the structure of the AMT1 complex bound to a DNA mimic (the Overcoming Classical Restriction [OCR] protein from bacteriophage T7), the AMTP1 DBD is positioned in close proximity to the OCR^30^. In line with these findings, the reconstituted AMT1 complex incorporating AMTP1 mutations, including Q67A, R68A, and Q73A, showed diminished MTase activities (Figure 4D). *Tetrahymena* Q73A mutant also displayed diminished endogenous 6mA levels (Figure 4E). These results support the functional importance of the AMTP1 DBD.

To elucidate the structural details of AMTP1 DNA binding, we examined the MD trajectories of the AMT1 ternary complex. The AMTP1 DBD co-occupies the major groove with the AMT1 *Major Groove Insertion Loop* (Figure 4A). In this cooperative binding mode, the insertion of the AMTP1 α3 helix into the major groove is shallow, and its direct interaction with the target strand ApT is blocked by AMT1 (Figure 4A). Nonetheless, AMTP1 α3 and its C-terminal loop (Y63, Q67, and Q73) can still contact the non-target strand ApT (Figure 4A, C, Figure S8D). Using AMTP1 Y63 as an anchor, we tracked its distance to A/6mA (Figure S8E). For most AF3 models of the AMT1 ternary complex in the base flipping mode, the distance between AMTP1 Y63 and the non-target adenine fluctuated around 10 Å, consistent with shallow insertion of the AMTP1 DBD into the major groove (Figure S8E). In their corresponding MD trajectories, this distance also converged to similar values (Figure S8F).

In the cooperative binding mode, extensive interactions occur between the tip of the AMT1 *Major Groove Insertion Loop* (89-93: PSRGV) and AMTP1 α3 (58-63: SIGQIY) (Figure 4F)—two regions that are mostly conserved (Supplementary File S1 and S3). The tip structure is stabilized by the overlying AMTP1 α3 as well as the underlying DNA (Figure 4A, F). We tracked the distances between AMT1 S90 on one side, and AMTP1 G60 and Y63 on the other (Figure S8G). For most AF3 models of the AMT1 ternary complex in the base flipping mode, both distances were small and narrowly distributed, supporting close contact between the AMT1 *Major Groove Insertion Loop* and the AMTP1 DBD (Figure S8G). Similar findings emerged from their MD trajectories (Figure S8H). Strikingly, these distances increased and were more diffusely distributed across AF3 models of the AMT1 ternary complex in the base pairing mode, suggesting reduced cooperativity (Figure S8G). This cooperative binding all but disappeared in AF3 models of the cofactor-free binary complex (Figure S8I). These data indicate that cooperative binding by AMT1 and AMTP1—much like base flipping—is strongly promoted by cofactor binding.

### AMTP1 N-terminal Loop

The AMTP1 *N-terminal Loop* contributes to these cooperative binding events within the AMT1 ternary complex (Figure 4G). It contacts the AMT1 *Major Groove Insertion Loop* and the *Regulation Loop*, which are shared by the adenine-binding and cofactor-binding pockets (Figure 3B). It also contacts AMT1 at the β7-β8 connecting loop and the C-terminus, which are part of the cofactor-binding pocket (Figure 3A). Important contacts include AMTP1 S13, K14, and S15 on one side, and AMT1 A96, Y97, D98, Q201, R227, R228, and E241 on the other (Figure 4G). We tracked the distances between S15 in the AMTP1 *N-terminal Loop* and Y97 and E241 in the AMT1 active site (Figure 4H, Figure S9A, B). For most AF3 models of the AMT1 ternary complex in the base flipping mode, both distances fluctuated narrowly around two small values, consistent with the AMTP1 *N-terminal Loop* docking at the AMT1 active site (Figure 4H). These distances were also stable in their corresponding MD trajectories (Figure S9C). However, they increased dramatically and variably for most AF3 models of the AMT1 ternary complex in the base pairing mode (Figure 4H, Figure S9A). This shift was all but complete in the cofactor-free holo-complex and apo-complex (Figure 4H, Figure S9A, B). When the AMTP1 N-terminal tail is not docked at the AMT1 active site, it loses its well-defined local structure as well as its global arrangement (Figure S9D). These observations directly link adenine and cofactor binding to the disorder-to-order transition in the AMTP1 N-terminal region and its docking at the AMT1 active site.

The AMTP1 *N-terminal Loop* is connected to the AMTP1 DBD via a conserved short linker (23-25: PGW) (Figure S9E). As the AMTP1 *N-terminal Loop* docks at the AMT1 active site, it brings this linker region close to the target strand, enhancing DNA binding by the AMTP1 DBD as well as its interaction with the tip of the AMT1 *Major Groove Insertion Loop* (Figure S9E). Cooperative binding by AMT1 and AMTP1 is thereby coordinated with cooperative binding of adenine and cofactor. The AMTP1 N-terminal region is mostly conserved (Supplemental File S3). Its functional importance was confirmed by the diminished MTase activity in the reconstituted AMT7 complex featuring a truncated AMTP1 (Δ1-15) (Figure 4I).

### Independent ApT recognition by AMTP1

We also identified AF3 models in which the AMTP1 DBD inserted deeper into the major groove at the ApT site, revealed by significant decreases in the distance between AMTP1 Y63 and A/6mA relative to the cooperative binding mode (Figure S10A). In this AMTP1-dominated binding mode, the distal portion of the AMT1 *Major Groove Insertion Loop* is displaced from the major groove and separated from the AMTP1 DBD (Figure 5A). The former was evident from increased distances between AMT1 R91 and A/6mA, while the latter was evident from increased distances between AMT1 S90 and AMTP1 G60/Y63 (Figure S10B, C). The AMT1 *Major Groove Insertion Loop* is structurally heterogeneous after losing contact with the major groove as well as the AMTP1 DBD (Figure S10D).

**Figure 5.**
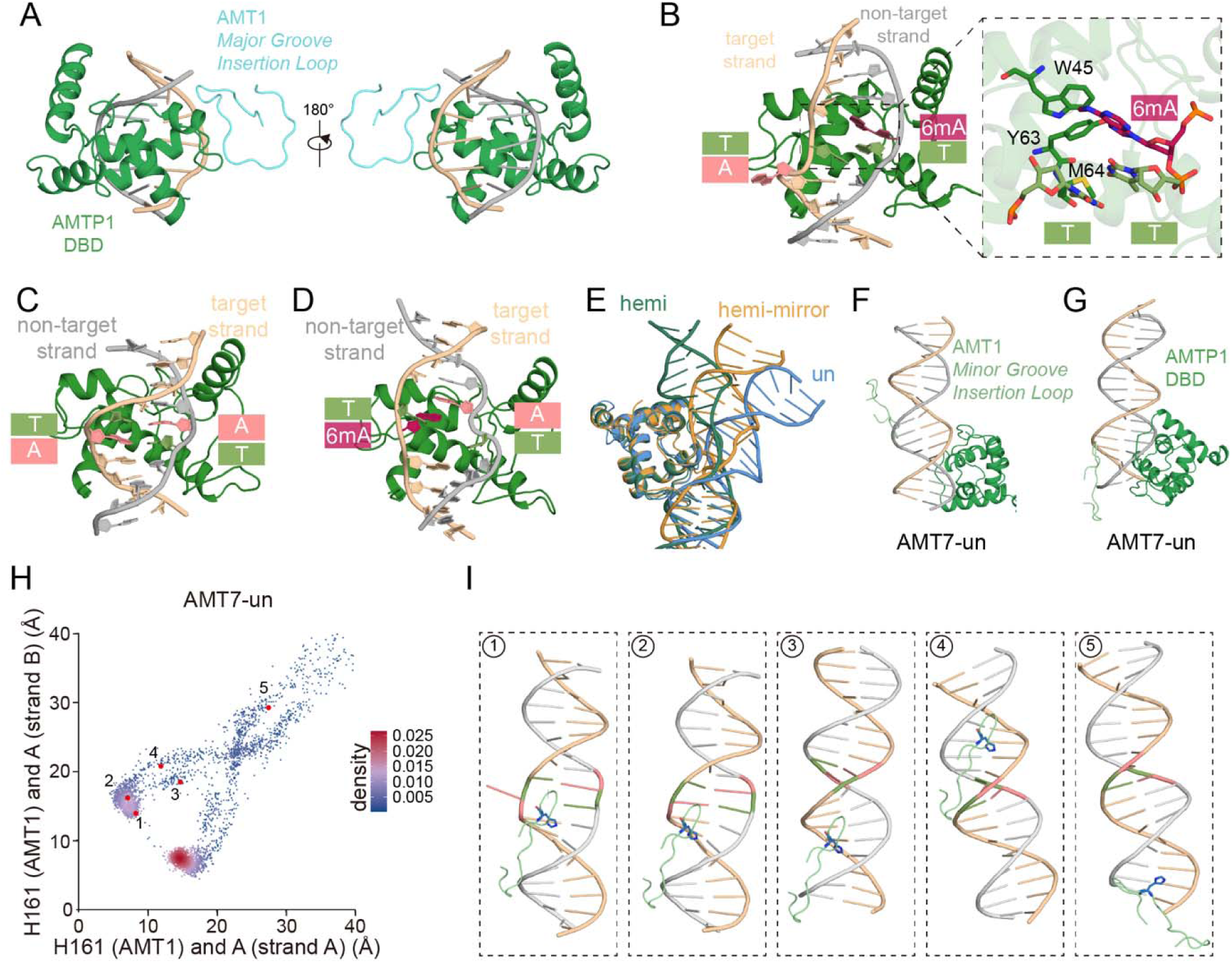
AMTP1-dominated DNA binding and offsite binding. A. Opposite views of the AMTP1-dominated binding of the dsDNA. A representative AF3 model for the AMT1 binary complex (AMT7-hemi) in the base pairing mode. Left/right: DNA major groove facing outward/inward. Note that the AMT1 *Major Groove Insertion Loop* shifts away from the AMTP1 DBD and the ApT site. B. A representative MD model for the AMTP1 DBD in association with the hemi-methylated dsDNA. Note the disruption of the ApT duplex and the interactions with the centrally positioned hydrophobic residues. 6mA is on the non-target strand, while the adenine, flipped out of the dsDNA helix, is on the target strand. C. A representative MD model for the AMTP1 DBD in association with the un-methylated dsDNA. Note the normal base pairing of the ApT duplex. D. A representative MD model for the AMTP1 DBD in association with the symmetry-related hemi-methylated dsDNA. 6mA is on the strand, while the adenine is on the non-target strand. Note the absence of base flipping on the target strand. E. Superimposition of the MD models for the AMTP1 DBD in association with the differentially methylated dsDNA (B-D), aligned by the AMTP1 DBD. F. Offsite binding by the AMTP1 DBD (forest green) and onsite binding by the AMT1 *Minor Groove Insertion Loop*. G. Offsite binding by both the AMTP1 DBD (forest green) and the AMT1 *Minor Groove Insertion Loop*. H. Conformational analysis tracking DNA binding by the AMT1 *Minor Groove Insertion Loop* in the AMT1 binary complex (AMT7-un). Scatterplot: distance between AMT1 H161 (C_α_) and A/6mA (C3’) on DNA strand A/B (x/y-axis, Å); colored as per local density of AF3 models. Important conformations: 1) onsite binding in the base flipping mode, 2) onsite binding in the base pairing mode, 3) offsite binding in the major groove, 4) offsite binding in the minor groove, and 5) faraway offsite binding. I. Correponding AF3 models in the AMT1 binary complex (AMT7-un). Only the AMT1 *Minor Groove Insertion Loop* and the dsDNA are shown. Strand A: silicon. Strand B: silver. Adenine: salmon. Thymine: smudge. The AMT1 *Minor Groove Insertion Loop*: palegreen. AMT1 H161: stick model.

This AMTP1-dominated binding mode was virtually absent from the AMT1 ternary complex, but emerged in the base pairing mode of the AMT1 binary complex (Figure S8E, S10A). We focused on one such high-scoring AF3 model (AMT7-hemi), in which the 6mA was placed on the non-target strand while the unmodified A was placed on the target strand, corresponding to a key intermediate in the reaction pathway (Figure 5A). To better understand its DNA binding behavior, we isolated the AMTP1 DBD alongside its associated dsDNA from this model and performed MD simulations. Strikingly, we observed consistent dsDNA deformation following extended MD runs, while the AMTP1 DBD remained largely unchanged (Figure 5B, Figure S10E). The substantial structural differences from the starting AF3 model arise as MD simulations explore the local energy landscape, thereby allowing us to evaluate structural stability and dynamics. Here, the dsDNA is stretched and unwound around the ApT duplex (Figure S10E). Base pairing within the ApT duplex is disrupted, with the four bases frequently stacking on top of one another (Figure 5B). The distorted dsDNA bends towards the AMTP1 DBD and makes more contacts, especially at the positively charged edges of its DNA-binding interface (Figure S10E). At the center, a stretch of hydrophobic residues (W45, Y63, M64) align with the N6-methyl of the non-target 6mA, the 5-methyl of the target thymine, and the 5-methyl of the non-target thymine (Figure 5B). Intriguingly, the target adenine—no longer base paired—can readily flip out from the minor groove (Figure 5B, Figure S10E).

We performed control MD simulations starting from essentially the same structure, but with different methylation states of the ApT duplex. First, we altered the non-target 6mA to unmodified adenine (Figure 5C). As a result, normal base pairing is restored, and the dsDNA becomes more compressed and overwound relative to the original hemi-6mApT structure (Figure 5C, Figure S10F). The dsDNA no longer bends towards the AMTP1 DBD, making fewer contacts (Figure S10F). We also swapped the N^6^-methyl group from the non-target adenine to the target adenine (Figure 5D). This leads to an intermediate structure: the ApT duplex is still disrupted, but the dsDNA overall is less deformed, making fewer contacts with the AMTP1 DBD than in the hemi-6mApT structure, but more than in its unmodified counterpart (Figure 5D, E, Figure S10G). Importantly, the 6mA on the target strand, while still unpaired, remains intrahelical (Figure S10G). These MD simulations suggest that the AMTP1 DBD *per se* prefers hemi-6mApT and, upon binding, promotes flipping of the unmodified adenine on the target strand.

### Offsite DNA binding

The AMTP1 DBD is positioned at the major groove and near the ApT site in both the cooperative and AMTP1-dominated binding modes, corresponding to onsite binding. In contrast to onsite binding, the AMTP1 DBD can also be positioned away from ApT, as indicated by increased distances between AMTP1 Y63 and A/6mA, relative to the cooperative binding mode (Figure 5F, Figure S8E). AMTP1 offsite binding includes a continuum of structures, in which the AMTP1 DBD progressively shifts away from ApT, initially moving along the major groove but ultimately reaching the minor groove (Figure 5F). This is also accompanied by separation of the AMTP1 DBD and the AMT1 *Major Groove Insertion Loop* (Figure 5F). For the AMT1 ternary complex, AMTP1 offsite binding represents only a small minority in the base flipping mode, but substantially more in the base pairing mode; it is common in the AMT1 binary complex (Figure S11A).

AMTP1 offsite binding generally occurs when the AMT1 *Minor Groove Insertion Loop* still engages the ApT site (Figure 5F). As part of the rigid heterodimerization interface between the MT-A70 domains, the AMT1 *Minor Groove Insertion Loop* is the most stable protein-DNA interface in the AMT1 holo-complex, in strong contrast to dynamic DNA binding by the AMT1 *Major Groove Insertion Loop* and the AMTP1 DBD. Using H161 in the AMT1 *Minor Groove Insertion Loop* as an anchor point, we tracked its distances to A/6mA (Figure S11B, C). AMT1 onsite binding was found in virtually all AF3 models of the AMT1 ternary complex, as well as in their corresponding MD simulations (Figure S11B, C). These results support that AMT1 initially engages the ApT site by its *Minor Groove Insertion Loop*; it subsequently pivots around the extrahelical adenine and makes additional contacts with its *Major Groove Insertion Loop*, in coordination with the AMTP1 DBD. AMT1 offsite binding is readily observed only in the AMT1 binary complex, especially with unmodified ApT: AMT1 mainly tracks along the minor groove and distributes rather evenly along the dsDNA (Figure 5H, I, Figure S11B, D, E). This suggests a scanning mechanism for processive methylation, whereby translocation of the AMT1 complex along the dsDNA enables an efficient one-dimensional search.

### Interactions securing the closed form

In the AMT1 ternary complex (AMT7-hemi-SAM), the AMTP1 DBD wraps around the non-target strand of the dsDNA and contacts the AMT7 MT-A70 domain, forming an interface absent in the apo-complex, hereafter referred to as the clasp (Figure 6A, Figure S12A). Most contacts in the clasp are between the N-terminal portion of the AMTP1 *Clasp Loop* and the C-terminal portion of the AMT7 *Multiple Interaction Loop*; in particular, AMT7 R282 side chain can reach across the interface and form hydrogen bonds with AMTP1 D110 side chain (Figure 6A). Importantly, the C-terminal portion of the AMT7 *Multiple Interaction Loop* (ELRHQR; AMT6: 202-207; AMT7: 280-285)— absolutely conserved between AMT6 and AMT7 (File S2)—simultaneously engages ***1)*** the AMT1 *Minor Groove Insertion Loop* (mediated by AMT7 L281, H283 and AMT1 F163, Y164), ***2)*** the AMT7 *Hydrophobic Loop* (mediated by AMT7 L317, L318, P319, K320 and AMT7 E280, L281), and ***3)*** the AMTP1 *Clasp Loop* (Figure 2C, Figure 6A, Figure S12A). The AMTP1 *Clasp Loop* (107-111: NTGDN) is also mostly conserved (File S3). Similar interactions are present in other AMT6/AMT7 complexes (Figure S12B), supporting the clasp connecting AMTP1 and AMT6/AMT7 as a general feature securing the closed form AMT1 complex.

**Figure 6.**
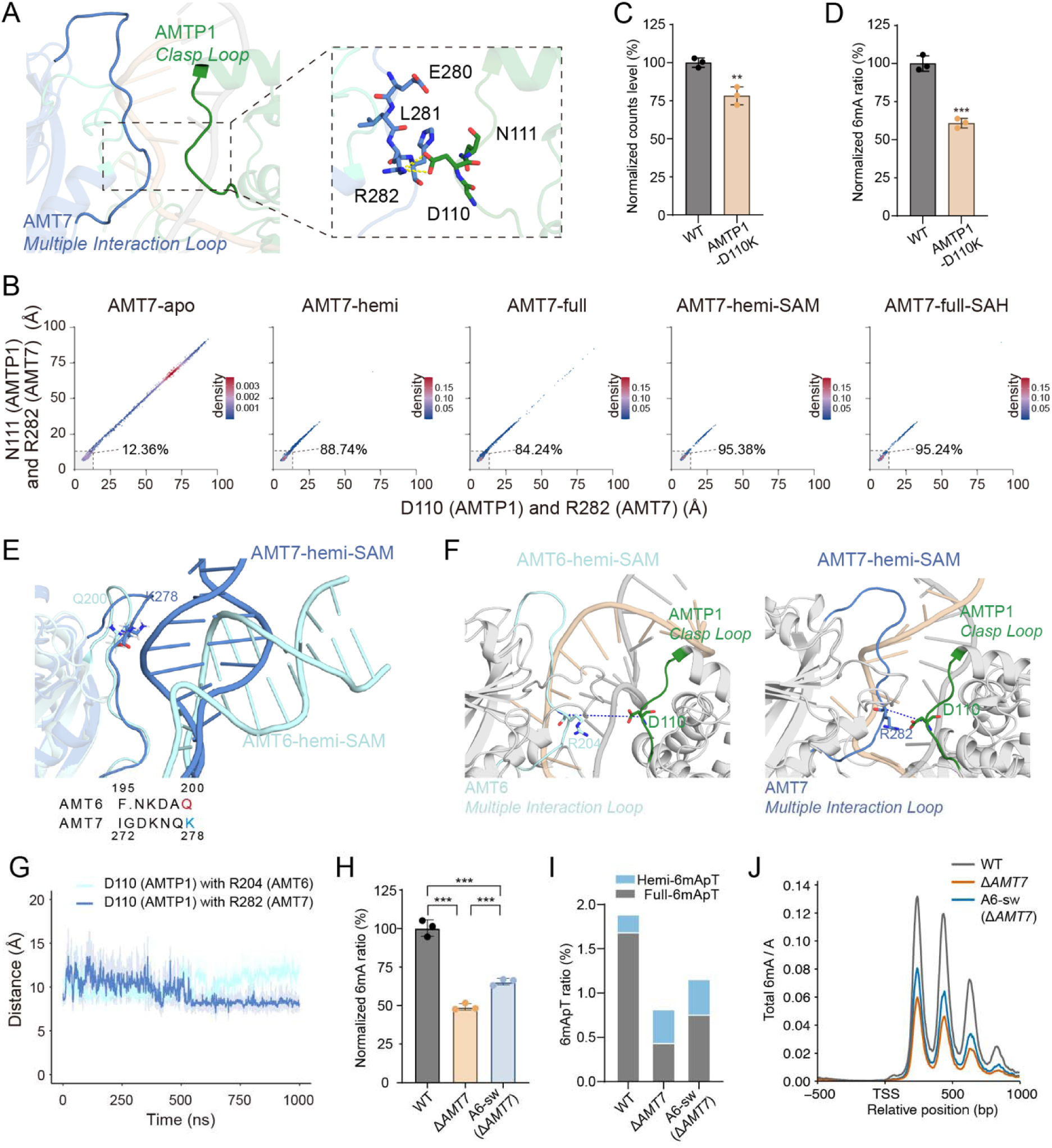
Interactions securing the closed form AMT1 complex. A. The clasp securing the closed form AMT1 ternary complex (AMT7-hemi-SAM). AMTP1 *Clasp Loop* and AMT7 *Multiple Interaction Loop* are highlighted. Expanded view: residues critical for clasp formation. Yellow dashed line: hydrogen bond. B. Conformational analysis tracking clasp formation in the AMT7 complex (AMT7, AMT7-hemi, AMT7-full, AMT7-hemi-SAM, AMT7-full-SAH). Scatterplot: distance between AMT7 R282 (C_α_) and AMTP1 D110/N111 (C_α_; x/y-axis, Å); colored as per local density of AF3 models. Percentage of AF3 models consistent with clasp formation is indicated; threshold is set at 12.5 Å for both distances. C. In vitro MTase activity of the AMT7 complex with the AMTP1 D110 K mutation. Two-tailed Student’s t-test. **p < 0.01. D. MS quantification of the 6mA-to-A ratio (6mA/A, %) in WT and the AMTP1 D110K mutant. Two-tailed Student’s t-test. ***p < 0.001. E. Differential interactions between the AMT6/AMT7 *Multiple Interaction Loop* and the dsDNA. MD models for the AMT1 ternary complex (AMT6/AMT7-hemi-SAM). AMT6 Q200 and AMT7 K278 are shown in the stick model. Bottom: the AMT6/AMT7 sequences for the variance region. F. Differential interactions between the AMT6/AMT7 *Multiple Interaction Loop* and the AMTP1 DBD. MD models for the AMT1 ternary complex (AMT6/AMT7-hemi-SAM). AMTP1 D110, AMT6 R204, and AMT7 R282 are shown in the stick model. G. MD trajectories for the AMT1 ternary complex (AMT6/AMT7-hemi-SAM) tracking the distance between AMTP1 D110 (C_α_) and AMT6 R204/AMT7 R282 (C_α_). The light-colored traces represent raw frame-by-frame values; the dark-colored traces represent the 20-frame moving average. H. MS quantification of the 6mA-to-A ratio (6mA/A, %) in WT, Δ*AMT7*, and AMT6 swap (A6-sw) cells. Note that *AMT7* is deleted from both mutants. Two-tailed Student’s t-test. ***p < 0.001. 6mA levels in A6-sw was still substantially lower than WT (containing WT AMT6 and AMT7, both expressed from their endogenous loci), which is attributable to other sequence variations between AMT6 and AMT7, as well as differences in their expression levels. I. Quantification of 6mApT in WT, Δ*AMT7* and AMT6 swap cells by PacBio sequencing. Full- and hemi-6mApT are represented separately. J. 6mA distribution around the transcription start site (TSS) of Pol II-transcribed genes in WT, Δ*AMT7*, and AMT6 swap cells. 6mA penetrance is tallied across 6mA+ genes, as previously reported^6^.

The dynamics of the clasp are dictated by the reaction cycle: the clasp can stabilize DNA binding and promote catalysis, while its unhooking is required for loading and unloading of the dsDNA. For most AF3 models of the AMT1 ternary complex, the distances between the AMTP1 *Clasp Loop* and the AMT6/AMT7 *Multiple Interaction Loop* fluctuated around values optimal for interactions, supporting clasp formation (Figure 6B, Figure S12C). Substantial increases in the distances revealed conformations incompatible with the clasp formation and the closed form: such conformations increased in the AMT1 binary holo-complex; they became predominant in the AMT1 apo-complex (Figure 6B, Figure S12C). The functional importance of the clasp was confirmed by the reduced *in vitro* MTase activity of the reconstituted AMT1 complex incorporating the AMTP1 D110K mutation (Figure 6C). The corresponding *Tetrahymena* mutant also showed diminished endogenous 6mA levels (Figure 6D).

### Structural basis for differential activities of the AMT6 and AMT7 complexes

In *Tetrahymena*, the AMT7 complex, with higher DNA-binding and catalytic activities, is important for processive methylation, while the AMT6 complex is much less active and processive^6^. We next investigated the structural basis for their differential activities. Most regions involved in DNA binding are conserved across AMT6 and AMT7, except for the N-terminal portion of the *Multiple Interaction Loop*, which is represented by divergent sequences in AMT6 (195-200: F·NKDA**<u>Q</u>**) and AMT7 (272-278: IGDKNQ**<u>K</u>**) (Figure 6E). Critically, the positively charged K278 in AMT7 is replaced by the neutral Q200 in AMT6. Indeed, MD simulations revealed in the AMT6 ternary complex a severe bending of the dsDNA at the ApT site, away from AMT6 but toward the AMTP1 DBD; similar bending was not observed in its AMT7 counterpart (Figure S12D). Consequently, in the AMT7 complex, this variance region inserts into the major groove and makes extensive DNA interactions, including contacts between AMT7 K278 and the phosphate backbone (Figure 6E). By contrast, the AMT6 counterpart is much further removed from the dsDNA, precluding contacts between AMT6 Q200 and the phosphate backbone (Figure 6E).

The differential placement of the variance region is propagated to the conserved C-terminal portion of the AMT6/AMT7 *Multiple Interaction Loop* and across the clasp, leading to the AMTP1 DBD being placed closer to AMT7 than AMT6 within their respective ternary complexes (Figure 6F, Figure S12E). MD trajectories showed that the distance between the AMTP1 *Clasp Loop* and the *Multiple Interaction Loop* converged to much smaller values in the AMT7 ternary complex than its AMT6 counterpart (Figure 6G). AF3 modeling revealed a warped form, in which the AMTP1 *Clasp Loop* is shifted relative to the *Multiple Interaction Loop*, prepositioning the AMTP1 DBD for offsite binding and the AMT1 complex for transition into the open form (Figure S12F). The transition from the closed from, through the warped form, to the open form was more obviously observed in the AMT6 complex than the AMT7 complex (Figure S12G). These analyses strongly suggests that the variance regions of AMT6 and AMT7 contribute to their differential MTase activities by affecting DNA binding and clasp stability.

To validate its functional importance, we swapped the variance region in AMT6 with its AMT7 counterpart and introduced this mutation into Δ*AMT7* cells. We isolated genomic DNA from the mutant and quantified its 6mA levels (Figure 6H, I). MS analysis showed that the mutant (containing the region-swapped AMT6 expressed from the endogenous *AMT6* locus) had significantly elevated total 6mA levels relative to the original Δ*AMT7* strain (containing WT AMT6) (Figure 6H). PacBio sequencing showed that 6mApT levels in the mutant increased by almost 50% compared to Δ*AMT7* (6mApT/ApT: 0.82% vs. 1.16%); furthermore, the global hemi-6mApT levels dropped from 46.6% to 34.7%, while the global full-6mApT jumped from 53.4% to 65.3% (Figure 6I, Figure S12H, I). PacBio sequencing also revealed that in the mutant 6mA levels increased proportionally across the genome, maintaining its specific distribution patterns along the gene body (Figure 6J). These results pinpoint the variance region as a key contributor to the differential activities between the AMT6 and AMT7 complexes.

## Discussion

### Accurate modeling of the AMT1 complex

As this study was under review, cryo-electron microscopy (cryo-EM) structures of the AMT1 holo-complex were published^46^. While our work primarily focuses on the AMT7 complex and theirs on the AMT6 complex, there is a striking similarity in the overall structural organization of this prototypical eukaryotic DNA 6mA-MTase (Figure 7A, Figure S13A). Both studies support the following structural features regarding the ES complex for maintenance methylation: ***1)*** the dsDNA is directionally inserted into a torus formed by the four protein subunits, ***2)*** the ApT duplex is specifically recognized by the AMT1 substrate-binding loops and the AMTP1 DBD, ***3)*** the target adenine is flipped out of the dsDNA helix and into the AMT1 active site, ***4)*** methyl transfer is promoted by the cooperative binding and optimal alignment of the target adenine and the cofactor at the active site (Figure 7A, Figure S13A). While their work used AlphaFold2 to model individual subunits, which were subsequently docked onto cryo-EM maps and refined^46^, we applied AF3 to directly model the entire protein-DNA-cofactor ternary complex. The close match between these two results is a testimony of the accuracy of the state-of-the-art AI-based modeling and its power to dissect biomolecular interactions.

**Figure 7.**
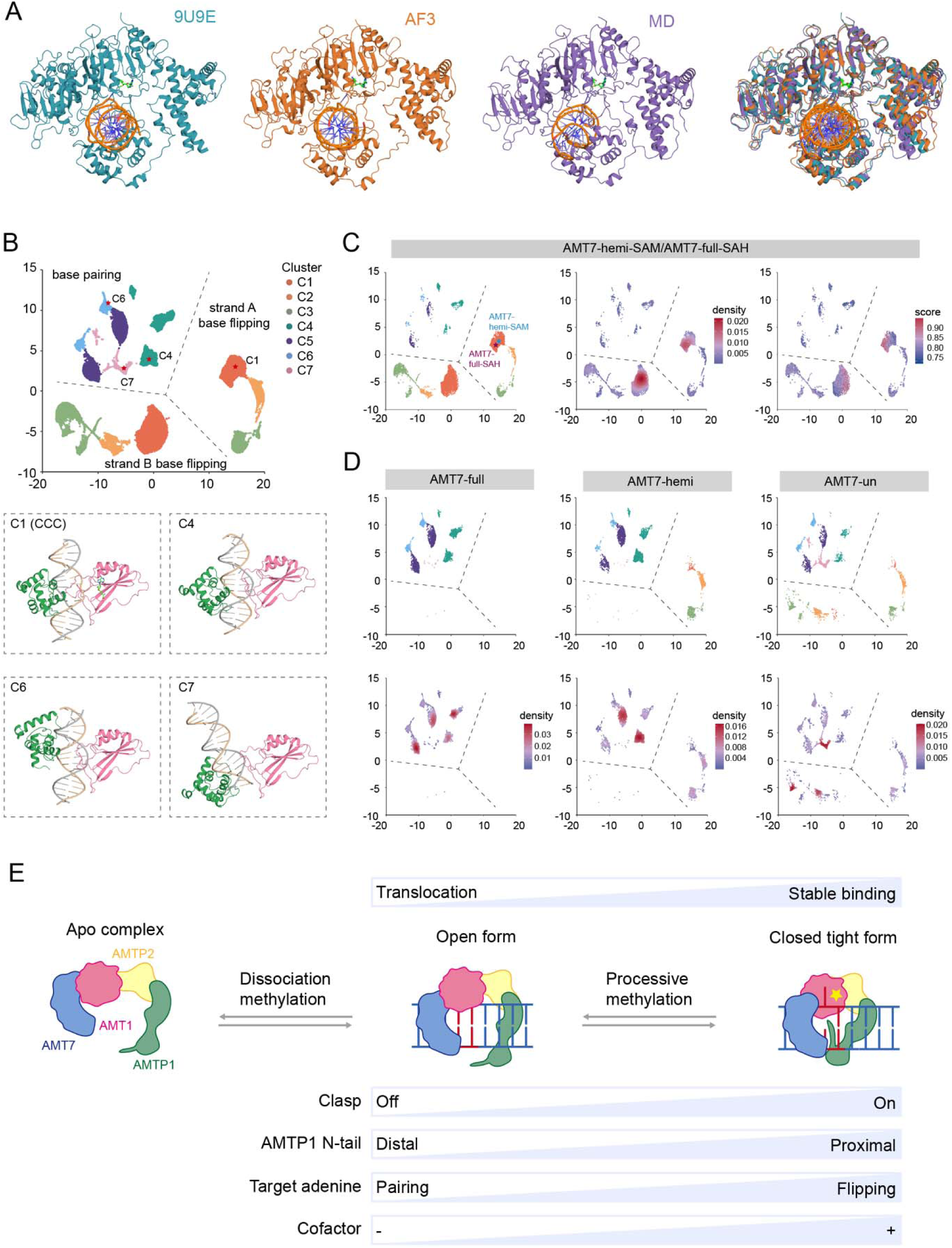
Reaction pathway for DNA adenine methylation catalyzed by the AMT1 complex. A. Side-by-side comparison of the AMT1 ternary complex structural models: the cryo-EM structure (AMT6-hemi-SAM cryo-EM, 9U9E), the AF3 model after initial MD solvation and equilibration (AMT7-hemi-SAM, AF3), and the AF3 model after extended MD simulations (AMT7-hemi-SAM, MD). Overlay of the three structures is on the right. B. UMAP analysis of the AMT7 holo-complex. 7 clusters (C1 to C7), featuring distinct structural characteristics, are demarcated and differentially colored. Demarcations separating the base pairing mode and the base flipping mode are shown; the latter is further divided by selection of the target strand for base flipping (A/B). Below: representative structures in C1 (CCC), C4 (AMTP1-dominated binding), C6 (AMTP1 offsite binding), and C7 (AMT1 offsite binding); only the AMTP1 DBD, AMT1 MT-A70 domain, and dsDNA are shown. C. UMAP analysis of the AMT7 ternary complex (AMT7-hemi-SAM and AMT7-full-SAH). Colored by cluster (left), density (middle), and AF3 score (right). D. UMAP analysis of the AMT7 binary complex (AMT7-full, AMT7-hemi, and AMT7-un). Colored by cluster (top) and density (bottom). E. A simplified reaction pathway for processive maintenance methylation catalyzed by the AMT1 complex. See text for details.

Intriguingly, one feature of the cryo-EM structure is poorly predicted by AF3. In the cryo-EM atomic model, the non-target thymine partially flips out from the minor groove, alongside flipping of the target adenine; by contrast, single flipping of the target adenine is predominant in AF3 models (Figure S13B). Out of the 60,000 AF3 models generated for all combinations of the AMT1 ternary complex (AMT6/AMT7, SAM/SAH, un/hemi/full), less than 0.1% exhibited base flipping on the non-target strand as well as the target strand. Among them, only a handful exhibited flipping of the non-target thymine, which cannot be singled out with confidence *a priori*. Furthermore, in the cryo-EM model, the extrahelical thymine rotates around its glycosidic bond; this shift from the *anti* to the *syn* conformation was completely absent in our models (Figure S13B). The unusual double base-flipping configuration is apparently beyond the limit of AF3 prediction.

### Catalytically competent conformation of the AMT1 complex

While AF3 is powerful at modeling biomolecular complexes, it is essentially agnostic to physical interactions. Complementarily, MD simulations can effectively explore the local energy landscape and generate realistic atomic models. By combining both AF3 modeling and MD simulations, we can home in on critical structures along the reaction pathway and envision the transition between them. A strong consensus regarding the placement of the AMTP1 DBD in the AMT1 ternary complex has emerged (Figure 7A). This region, absent from the AMT1 apo-complex structures^28, 29^, has long been proposed to bind DNA, analogous to the target recognition region (TRD) of prokaryotic adenine MTases such as CcrM and EcoP15I^26, 27^. With the AMTP1 DBD on one side and the heterodimeric MT-A70 domains on the other, a functional clamp is formed. This clamp clutches the dsDNA and induces severe distortions, including widening of the major and minor grooves, underwinding and bending of the DNA backbone, and, ultimately, flipping of the target adenine. Furthermore, the placement of the adenine and the cofactor within the AMT1 active site is rigidly defined and optimally aligned for methyl transfer. We identify this structure as the catalytically competent conformation (CCC).

CCC assembly involves the coordination of the AMT1 *Major Groove Insertion Loop* and the AMTP1 DBD, along with its N-terminal tail. In CCC, the distal portion of the AMT1 *Major Groove Insertion Loop* is sandwiched in between the underlying DNA and the overlying AMTP1 DBD. The AMTP1 DBD, in turn, is anchored by ***1)*** its long C-terminal linker to the AMT1 N-terminal extension module, ***2)*** its short N-terminal linker to the AMTP1 *N-terminal Loop* and the AMT1 active site, and ***3)*** the clasp to the AMT6/AMT7 MT-A70 module. Furthermore, the AMTP1 *N-terminal Loop* fills up the space between the AMT1 active site and the DNA major groove. It functions as a linchpin that limits movement of the AMT1 complex, thereby dictating the transition between stable binding at the target site and facile translocation in search of the next. By coordinating all these interactions for CCC assembly, the transition state for methyl transfer is effectively stabilized. Critically, the active site of the AMT1 complex CCC is structurally conserved in METTL3-METTL14, supporting a unified catalytic mechanism for the MT-A70 family of eukaryotic amino-MTases. Our structural insight can therefore promote the rational design of small-molecule inhibitors for not only the AMT1 complex, but also the METTL3-METTL14 complex—a promising therapeutic target in cancer and leukemia^35, 36^.

### Conformational heterogeneity and dynamics of the AMT1 complex

AMT1-catalyzed DNA methylation is inherently dynamic, as it entails loading and unloading of the DNA substrate and cofactor, as well as flipping of the target adenine in and out of the active site. It involves large-scale and/or long-range conformational changes that is kinetically slow and computationally prohibitory for MD simulations. We argue that the requisite flexibility of the AMT1 complex is reflected by the diverse conformations observed in our AF3 models. In other words, the various protein-protein and protein-DNA interactions underpinning these dynamic behaviors are optimized by evolution, encoded within multiple sequence alignments, and interpreted by the AF3 to output the corresponding heterogenous conformations. Intriguingly, these conformations are readily perturbed by AF3 input settings, particularly ***1)*** the presence or absence of the DNA substrate and cofactor, ***2)*** DNA substrates with different methylation states, and ***3)*** alternative protein subunits. In this study, extensive AF3 sampling across a full array of input settings is exploited to survey the conformational space of the AMT1 complex.

Using Uniform Manifold Approximation and Projection (UMAP)^47^, we provide a comprehensive analysis of 90,000 AF3 models for all combinations of the AMT1 binary and ternary holo-complexes, tracking 8 pairs of distances across the dynamic protein-protein and protein-DNA interfaces (Figure 7B, Figure S14A-C, Supplemental Table S3). 7 clusters (C1 to C7) can be readily identified on the UMAP. Due to the palindromic nature of the dsDNA, each cluster contains two sub-clusters that are separated on the UMAP but are related by symmetry. This is most obvious for the AF3 models in the base flipping mode (C1 to C3), in which selection of the target strand(with extrahelical 6mA/A proximal to the active site) and the non-target strand (with intrahelical 6mA/A distal to the active site) is largely random. Similar relationship also holds in the base pairing mode (C4 to C7).

We first focus on the AMT1 ternary complex, which correspond to the ES and EP states for maintenance methylation (Figure 7C, Figure S14B). The majority of these AF3 models are in the base flipping mode, tightly clustered in C1 (Figure 7C, Figure S14B). The high-scoring AF3 models are also enriched in C1 (Figure 7C, Figure S14B). Indeed, C1 models are closely related to the CCC of the AMT1 complex, highly uniform with regard to the specific structural features tracked by the parameters (Figure 7C). In other words, the CCC, as a typical structure in C1, is robustly predicted by AF3.

While the CCC represents a critical snapshot of DNA adenine methylation catalyzed by the AMT1 complex, even minority classes of conformations provide clues about the reaction pathway. While still in the base flipping mode, the AMTP1 N-terminal tail in C2 and C3 models is increasingly removed from the active site (Figure 7C, Figure S14B). This suggests that the AMTP1 N-terminal tail, properly docked at the active site to form the AMTP1 *N-terminal Loop*, may represent the keystone for CCC assembly. The conformational heterogeneity of the AMT1 *Major Groove Insertion Loop*, especially within the binary complex, is consistent with its role as a dynamic “lid” that controls the flipping of the target adenine (Figure 7D, Figure S14C).

### Reaction pathway for DNA adenine methylation catalyzed by the AMT1 complex

Our structural modeling support that the ApT duplex is recognized not only by the CCC (C1), but also by a distinct conformation of the binary complex in the base pairing mode (C4) (Figure 7B-D, Figure S14A-C). Importantly, the latter provides a plausible scenario for preferential binding of hemi-6mApT by the AMTP1 DBD, independent of AMT1 binding. This specific recognition, likely kinetic in nature, also destabilizes the intrahelical adenine on the target strand, which is poised to flip out from the minor groove. This quick search for the target site is followed by CCC assembly for its verification, which constitutes a double-check mechanism balancing both efficiency and accuracy (Figure 7E).

Processive methylation occurs in an AMT7-dependent manner in *Tetrahymena*^6^. Based on our structural modeling, we propose that the AMT1 complex performs a one-dimensional search for ApT sites by translocating along the dsDNA. This is likely achieved by a gated sliding clamp model, in which the dynamic clasp structure allows the AMT1 complex to alternate between the closed form and the open form (Figure 7E). In the closed form, clasp formation stabilizes the protein-DNA-cofactor ternary complex and promotes catalysis. The clasp is strained in the warped form and disrupted in the open form, loosening the protein-DNA interactions. The clasp opening is generally accompanied by removal of the AMT1 N-terminal tail from the AMT1 active site, leaving more room for the AMT1 complex to slide along the dsDNA. In this context, the AMTP1 and AMT1 offsite binding modes (C6 and C7) represent key intermediates (Figure 7D, E). As processivity is promoted by continuous DNA association, the lower DNA-binding affinity and clasp stability inherent to the AMT6 complex may account for its differential activities relative to the AMT7 complex^6^.

We conclude by integrating a reaction pathway for processive maintenance methylation catalyzed by the AMT1 complex (Figure 7E, Figure S15). First, the AMT1 complex, in its open form, is loaded onto the dsDNA (Figure 7E). In the binary holo-complex, the AMT1 tetramer functions as a gated sliding clamp, transitioning between the loose binding open form and the tight binding closed form (Figure 7E). The hemi-6mApT duplex is first recognized in the base pairing mode by AMTP1 DBD (Figure 7E). Subsequent base flipping, cofactor loading, and AMT1 binding lead to CCC assembly, recognition of hemi-6mApT in the base flipping mode, and methyl transfer (Figure 7E). Flipping back of the newly formed 6mA, CCC disassembly, and cofactor unloading convert the ternary complex in the base flipping mode to the binary complex in the base pairing mode, ready for further translocation along the dsDNA (Figure 7E).

## Materials and Methods

### AMT6/AMT7 complexes structure prediction using AlphaFold3

To model the structures of AMT6/7 complexes, we employed AlphaFold3.0.0^33^, leveraging its diffusion-based architecture for atomic coordinate generation and Pairformer module for modeling protein-nucleic acid interactions. Computations were performed within a Singularity-containerized environment on a high-performance GPU cluster. Structure prediction was executed in five batches of 200 independent prediction seeds to enhance conformational sampling of AMT1 complex. Each batch was submitted via SLURM with resource specifications of 12 CPU cores, 2 NVIDIA A100 GPUs, and 100 GB RAM per node, under a maximum runtime of 48 hours. Input configurations specifying DNA-protein sequences and stoichiometry were defined in customized JSON files, and predictions used default inference settings via run_alphafold.py with validated paths to model/database resources.

### Classify AF3 models by reaction coordinates

To characterize the conformational landscape across a series of protein complexes, we computed interatomic distances between predefined atom pairs representing key structural features. Distance matrices derived from eight distinct structural descriptors were integrated into a unified matrix, where rows correspond to individual AMT6/AMT7 complex structures predicted by AF3 and columns represent the eight structural features. Unsupervised dimensionality reduction and visualization were performed on this integrated distance matrix using Uniform Manifold Approximation and Projection (UMAP) with default parameters (R umap package). UMAP results were divided into 24 groups.

### MD simulations

All molecular dynamics (MD) simulations were performed using GROMACS 2024.3 with GPU acceleration^48^. The CHARMM36 all-atom force field was obtained from the official CHARMM distribution and adapted for use with GROMACS^49^. Parameters for 6mA and SAM were derived within the CHARMM force-field framework and modified to ensure compatibility with GROMACS while preserving the original CHARMM parameterization.

To systematically capture the structural dynamics of the flipped 6mA base, we defined three distinct functional states for the SAM/SAH-bound holo complexes, distinguished by the hybridization and conformation of extrahelical 6mA as specified in PubChem CID 67955. (1) Single-base flipping transition state: systems with 6mA flipped out and stabilized in the sp^3^ hybridized Conformer 3. (2) Single-base flipping ground state: the identical systems but with 6mA in Conformer 1, in which the N^6^ atom adopts an sp^2^ trigonal planar geometry and the N^6^ methyl group is in the stable syn conformation. (3) Double-base paired state: the identical systems but with the 6mA base proximal or distal to SAM/SAH, adopting the sp^2^ hybridized Conformer 2. In addition to these targeted states, a broader panel of AMT7/AMT6–DNA holo complexes was constructed to represent distinct DNA methylation and cofactor binding conditions. DNA substrates were classified as unmethylated (un), hemimethylated (hemi), or fully methylated (full), and simulations were performed in the presence of SAM, SAH, or without a bound cofactor (X), giving the systems A7_un_SAM, A7_hemi_SAM, A7_full_SAH, A7_hemi_X, A7_un_X, A6_hemi_SAM, A6_full_SAH, and A6_un_X.

Initial coordinates were obtained from AlphaFold3 (AF3) generated structural models, with the top-ranked conformations selected based on per-residue pLDDT scores to ensure backbone reliability. Missing hydrogen atoms in the cofactors (SAM, SAH) and in 6mA residues were added using ChimeraX v1.8 to guarantee correct protonation states. The complete structures were then converted into GROMACS compatible topologies with the pdb2gmx module. Each complex was centered in a cubic simulation box with a minimum distance of 1.0 nm between the solute and box boundaries. The systems were solvated with explicit TIP3P water molecules using the CHARMM-compatible solvent model supplied with GROMACS. Potassium and chloride ions were added to neutralize the net system charge and to establish a physiological salt concentration of 0.15 M. Energy minimization was performed using the steepest-descent algorithm for 5,000 steps with positional restraints applied to heavy atoms. This step corrected local deviations in side-chain rotamers of catalytic residues and optimized base-stacking geometries near the flipped base, ensuring that each system resided in a local potential-energy minimum. Following minimization, the systems were equilibrated in two stages with positional restraints maintained on the solute. First, a 125-ps NVT equilibration was conducted using a 1-fs integration time step. Temperature was maintained at 303.15 K using the velocity-rescaling thermostat with separate coupling groups for solute and solvent; initial atomic velocities were assigned from a Maxwell–Boltzmann distribution at 303.15 K with randomized seeds. Subsequently, a 5-ns NPT equilibration was performed using a 2-fs integration time step. Temperature was maintained at 303.15 K using the velocity-rescaling thermostat, and pressure was controlled at 1 bar using the stochastic cell-rescaling (C-rescale) barostat under isotropic coupling with a compressibility of 4.5 × 10^-5^ bar^-1^. Periodic boundary conditions were applied in all three spatial dimensions. For both equilibration stages, short-range electrostatic and van der Waals interactions were evaluated with a 1.2 nm cutoff; van der Waals interactions were treated using a force-switching function between 1.0 and 1.2 nm, while long-range electrostatic interactions were calculated using the particle mesh Ewald (PME) method. Covalent bonds involving hydrogen atoms were constrained using the LINCS algorithm.

After equilibration, positional restraints were removed and production MD simulations were performed under NPT conditions. Temperature and pressure were maintained as in the NPT equilibration step. Long-range electrostatic interactions were treated using PME with a 1.2 nm real-space cutoff. Van der Waals interactions were evaluated using a force-switching scheme between 1.0 and 1.2 nm, and covalent bonds involving hydrogen atoms were constrained using LINCS, enabling a 2-fs integration time step. Production trajectories were generated for durations ranging from 20 ns to 1 μs, depending on the specific complex and simulation objective. Shorter simulations were employed for conformational analyses, whereas longer simulations were conducted to characterize large-scale conformational transitions, DNA base-flipping events, and cofactor-dependent dynamics. Coordinates were saved at regular intervals and analyzed using GROMACS utilities and custom Python scripts.

### Generation of *Tetrahymena* strains

The *AMT1, AMT7* and *AMTP1* mutations were respectively introduced to *AMT1*-NHA, *AMT7*-CHA and *AMTP1*-CHA construct by fusion PCR with primers containing the designed mutations^6^. These constructs were individually transformed into wild type (WT) *Tetrahymena* cells (SB210) and selected in increasing concentration of paromomycin to generate a somatic gene mutation strain^50, 51^. The *AMT6* mutations were introduced to *AMT6*-CHA construct by fusion PCR with primers containing the designed mutations. The constrct was transformed into *AMT7*-KO cells and selected in increasing concentration of blasticidin S to generate a somatic gene mutation strain.

### UHPLC-QQQ-MS/MS analysis

*Tetrahymena* genomic DNA was purified and treated following established protocols^6^. Samples were analyzed by ultra-high-performance liquid chromatography-tandem mass spectrometry (UHPLC-QQQ-MS/MS) on an Acquity BEH C18 column (75 mm × 2.1 mm, 1.7 μm, Waters, MA, USA), using a Xevo TQ-XS triple quadrupole mass spectrometer (Waters, Milford, MA, USA) or a Hypersil GOLD column (100 x 2.1 mm, 1.9 µm, Thermo scientific) using a TSQ Quantiva mass spectrometer (Thermo scientific). The mass spectrometer was set to multiple reaction monitoring (MRM) in positive electrospray mode. The selective MRM transitions were detected under m/z 266/150 for 6mA and m/z 252/136 for dA. The ratio of 6mA/A was quantified by the calibration curves of nucleoside standards running at the same time. UHPLCQQQ-MS/MS was performed at the Key Laboratory of Marine Medicine at Ocean University of China (OUC).

### SMRT sequencing and data analysis

SMRT sequencing data were analyzed as previously described^52, 53^. Only CCS reads supported by ≥ 20 passes were retained to ensure high per-molecule sequencing accuracy. To minimize potential end effects, 20 bp were trimmed from both the 5′ and 3 ′ ends of each read prior to analysis. The trimmed reads were aligned to the reference sequence using BLAST, and only molecules with ≥98% alignment coverage and ≥95% sequence identity were retained. To evaluate the stability of kinetic signals, the standard deviation of IPD-ratio values was calculated from unmodified adenine (IPD ratio ≤ 2.8) within the trimmed terminal regions of each molecule. Molecules were retained only if the standard deviation was ≤0.35. To further remove molecules with localized kinetic artifacts, we identified clusters of non-adenine bases (G, C, or T) exhibiting elevated IPD-ratio values (IPD ratio ≥ 2.8). A cluster was defined as at least three modified non-adenine bases on the same strand, with adjacent bases separated by ≤25 bp. The remaining high-quality Smart-CCS reads were used for all downstream analyses.

Gene-level analysis was performed as previously described^5^ using the same curated set of well-annotated Pol II-transcribed genes. Genes were aligned at their transcription start sites (TSSs), and a region extending 500 bp upstream and 1 kb downstream of the TSS was analyzed. Within this window, the numbers of 6mA calls and total adenine (A) sites were aggregated across all genes at each genomic position. The normalized 6mA level at each position was calculated as the ratio of aggregated 6mA counts to aggregated adenine counts (6mA/A), and these normalized values were used to generate the metagene profiles.

### Recombinant protein purification

Full length AMT1, AMT6, AMT7, AMTP1, AMTP2 ORFs were codon-optimized for bacterial expression as previously mentioned^6^. AMT1 was cloned into a pRSF-Duet vector. AMTP1 and AMTP2 were cloned into a pET-Duet vector. AMT7 was cloned into a pET-28b vector. Mutations were generated by fusion PCR. WT or mutant AMT1 subcomplex components (AMT1, AMTP1 and AMTP2) were co-expressed in *E. coli* BL21 (DE3) cells with a 6 × His-SUMO tag attached to AMT1 and AMTP2 N-termini. Wild type or mutant AMT7 possessing a 6 × His-SUMO tag was expressed in *E. coli* strain BL21 (DE3).

*E. coli* was cultured to an optical density (OD600) of 0.8 at 37°C and protein expression was induced by 0.25 mM IPTG at 16°C overnight. Induced cells expressing subcomplex or AMT7 protein were mixed and harvested. The cells were then resuspended in lysis buffer (50 mM Tris-HCl, pH 8.0, 300 mM NaCl, 25 mM imidazole, 10% glycerol) and homogenized with an ultra-high-pressure cell disrupter at 4°C. Lysates were cleared by centrifugation at 13,000 g for 45 min at 4°C. Proteins were first purified by Ni-NTA (Novagen, USA) affinity chromatography and eluted through on-column tag cleavage by the ULP1 protease. The elute was further purified by passage through a Heparin ion-ex-change column (GE Healthcare, USA) with buffer A (20 mM Tris-HCl, pH 7.5, 50 mM NaCl, 5% glycerol) and buffer B (20 mM Tris-HCl, pH 7.5, 1 M NaCl, 5% glycerol), before being loaded onto a Superdex 200 10/300 Increase column (GE Healthcare, USA) with buffer C (20 mM Tris-HCl, pH 7.5, 100 mM NaCl, and 4 mM DTT).

### Methyltransferase assay

For radioactive methyltransferase assay, 0.1 μM protein was mixed with 1 μM ^3^H-labeled S-adenosyl-L-methionine ([^3^H]SAM, Perkin Elmer) and 1μM dsDNA in 15 μLreaction buffer (50 mM Tris-HCl, pH 7.5, 100 mM NaCl, 1 mM EDTA, 0.05% β-Me). Samples were incubated at 30°C for 30 min and subsequently spotted onto Hybond-XL membrane (GE Healthcare). Membranes were then washed three times with 0.2 M ammonium bicarbonate, once with distilled water, twice with 100% ethanol, and finally air-dried for 1 h. Each membrane was immersed in Ultima Gold (PerkinElmer) and used for scintillation counting on a PerkinElmer MicroBeta2 (PerkinElmer). Statistical significance between two groups was assessed using a one-tailed unpaired Student’s t-test with three independent replicates per group. P-values were calculated accordingly.

### Electrophoretic mobility shift assay (EMSA)

EMSA was performed using dsDNA and proteins purified as above. Briefly, dsDNA (4 μM) was incubated with a gradient dose of 2, 4, and 8 μM full length or truncated AMTP1 protein in buffer containing 50 mM Tris-HCl, pH 7.5, 100 mM NaCl, 1mM EDTA, 0.05% β-Me for 30 min at 4°C. 2 μM AMT7 complex proteins was used as positive control. Reactions were run on a 6% native polyacrylamide gel (acrylamide:bis-acrylamide=37.5:1) in 1 x TBE buffer at 100 V for 30 min. The gel was stained with ethidium bromide.

### In vitro cross-linking of AMT1 complex and DNA

Purified AMT6 and AMT7 complexes were incubated with synthetic oligonucleotides and SAH at 4°C for 30min, and further purified by size exclusion chromatography on a Superdex 200 increase 10/300 column (GE Healthcare). The purified protein-DNA complexes were subjected to cross-linking experiments as per the modified procedures adapted from previously established protocols^6^. Cross-linking mass spectrometry was performed at the Technology Center for Protein Sciences (Proteomics Facility), Tsinghua University.

## Acknowledgements

S.G. was supported by the National Natural Science Foundation of China (32125006), Laoshan Laboratory (LSKJ202203203), and the Natural Science Foundation of Shandong Province (ZR2024ZD40). Y.L. was supported by the NSF award (MCB-2435178). M.X. were supported by the National Key R&D Program of China (2025YFC2817103). High-performance computing resources for data processing were provided by the Center for Advanced Research Computing (CARC) at the University of Southern California, as well as the Institute of Evolution and Marine Biodiversity IEMB1, the High-Performance Biological Supercomputing Center, and Marine Big Data Center of Institute for Advanced Ocean Study at OUC. Our special thanks are given to Prof. Weibo Song (OUC) for his helpful suggestions during drafting the manuscript.

## Author contribution

Y.L. and S.G. conceived the study and designed the experiments; B.N. and L.N. performed most of the experiments; W.Y. performed AlphaFold 3 prediction; W.Y. and J.N. analyzed SMRT data; Y.W. collected and organized the data. M.X. performed molecular dynamics simulation; D.W. and W.Y. performed conformational heterogeneity analysis; W.G., Z.C. and G.Z performed the radioactive methylation assay; B.N., Y.L. and S.G. wrote the paper, with inputs from all authors. All authors read and approved the final manuscript.

The authors declare no conflict of interest.

## Data availability

The AlphaFold3 predicted structures and SMRT sequencing data presented in this work are available on Zenodo at https://doi.org/10.5281/zenodo.21684694

## Supplemental tables

Table S1. AF3 models with their ipTM scores.

Table S2. Percentage of base flipping and base pairing in various AF3 input settings.

Table S3. Key inter-atomic distances used to sample the conformational space of the AMT1 complex.

## Supplemental files

File S1. AMT1 sequence alignment.

File S2. AMT6/7 sequence alignment.

File S3. AMTP1 sequence alignment.

