## Supplemental Figures for "Structural basis of DNA N^6^-adenine methylation in eukaryotes"

#These authors contributed equally

*Corresponding author:

**Supplemental Figures 1-15**

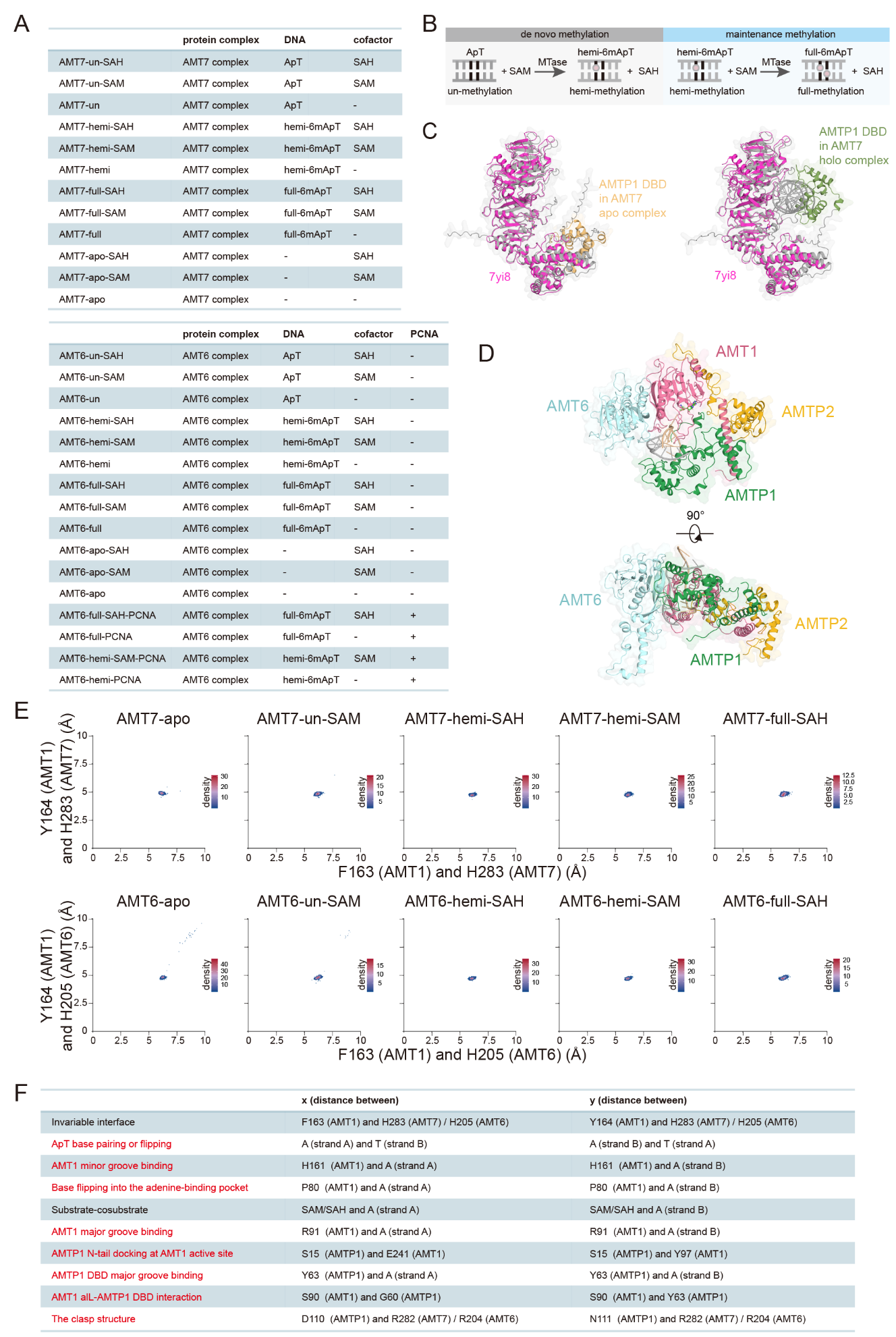

**Figure S1. Structural modeling of the AMT1 complex.**

A. AF3 modeling input settings of the AMT1 apo and holo-complexes.
B. Schematic for *de novo* and maintenance methylation of the dsDNA at the ApT duplex. Unmodified ApT is methylated by the MTase using SAM to form hemi-6mApT (*de novo* methylation), which can be further methylated to full-6mApT (maintenance methylation).
C. Structural superimposition of the AF3-predicted structures of the AMT7 apo- and holo-complexes with the previously solved structure of the AMT7 apo-complex (7yi8, magenta)^1^. Note the differential placement of the AMTP1 DBD.
D. Front (top) and side (bottom) views of the AMT6 ternary complex (AMT6-hemi-SAM). The DNA target and non-target strands are colored in silicon and silver, respectively.
E. Conformational analysis tracking the interaction between the AMT1 *Minor Groove Insertion Loop* and AMT6/7 *Multiple interaction Loop*. Top: the distance between AMT7 H283 (C_α_) and AMT1 F163/Y164 (C_α_, x/y-axis, Å); bottom: the distance between AMT6 H205 (C_α_) and AMT1 F163/Y164 (C_α_, x/y-axis, Å); colored as per local density of the AF3 models.
F. List of the critical interfaces and associated inter-atomic distances. Interfaces used for UMAP analysis are highlighted in red.

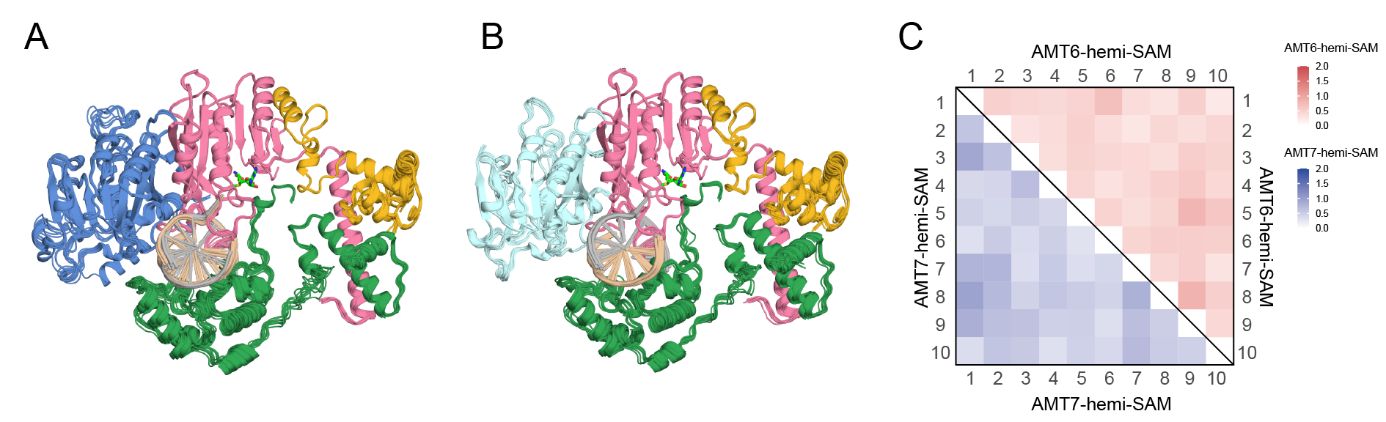

**Figure S2. AF3 modeling of the AMT1 ternary complex.**

A. Structural overlay of the AMT7 ternary complex (AMT7-hemi-SAM). Top-10 AF3 models (ranked by ipTM), all in the base flipping mode.
B. Structural overlay of the AMT6 ternary complex (AMT6-hemi-SAM). Top-10 AF3 models (ranked by ipTM), all in the base flipping mode.
C. Pairwise comparison of the top-10 AF3 models. Heatmap: RMSD (C_α_), color scale (Å). Lower-left: the AMT7 ternary complex (AMT7-hemi-SAM). Upper-right: the AMT6 ternary complex (AMT6-hemi-SAM).

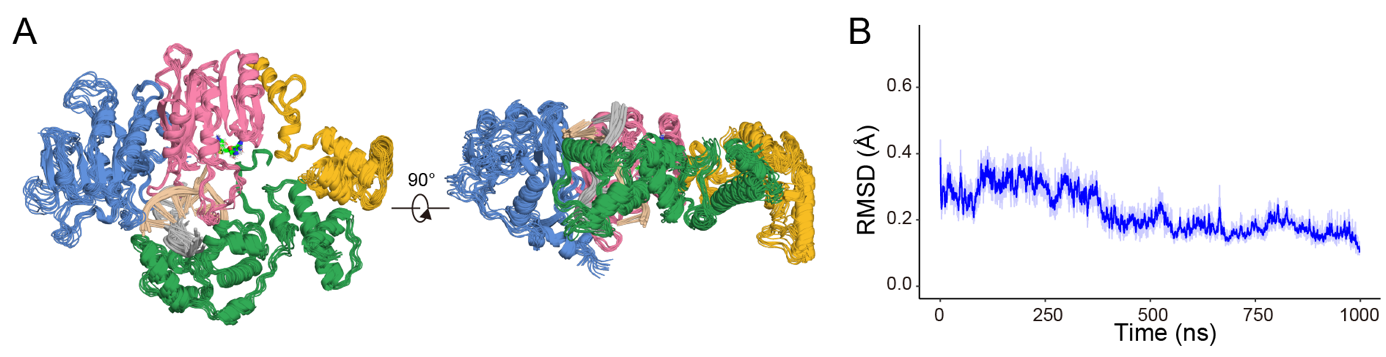

**Figure S3. MD simulations of the AMT1 ternary complex.**

A. Front (left) and side (right) views of the AMT7 ternary complex (AMT7-hemi-SAM). Overlay of 10 representative structures from the last 100 ns of the MD trajectory.
B. MD trajectory for the AMT7 ternary complex (AMT7-hemi-SAM). RMSD values (C_α_, Å) are calculated after all structures are aligned to the last frame. The light-colored trace represents raw frame-by-frame values; the dark-colored trace represents the 20-frame moving average.

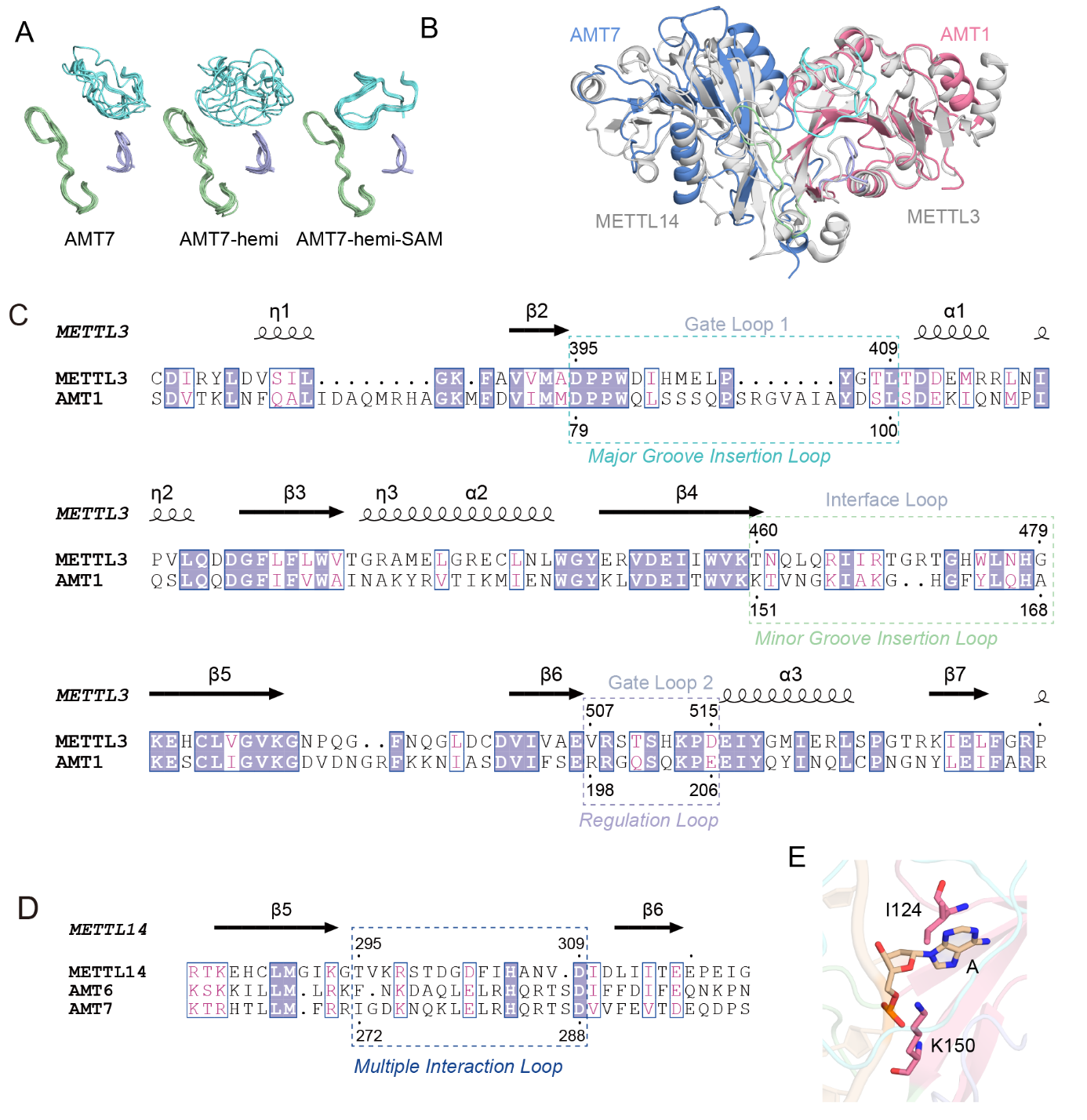

**Figure S4. Comparison of the AMT1 and METTL3-METTL14 complexes.**

A. Structural overlay of the three AMT1 substrate-binding loops: the *Major Groove Insertion Loop* (saxe blue), the *Minor Groove Insertion Loop* (dark sea green), and the *Regulation Loop* (lavender). Top-10 AF3 models (ranked by ipTM) of the AMT1 apo-complex (AMT7), the AMT1 binary complex (AMT7-hemi, base pairing), and the AMT1 ternary complex (AMT7-hemi-SAM, base flipping) are aligned by AMT1.
B. Structural overlay of the AMT1 ternary complex (AMT7-hemi-SAM) and the METTL3-METTL14 complex (5il1)^2^.
C. Structure-based sequence alignment of AMT1 and METTL3. Numbered according to AMT1 (truncated) and METTL3 (5il1). AMT1 *Major Groove Insertion Loop*, *Minor Groove Insertion Loop*, and *Regulation Loop* are boxed and labeled.
D. Structure-based sequence alignment of AMT6, AMT7, and METTL14. Numbered according to AMT7 (truncated) and METTL14 (5il1). AMT6/AMT7 *Multiple interaction Loop* is boxed and labeled.
E. AMT1 I124 and K150 interacting with the sugar-phosphate backbone of the extrahelical adenine.

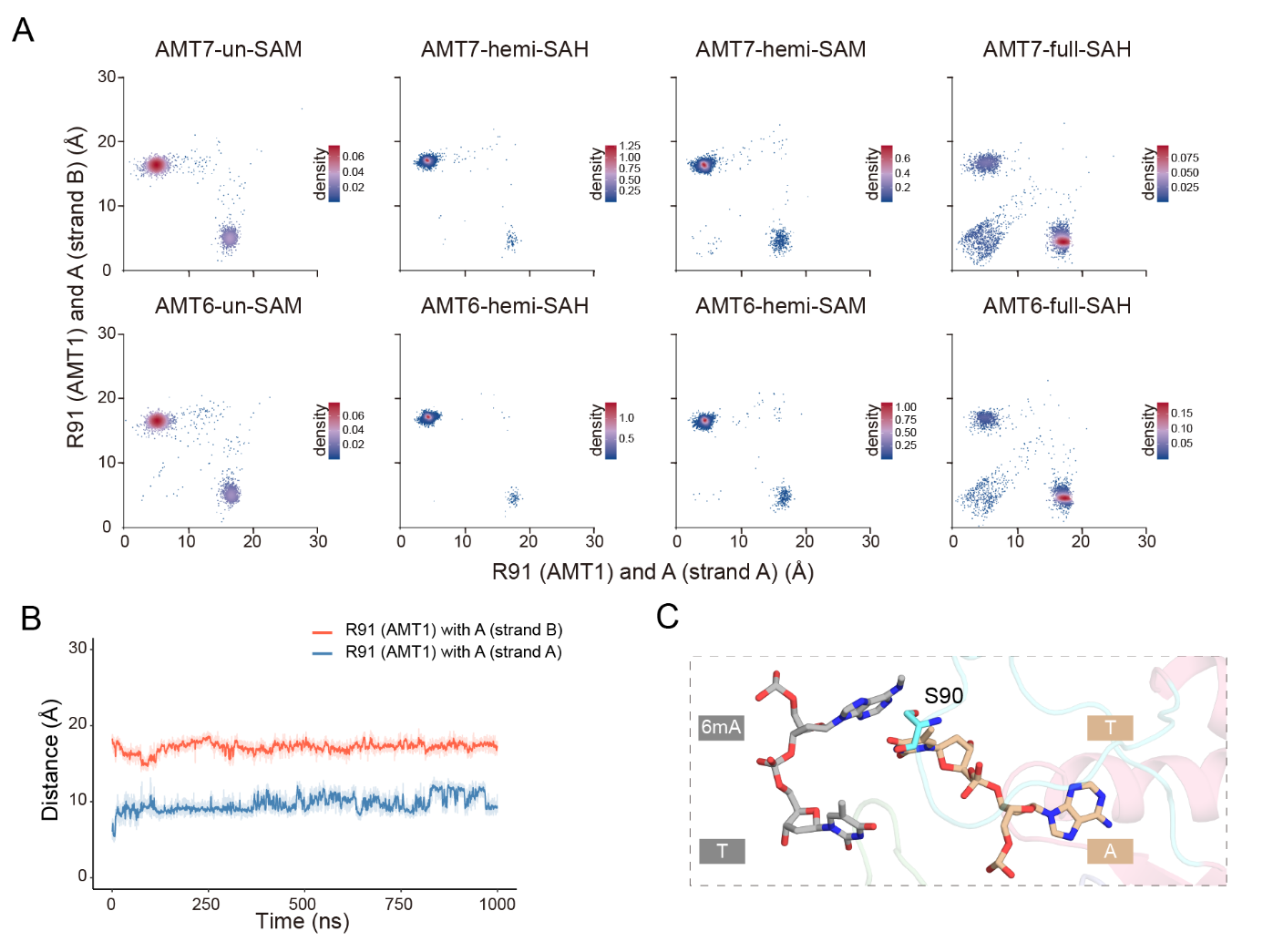

**Figure S5. ApT duplex recognition by AMT1.**

A. Conformational analysis tracking the distance between AMT1 R91 (C_α_) and A/6mA (N^6^) on DNA strand A/B (x/y-axis, Å) in the AMT1 ternary complex; colored as per local density of AF3 models.
B. MD trajectory (AMT7-hemi-SAM) tracking the distance between AMT1 R91 (C_α_) and A/6mA (N^6^) on DNA strand A/B. The light-colored traces represent raw frame-by-frame values; the dark-colored traces represent the 20-frame moving average.
C. Contacts between DNA and AMT1 S90, located at the distal portion of the AMT1 *Major Groove Insertion Loop* (AMT7-hemi-SAM). Note the interaction between AMT1 S90 and the 6mA N^6^-methyl group (in the *anti*-conformation required for base pairing with the target strand thymine^3^).

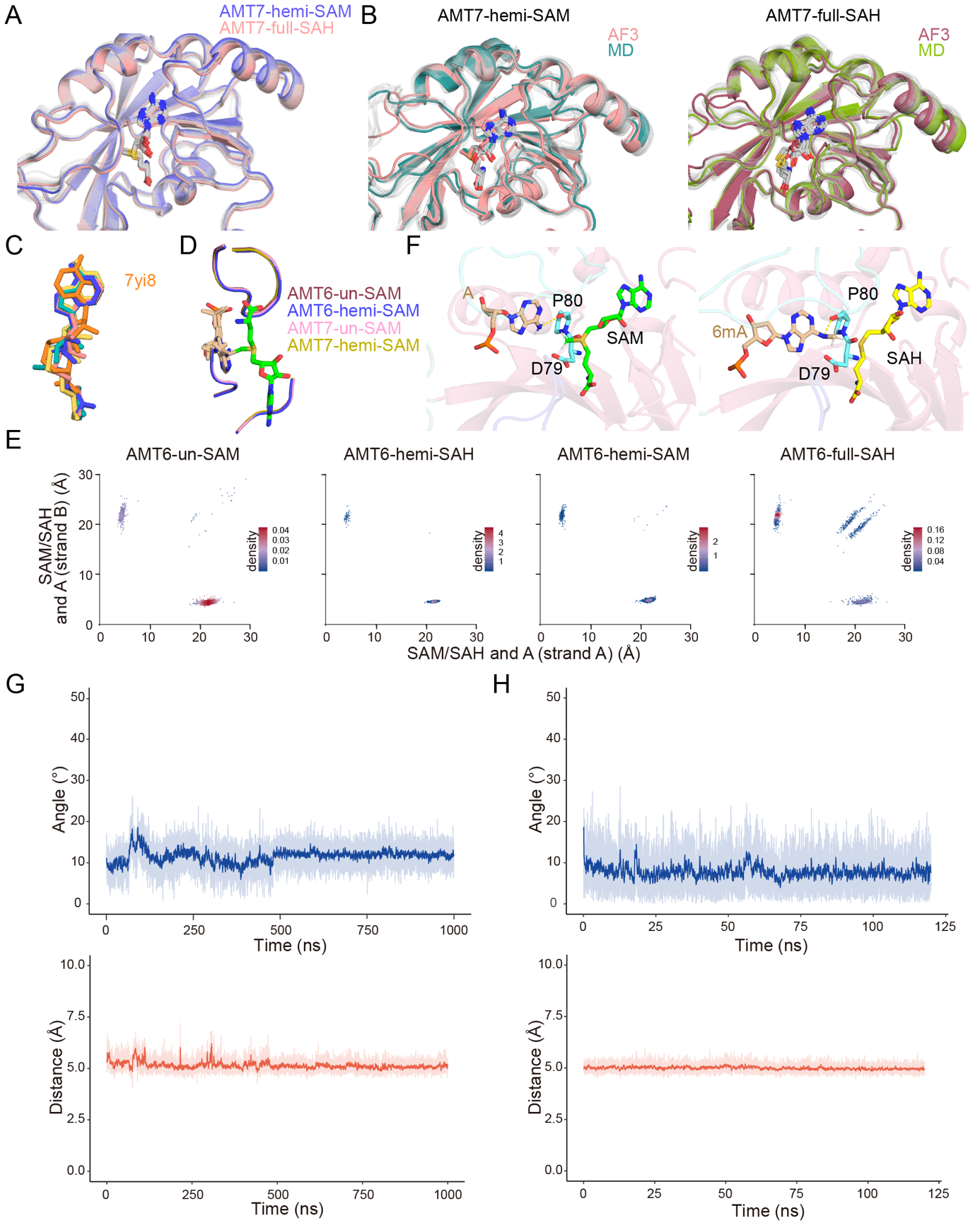

**Figure S6. AMT1 active site.**

A. Consistent SAM/SAH placement in AF3 models of the AMT1 ternary complex (AMT7-hemi-SAM and AMT7-full-SAH). Top 10 scored AF3 models are aligned by AMT1. The highest-ranking model for each complex is shown in color, whereas the remaining models are shown in gray. SAM/SAH: stick model.
B. Comparison of SAM/SAH placement in representative AF3 and MD models of the AMT1 ternary complex (AMT7-hemi-SAM and AMT7-full-SAH). For each MD trajectory, the structure from the last frame is shown in color, whereas other sampled MD structures are shown in gray. SAM/SAH: stick model.
C. Comparison of SAM/SAH placement in the AMT1 ternary complex and the apo-complexes (7yi8, 7f4n and 5il2). SAH placement in 7yi8 is an outlier.
D. Structural overlay of the MD models for the AMT1 ternary complex (AMT6/AMT7-un-SAM, AMT6/AMT7-hemi-SAM, ES state). Note the consistent placement of the extrahelical adenine, SAM, and the surrounding loops in the active site.
E. Conformational analysis tracking base flipping into the active site. The AMT6 ternary complexes corresponding to the ES/EP state for *de novo* and maintenance methylation. Scatterplot: distance between cofactor (S_δ_) and A/6mA (N^6^) on DNA strand A/B (x/y-axis, Å); colored as per local density of the AF3 models.
F. Structural details of the AMT1 active site in the ES (left) and EP (right) states. AMT1 D79 and P80 are shown alongside A/6mA and SAM/SAH, in stick models. Yellow dashed line: hydrogen bond between P80 and A/6mA.
G. Tracking the C_me_-N^6^-S_δ_ angle and the N^6^-S_δ_ distance in the MD trajectory of the ES complex (AMT7-hemi-SAM). The light-colored traces represent raw frame-by-frame values; the dark-colored traces represent the 20-frame moving average.
H. Tracking the C_me_-N^6^-S_δ_ angle and the N^6^-S_δ_ distance in the MD trajectory of the EP complex (AMT7-full-SAH, 6mA protonated).

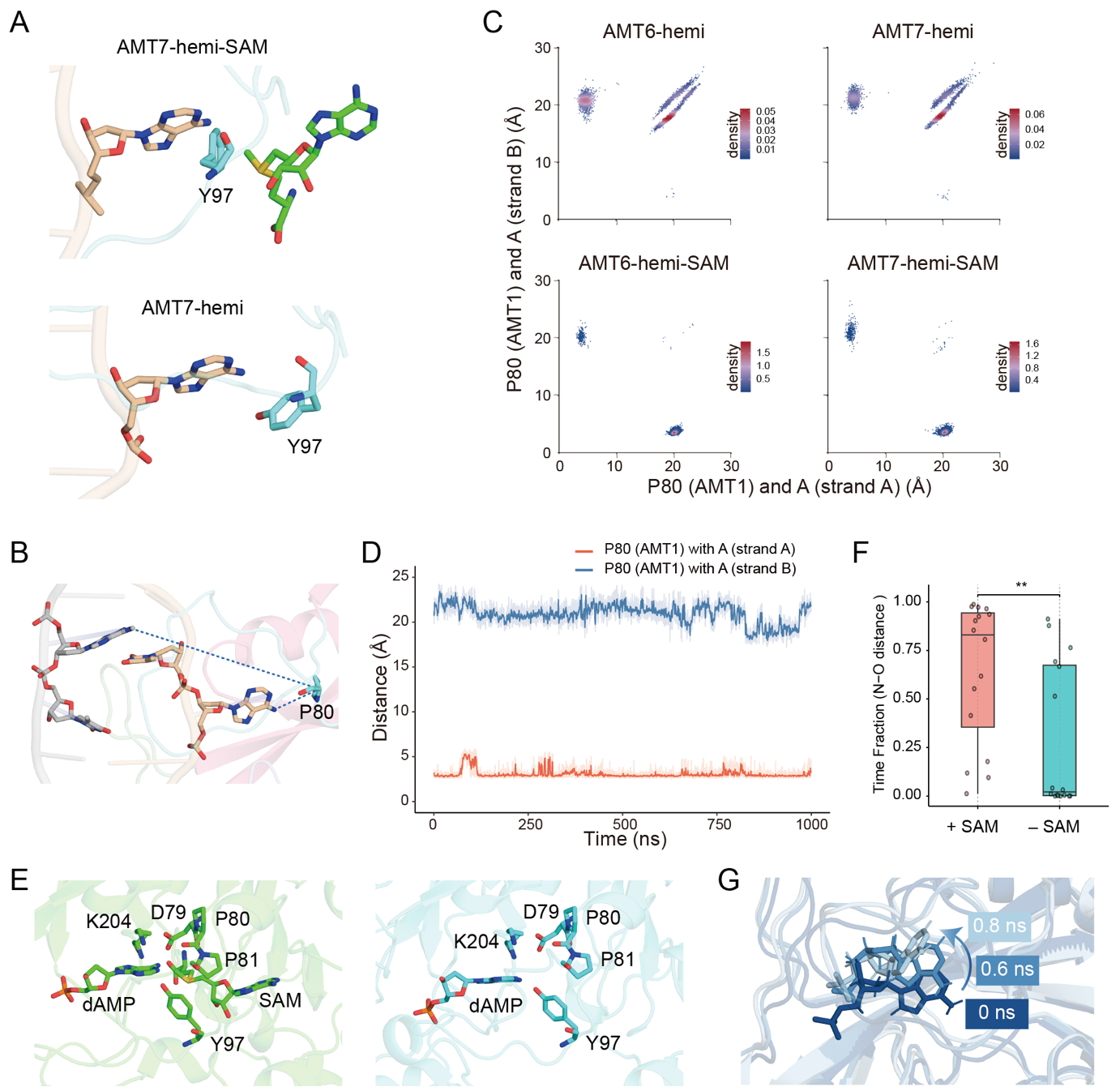

**Figure S7. Cooperation between adenine and cofactor binding.**

A. The active site within the MD models for AMT7-hemi-SAM (top) and AMT7-hemi (bottom). Note that in AMT7-hemi, AMT1 Y97 is no longer proximally positioned next to the target adenine, as in AMT7-hemi-SAM.
B. The distance between AMT1 P80 (O) and A/6mA (N^6^) on the target and non-target strands are shown respectively as the short and long dashed lines in the base flipping mode for AMT7-hemi-SAM.
C. Conformational analysis tracking DNA binding by AMT1 *Major Groove Insertion Loop*. AMT6 and AMT7 binary and ternary complexes (AMT6/AMT7-hemi and AMT6/AMT7-hemi-SAM). Scatterplot: distance between AMT1 P80 (O) and A/6mA (N^6^) on DNA strand A/B (x/y-axis, Å); colored as per local density of AF3 models.
D. MD trajectory (AMT7-hemi-SAM) tracking the distance between AMT1 P80 (O) and A/6mA (N^6^). The light-colored traces represent raw frame-by-frame values; the dark-colored traces represent the 20-frame moving average.
E. Starting structures of AMT7-dAMP-SAM and AMT7-dAMP for MD stimulations. Critical residues of the AMT1 adenine-binding pocket as well as dAMP are shown in the stick model.
F. dAMP retention in the adenine-binding pocket of AMT7-dAMP-SAM and AMT7-dAMP: summary of 16 independent MD trajectories. The y-axis indicates the fraction of simulation time during which dAMP remained within the specified N-O distance range (2.2-2.6 Å), reflecting its retention in the pocket.
G. Representative snapshots from the MD trajectory (AMT7-dAMP) showing adenine dissociation from its binding pocket. The progressive move of the adenine away from the pocket is stabilized by its contacts with hydrophobic residues in the AMT1 *Major Groove Insertion Loop*.

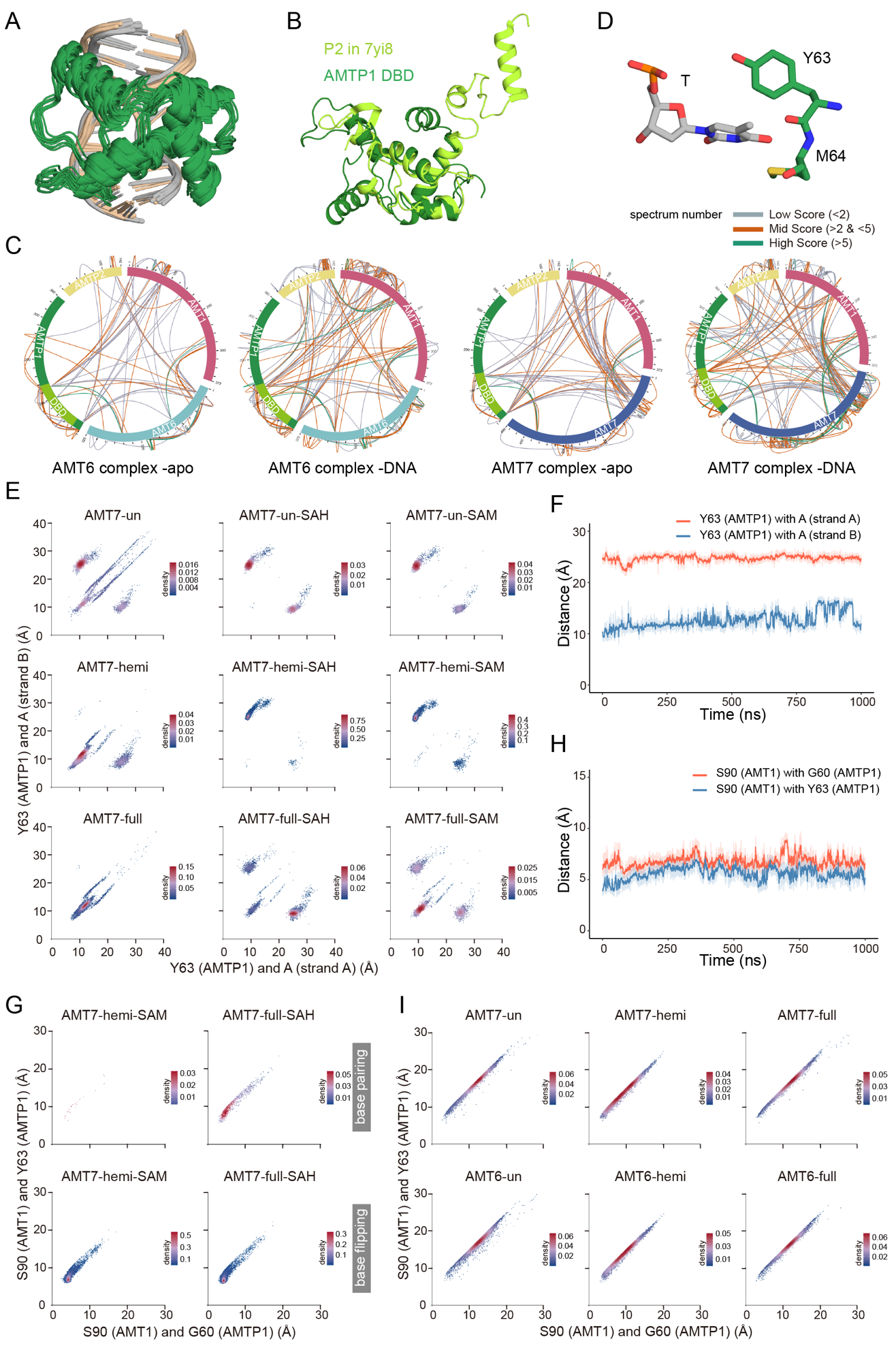

**Figure S8. AMTP1 DBD and its cooperative binding mode.**

A. Structural overlay of the AMTP1 DBD in the AMT1 ternary complex (AMT7-hemi-SAM). Top 10 AF3 models (ranked by ipTM), MD-optimized, all in the base flipping mode.
B. Structural overlay of AMTP1 DBD domain (predicted) and AMTP2 HD (7yi8).
C. Crosslinking Mass Spectrometry (XL-MS) results of the AMT7 or AMT6 complex. Circle plot: number of the corresponding crosslinks (spectral count) indicated by the line color; FDR≤5%.
D. AMTP1 residues in proximity to the non-target strand.
E. Conformational analysis tracking DNA binding by AMTP1 DBD in the AMT1 holo-complex. Scatterplot: distance between AMTP1 Y63 (C_α_) and A/6mA (N^6^) on DNA strand A/B (x/y-axis, Å); colored as per local density of AF3 models.
F. MD trajectory (AMT7-hemi-SAM) tracking the distance between AMTP1 Y63 (C_α_) and A/6mA (N^6^) on DNA strand A/B. The light-colored traces represent raw frame-by-frame values; the dark-colored traces represent the 20-frame moving average.
G. Conformational analysis tracking interactions between the AMT1 *Major Groove Insertion Loop* and the AMTP1 DBD. Scatterplot: distance between AMT1 S90 (C_α_) and AMTP1 G60/Y63 (C_α_) (x/y-axis, Å); colored as per local density of AF3 models. The base flipping/pairing modes within the AMT1 ternary complex (AMT7-hemi-SAM, AMT7-full-SAH) are plotted separately.
H. MD trajectory (AMT7-hemi-SAM) tracking the distance between AMT1 S90 (C_α_) and AMTP1 G60/Y63 (C_α_). The light-colored traces represent raw frame-by-frame values; the dark-colored traces represent the 20-frame moving average.
I. Conformational analysis tracking interactions between the AMT1 *Major Groove Insertion Loop* and the AMTP1 DBD within the AMT1 binary complex (AMT6/AMT7-un/hemi/full).

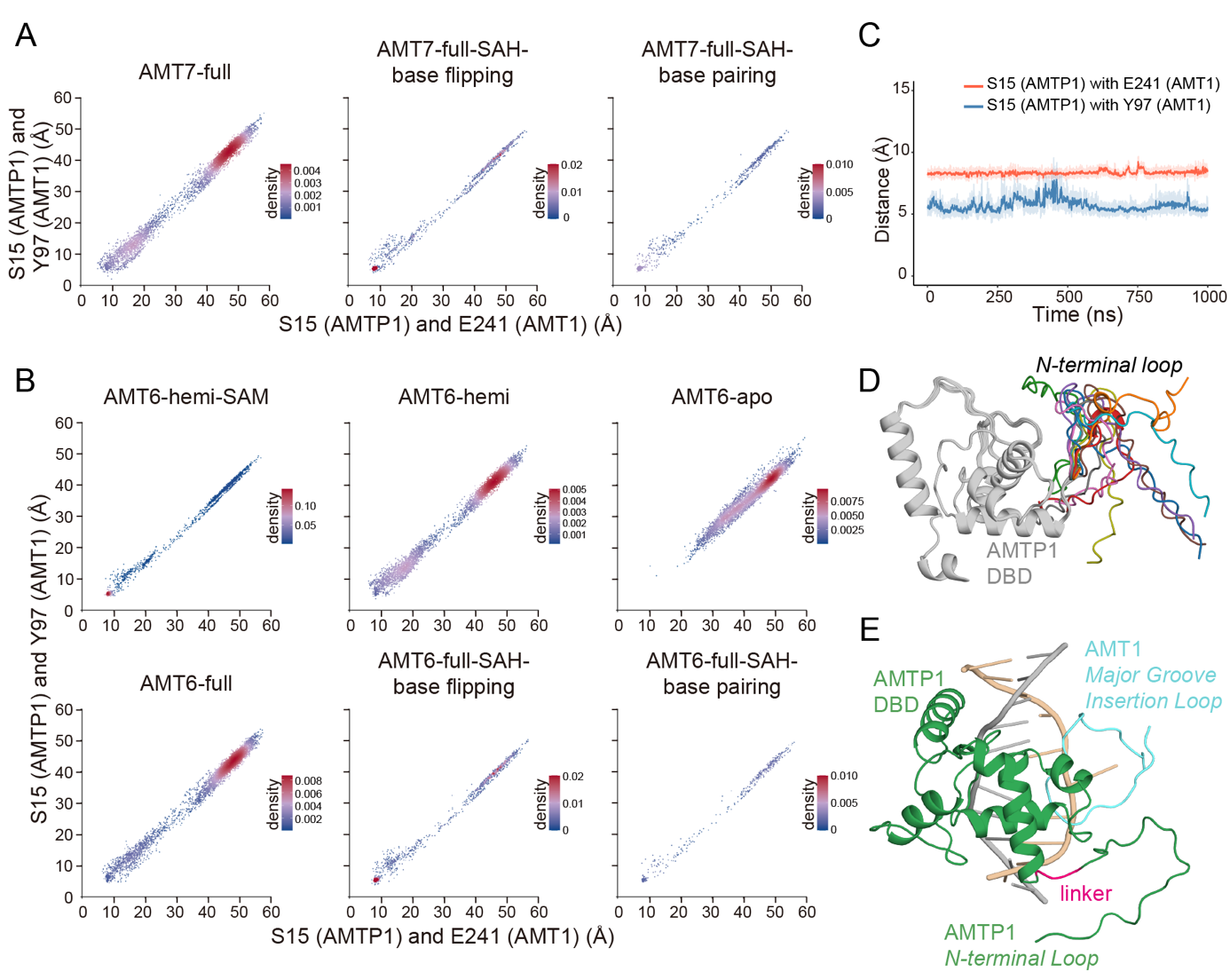

**Figure S9. AMTP1 *N-terminal Loop* and its role in cooperative binding.**

A. Conformational analysis tracking the AMTP1 *N-terminal Loop* docking at the AMT1 active site. AF3 models for the AMT1 binary complex (AMT7-full) and ternary complex (AMT7-full-SAH). Scatterplot: the distance between AMTP1 S15 (C_α_) and AMT1 E241/Y97 (C_α_) (x/y-axis, Å); colored as per local density of AF3 models. The base flipping/pairing modes within the AMT1 ternary complex (AMT7-full-SAH) are plotted separately.
B. Conformational analysis tracking the AMTP1 *N-terminal Loop* docking at the AMT1 active site in the AMT6 apo-complex, binary holo-complex, and ternary holo-complex.
C. MD trajectory (AMT7-hemi-SAM) tracking the distance between AMTP1 S15 (C_α_) and AMT1 E241/Y97 (C_α_). The light-colored traces represent raw frame-by-frame values; the dark-colored traces represent the 20-frame moving average.
D. Structural overlay of the AMTP1 DBD and its N-terminal tail. The AMTP1 DBD is shown in gray, whereas the AMTP1 N-terminal tail is shown in different colors. Within the AMT1 binary complex (AMT7-hemi), the AMTP1 N-terminal tail loses its structure when not docked at the AMT1 active site.
E. Ribbon representation of AMTP1 *N-terminal Loop*, DBD domain and AMT1 *Major Groove Insertion Loop*.

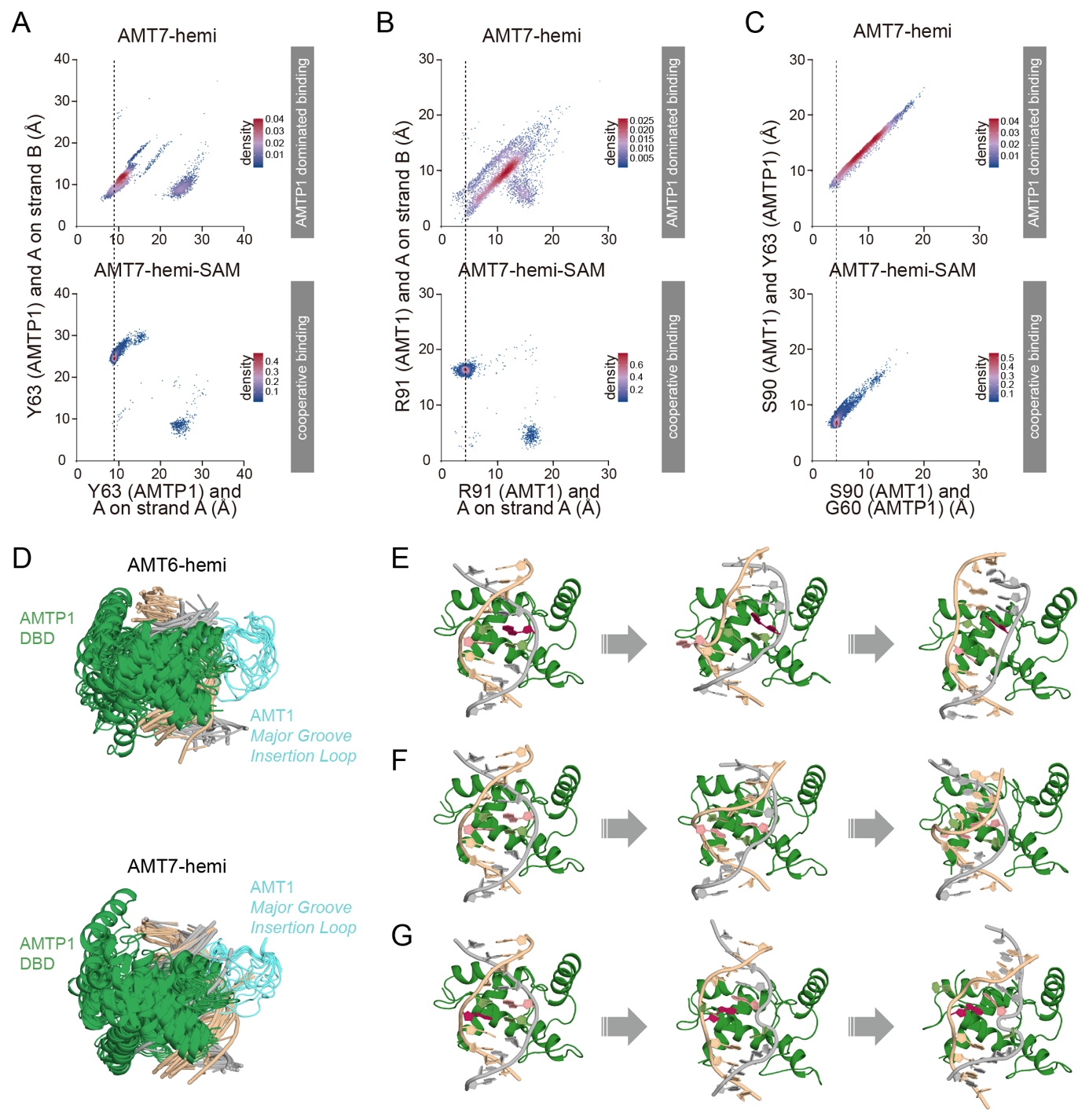

**Figure S10. AMTP1-dominated DNA binding.**

A. Conformational analysis tracking DNA binding by the AMTP1 DBD in the AMT1 complex (top: AMT7-hemi; bottom: AMT7-hemi-SAM). Scatterplot: distance between AMTP1 Y63 (C_α_) and A/6mA (N^6^) on DNA strand A/B (x/y-axis, Å); colored as per local density of AF3 models.
B. Conformational analysis tracking DNA binding by the AMT1 *Major Groove Insertion Loop* in the AMT1 complex (top: AMT7-hemi; bottom: AMT7-hemi-SAM). Scatterplot: distance between AMT1 R91 (C_α_) and A/6mA (N^6^) on DNA strand A/B (x/y-axis, Å).
C. Conformational analysis tracking the interaction between the AMT1 *Major Groove Insertion Loop* and the AMTP1 DBD in the AMT1 complex (top: AMT7-hemi; bottom: AMT7-hemi-SAM). Scatterplot: distance between AMT1 S90 (C_α_) and AMTP1 G60/Y63 (C_α_, x/y-axis, Å).
D. Structural overlay of AMTP1 DBD in the AMTP1-dominated binding mode. Ten AF3 models each for the AMT6 binary complex (AMT6-hemi, left) and the AMT7 binary complex (AMT7-hemi, right). AMT1 *Major Groove Insertion Loop* exhibits structural heterogeneity, after losing most specific contacts with the AMTP1 DBD and the DNA major groove.
E. Three MD models for the AMTP1 DBD in association with the hemi-methylated dsDNA. The left is the starting structure; the right is the final structure (200 ns); the middle is a representative MD model along the MD trajectory.
F. Three MD models for the AMTP1 DBD in association with the unmethylated dsDNA.
G. Three MD models for the AMTP1 DBD in association with the symmetry-related hemi-methylated dsDNA. 6mA: the target strand; A: the non-target strand.

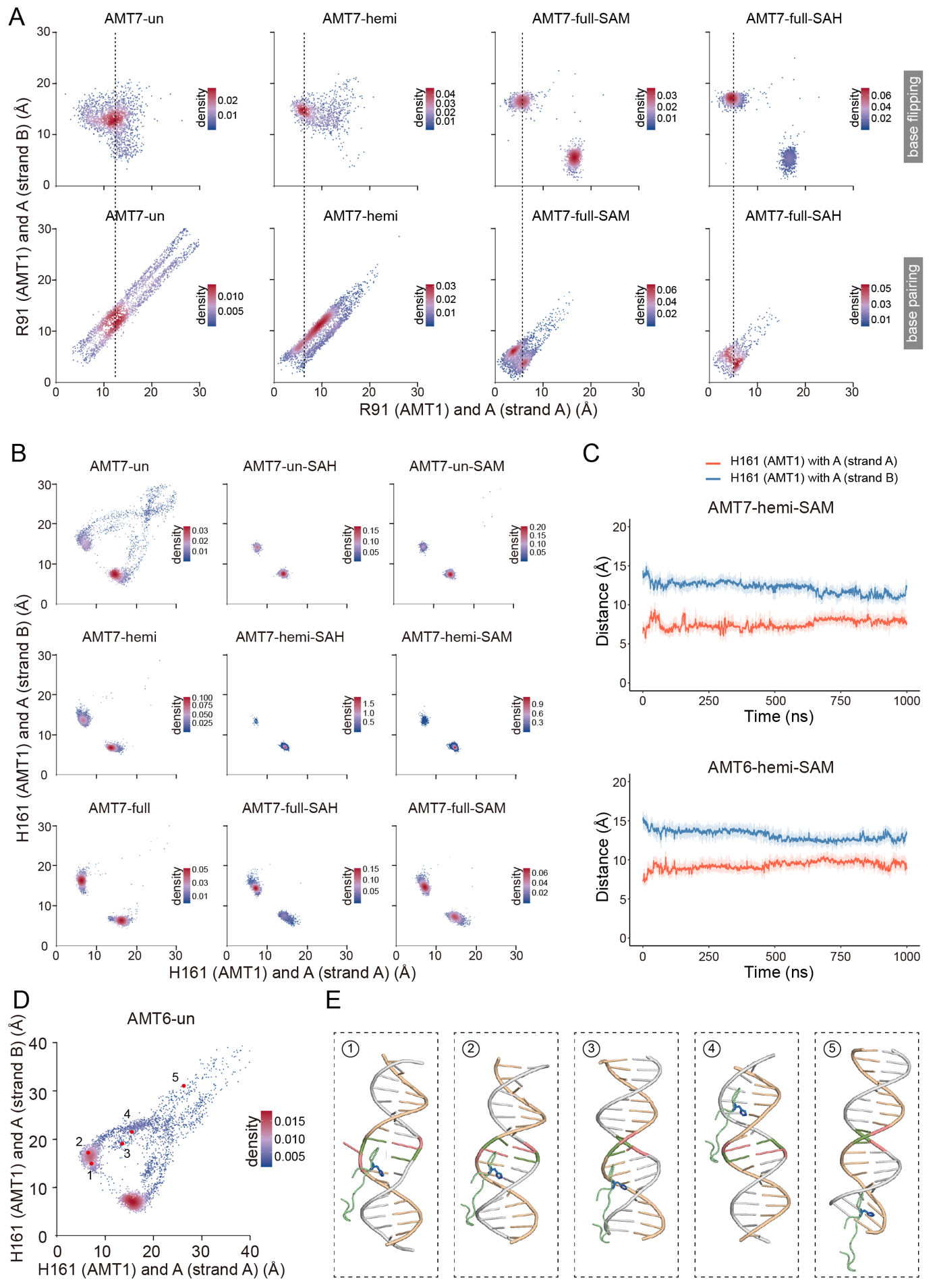

**Figure S11. Offsite DNA binding.**

A. Conformational analysis tracking DNA binding by the AMTP1 DBD. Scatterplot: distance between AMT1 R91 (C_α_) and A/6mA (N^6^) on DNA strand A/B (x/y-axis, Å); colored as per local density of AF3 models. The base flipping/pairing modes within the AMT1 binary (AMT7-un/hemi/full) and ternary complexes (AMT7-full-SAM/SAH) are plotted separately.
B. Conformational analysis tracking DNA binding by the AMT1 *Minor Groove Insertion Loop* in all 9 AMT7 binary and ternary complexes (AMT7-un/hemi/full-null/SAM/SAH). Scatterplot: distance between AMT1 H161 (C_α_) and A/6mA (C3’) on DNA strand A/B (x/y-axis, Å); colored as per local density of AF3 models. H161 is asymmetrically positioned in the minor groove, much closer to the target strand (5-6 Å) than to the non-target strand (14-16 Å).
C. MD trajectories of the AMT1 ternary complex (AMT6/AMT6-hemi-SAM) tracking the distance between AMT1 H161 (C_α_) and adenine (C3’). The light-colored traces represent raw frame-by-frame values; the dark-colored traces represent the 20-frame moving average.
D. Conformational analysis tracking DNA binding by the AMT1 *Minor Groove Insertion Loop* in the AMT6 binary complex (AMT6-un).
E. Correponding AF3 models in the AMT6 binary complex (AMT6-un).

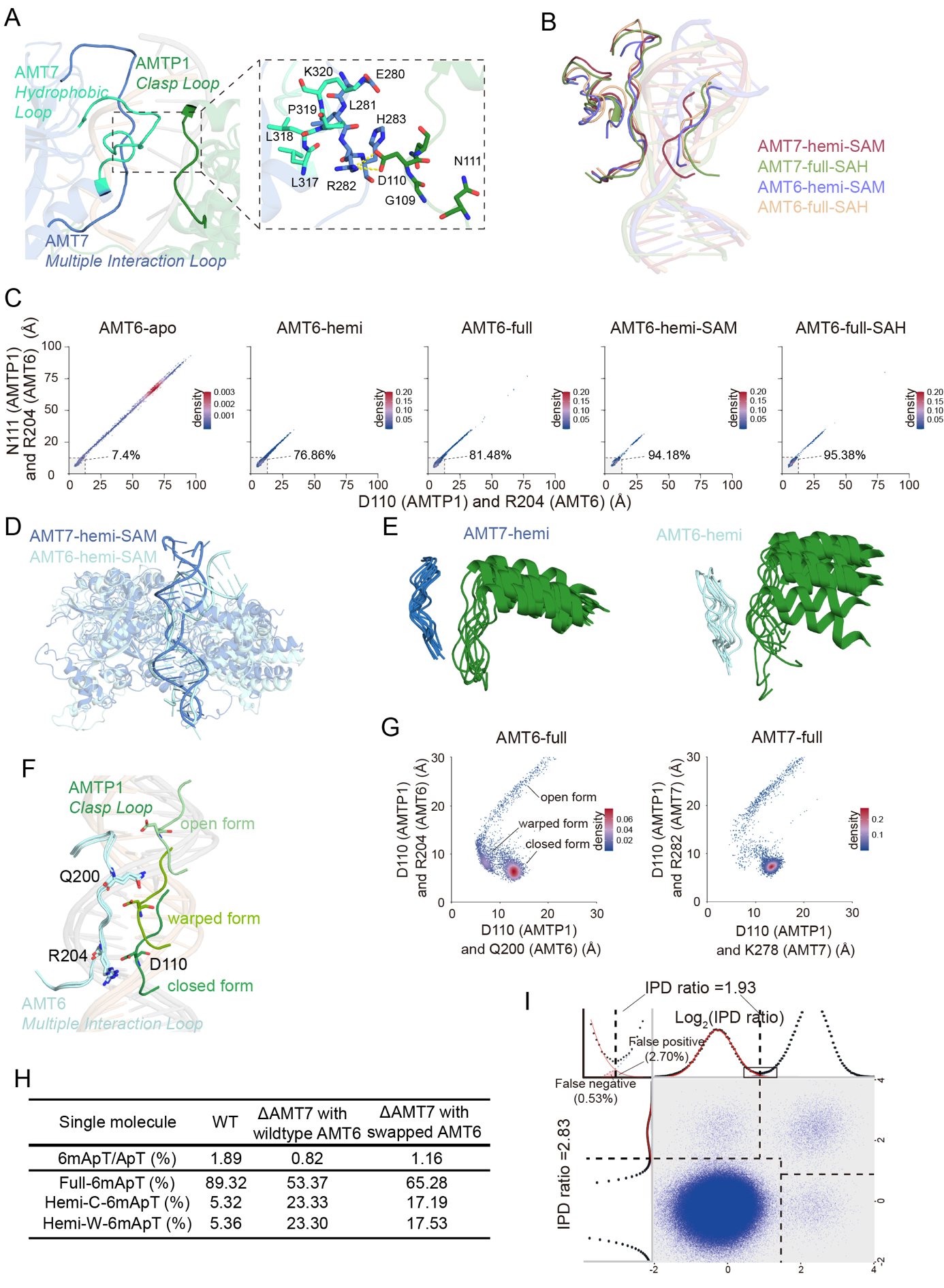

**Figure S12. Clasp formation in the AMT6 and AMT7 complexes.**

A. The clasp securing the closed form AMT1 ternary complex (AMT7-hemi-SAM). Three important loops are highlighted: the AMTP1 *Clasp Loop*, the AMT7 *Multiple Interaction Loop*, and the *Hydrophobic Loop*. Expanded view: residues critical for clasp formation. Yellow dashed line: hydrogen bond.
B. Structural overlay of the clasp in the AMT6/AMT7 ternary complexes. MD-optimized AF3 models for AMT7-hemi-SAM, AMT7-full-SAH, AMT6-hemi-SAM, and AMT6-full-SAH, in the base flipping mode.
C. Conformational analysis tracking clasp formation in the AMT6 complex (AMT6, AMT6-hemi, AMT6-full, AMT6-hemi-SAM, AMT6-full-SAH). Scatterplot: distance between AMT6 R204 (C_α_) and AMTP1 D110/N111 (C_α_; x/y-axis, Å); colored as per local density of AF3 models. Percentage of AF3 models consistent with clasp formation is indicated.
D. Structural overlay of the AMT1 ternary complex MD models (AMT6/AMT7-hemi-SAM). Note the severe dsDNA bending in the AMT6, but not AMT7, complex.
E. Structural overlay of AMT7 (left) and AMT6 (right) *Multiple Interaction Loop* and AMTP1 DBD of the top 10 scored AF3 models for the respective binary complex (AMT7/AMT6-hemi) in the base pairing mode.
F. Differential interactions between the AMT6 *Multiple Interaction Loop* and the AMTP1 *Clasp Loop*, within the closed (lower), warped (middle), and open (upper) forms. Three representative AF3 models for the AMT6 binary complex (AMT6-full) are aligned by AMT6.
G. Conformational changes in the clasp of the binary complex (AMT6/AMT7-full). Scatterplot: distance between AMTP1 D110 and AMT6 Q200/AMT7 K278, AMT6 R204/AMT7 R282 (all C_α_; x/y-axis, Å). The regions corresponding to the closed/warped/open forms are indicated.
H. Summary of PacBio sequencing results.
I. Demarcation of the four methylation states of ApT duplexes in AMT6-sw by their IPD ratio (IPDr) on W- and C-strand, respectively. For bulk ApT duplexes, the IPDr threshold for 6mA calling was set at 2.83, according to deconvolution based on Gaussian fitting of the small 6mA peak. For ApT duplexes with one 6mA, the IPDr threshold for calling 6mA on the opposite strand was set at 1.93, according to deconvolution based on Gaussian fitting of the small unmodified A peak.

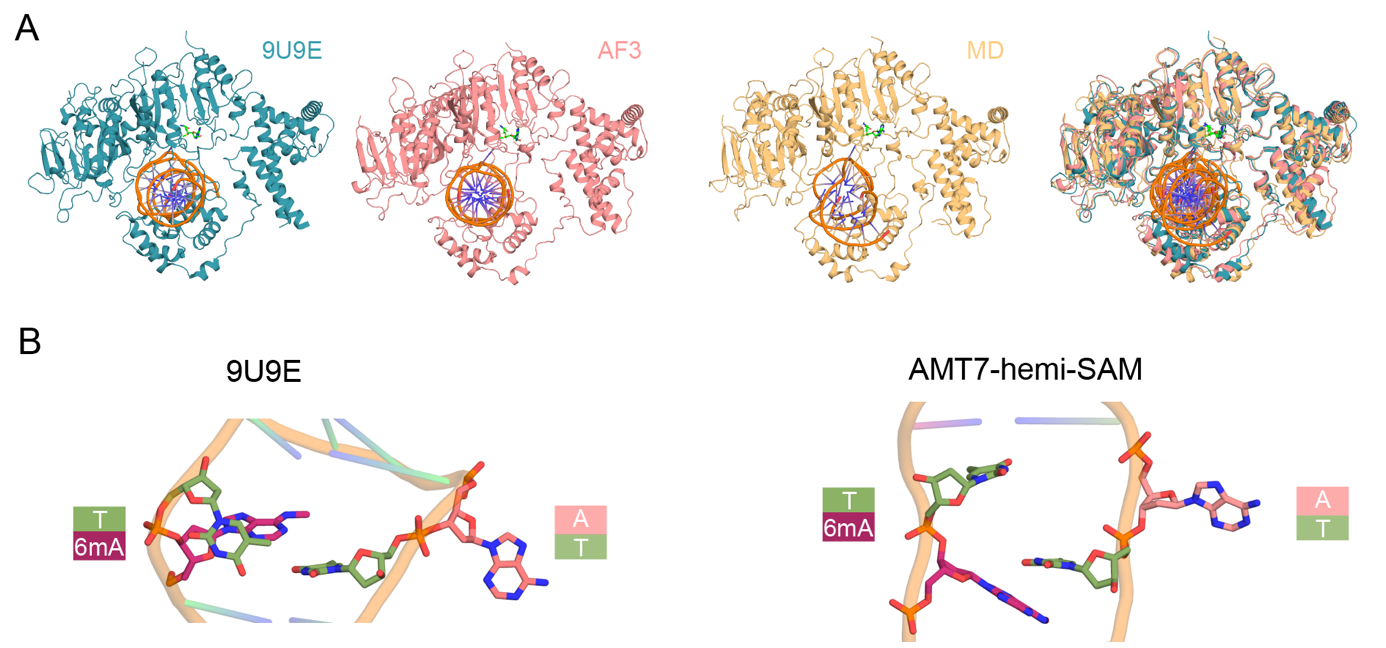

**Figure S13. Side-by-side comparison of the AMT1 ternary complex structural models.**

A. Side-by-side comparison of the AMT1 ternary complex structural models: the cryo-EM structure (AMT6-hemi-SAM cryo-EM, 9U9E), the AF3 model after initial MD solvation and equilibration (AMT6-hemi-SAM, AF3), and the AF3 model after extended MD simulations (AMT6-hemi-SAM, MD). Overlay of the three structures is on the right.
B. Comparison of the base flipping conformations in the cryo-EM structure (PDB: 9U9E) and the AF3 model (AMT7-hemi-SAM). AF3 predicts flipping of the target adenine only, whereas the cryo-EM structure additionally shows partial flipping of the opposing non-target thymine.

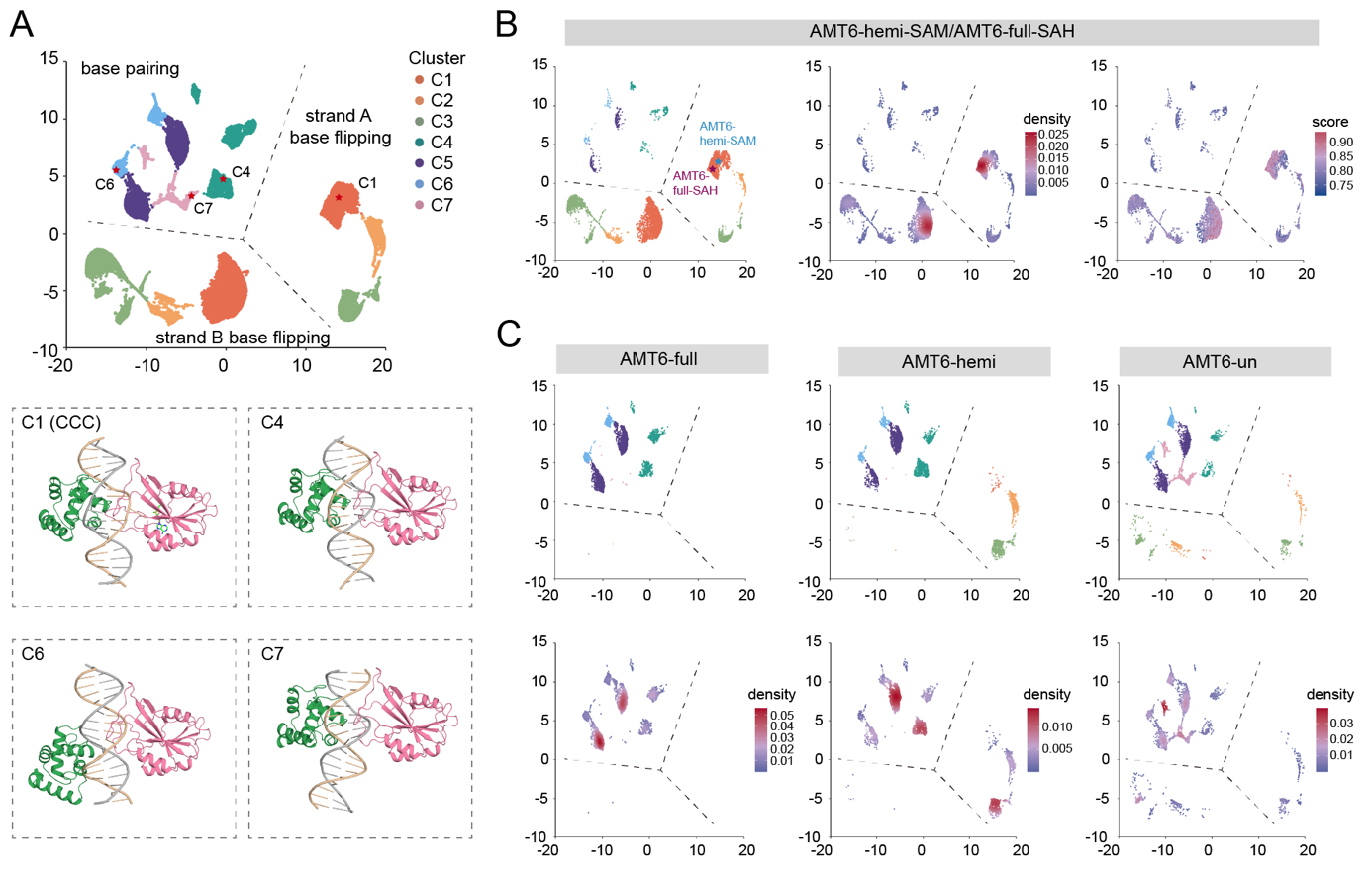

**Figure S14. UMAP analysis of the AMT6 holo-complex.**

A. UMAP analysis of the AMT6 holo-complex. 7 clusters (C1 to C7), featuring distinct structural characteristics, are demarcated and differentially colored. Demarcations separating the base pairing mode and the base flipping mode are shown; the latter is further divided by selection of the target strand for base flipping (A/B). Below: representative structures in cluster C1 (CCC), C4 (AMTP1-dominated binding), C6 (AMTP1 offsite binding), and C7 (AMT1 offsite binding); only the AMTP1 DBD, AMT1 MT-A70 domain, and dsDNA are shown.
B. UMAP analysis of the AMT6 ternary complex (AMT6-hemi-SAM and AMT6-full-SAH). Colored by cluster (left), density (middle), and AF3 score (right).
C. UMAP analysis of the AMT6 binary complex (AMT6-full, AMT6-hemi, and AMT6-un). Colored by cluster (top) and density (bottom).

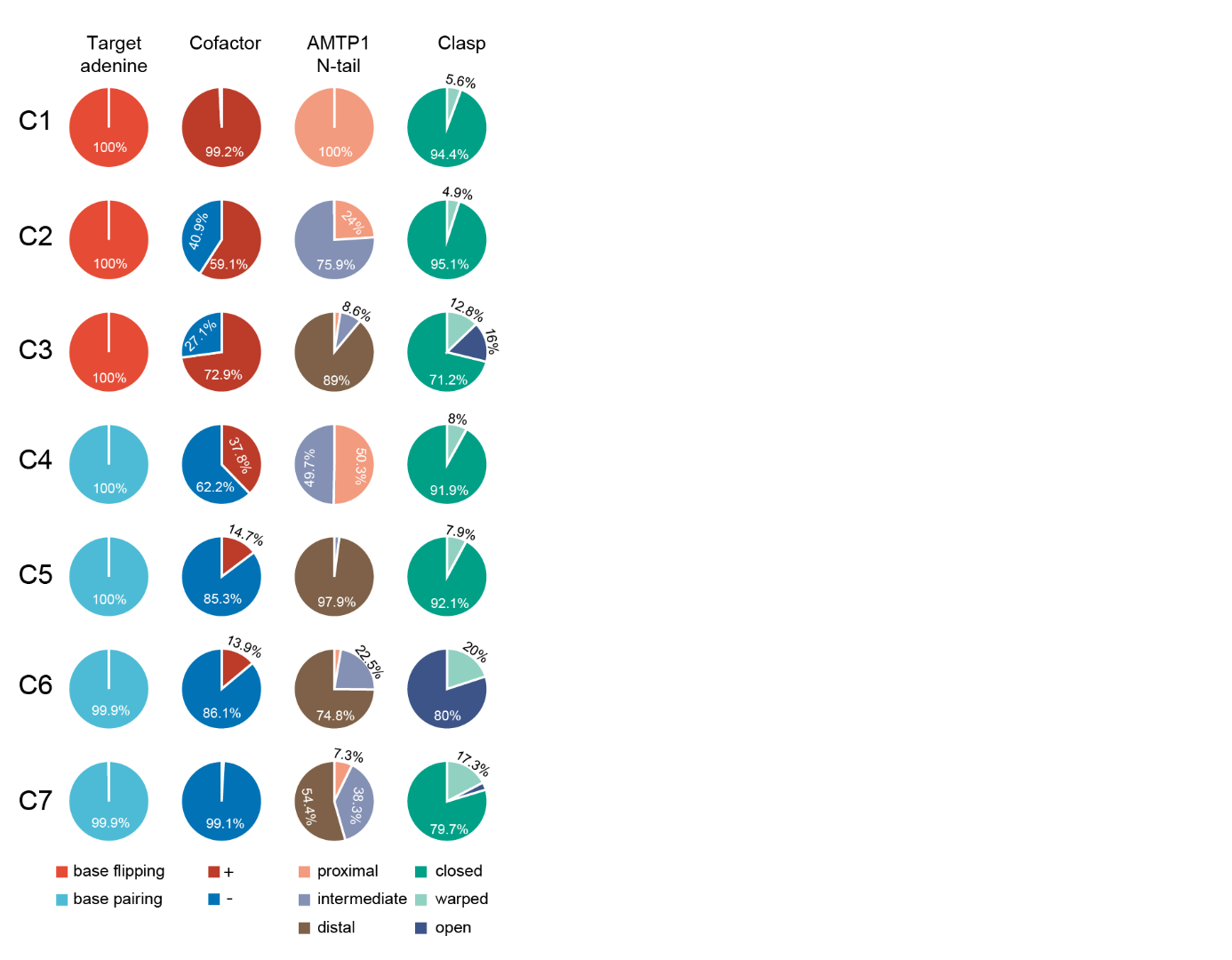

**Figure S15. Structural features of the seven AF3 model clusters.**

Distribution of key structural features in the seven AF3 model clusters (C1–C7). Pie charts summarize the conformational states of the target adenine, cofactor, AMTP1 N-tail, and clasp in each cluster. Percentages indicate the fraction of models exhibiting each state.
