## Supplemental file S1 for "Structural basis of DNA N^6^-adenine methylation in eukaryotes"

```
                          1        10        20        30        40        50        60
Oxy.tri_42188             MNQSSQDITTQKSSNGFNPQTQPETLIQVIRKESTFIFKYRKNPYYVPPPISSQTSPNLE
Oxy.tri_21422             MESTKNARGNSRDHNNSGASGADAATNSKKKNQPALTQFFKKQPEGAIPKGS........
Cam.sin_17781             ............................................................
Cam.sin_15892             ............................................................
Tet.the_AMT1              ............................................................
Tet.mal_00157670_PEP_     ............................................................
Tet.ell_00199380_PEP_     ............................................................
Tet.vor_00155570_PEP_     ............................................................
Tet.pyr_00179540_PEP_     ............................................................
Tet.can_00119340_PEP_     ............................................................
Tet.bor_00147860_PEP_     ............................................................
Tet.epi_00127660_PEP_     ............................................................
Tet.epi_00126830_PEP_     ............................................................
Tet.par_00146520_PEP_     ............................................................
Par.bur_P08360024         ............................................................
Par.bur_P03800026         ............................................................
Pse.per_1                 ............................................................
Par.bur_P04570033         ............................................................
Crei_1335048048           ............................................................
Crei_159474530            ............................................................
Arob_1183359562           ............................................................
Arep_1183482424           ............................................................

                                   70        80        90       100       110       120
Oxy.tri_42188             VETSNDLNQMSDYEGQIPNNYEINRNSTQFTNNDDQSDNDFYDNNSITTMQIDTSTAKIL
Oxy.tri_21422             .EANNRTSEQSNSNASANANHHSEHVQTRQSAQLSQQNRDSQNTKVNQINQNGATGVNSL
Cam.sin_17781             ............................................................
Cam.sin_15892             ............................................................
Tet.the_AMT1              ............................................................
Tet.mal_00157670_PEP_     ............................................................
Tet.ell_00199380_PEP_     ............................................................
Tet.vor_00155570_PEP_     ............................................................
Tet.pyr_00179540_PEP_     ............................................................
Tet.can_00119340_PEP_     ............................................................
Tet.bor_00147860_PEP_     ............................................................
Tet.epi_00127660_PEP_     ............................................................
Tet.epi_00126830_PEP_     ............................................................
Tet.par_00146520_PEP_     ............................................................
Par.bur_P08360024         ............................................................
Par.bur_P03800026         ............................................................
Pse.per_1                 ...........................................................M
Par.bur_P04570033         ............................................................
Crei_1335048048           ............................................................
Crei_159474530            ............................................................
Arob_1183359562           ............................................................
Arep_1183482424           ............................................................

                                  130       140       150       160       170       180
Oxy.tri_42188             NNGPLEYNPDLPNKEQKLKDSQVMQNQPPTATSTNSQQRTLQELINIMPSIEDISQQCKQ
Oxy.tri_21422             LQTPKKKLLETPSKKKTYADE...........PDEIDMREIQSLVDKKVKESAAAQQQLS
Cam.sin_17781             .......MFRFKDIENVTKKLLLNQILIYSFLCSLSRAVMLICIIIAIFDRYTIKYKINV
Cam.sin_15892             ............................................................
Tet.the_AMT1              .......MKKEQQFLIFKKSLIIAQKRKEINIKQLKQQFKNFLFVQIFSIIKLKLQDIII
Tet.mal_00157670_PEP_     ............................................................
Tet.ell_00199380_PEP_     ......................................SNLKIFCQDFYIIKLNLQDIII
Tet.vor_00155570_PEP_     ............................................................
Tet.pyr_00179540_PEP_     ............................................................
Tet.can_00119340_PEP_     ............................................................
Tet.bor_00147860_PEP_     ............................................................
Tet.epi_00127660_PEP_     ............................................................
Tet.epi_00126830_PEP_     ............................................................
Tet.par_00146520_PEP_     ............................................................
Par.bur_P08360024         ............................................................
Par.bur_P03800026         ............................................................
Pse.per_1                 ASYLQKQIQQKNNKLKNSDKDIQNKSLNIKRTIKNIEQDKNKNKIKEKLDDSKIKGRKQV
Par.bur_P04570033         ............................................................
Crei_1335048048           ............................................................
Crei_159474530            ............................................................
Arob_1183359562           ............................................................
Arep_1183482424           ............................................................
```

```
                              190       200       210       220       230       240
Oxy.tri_42188              QQQLKIQAKANSTQSASTANAANGGKGRKRGRTVRFDQPLLGKVRQRNGDASDDEEPDEI
Oxy.tri_21422              QSGIEDSAIKRSLRPRKVENYKNMLEG.........DEITLKTIQD..............
Cam.sin_17781              QEFELLFCIFLFIFNHEKPSFFKIIILY................................
Cam.sin_15892              ............................................................
Tet.the_AMT1               KFKMSKAVNKKGLRPRKSDSILDHIKNK................................
Tet.mal_00157670_PEP_      ...MSKAVNKKGLRPRKSDSILDHIKNK................................
Tet.ell_00199380_PEP_      KFKMSKAVNKKGLRPRKSDSILDHIKNK................................
Tet.vor_00155570_PEP_      ...MSKAATKKGLRPRKSDSLVDHIKNK................................
Tet.pyr_00179540_PEP_      ...MSKVATKKGLRPRKSDSLLDHIKNK................................
Tet.can_00119340_PEP_      ...MSKATSKKGLRPRKSDSLLDHIKDK................................
Tet.bor_00147860_PEP_      ...MSKAASKKGLRPRKSDSLLDHIKDK................................
Tet.epi_00127660_PEP_      ...MSKATTKKGLRPRKSDSVFDKIKNK................................
Tet.epi_00126830_PEP_      ...MSKAITKKGLRPRKSDSVFDHIKNK................................
Tet.par_00146520_PEP_      .....MSKVKKVQKGKKTESVFDHIKSK................................
Par.bur_P08360024          ............................................................
Par.bur_P03800026          ............................................................
Pse.per_1                  KKSEESENDNDNEHTSNEEEEQDEQTSK................................
Par.bur_P04570033          .......................MQGRR................................
Crei_1335048048            ............................................................
Crei_159474530             ............................................................
Arob_1183359562            ............................................................
Arep_1183482424            ............................................................

                              250       260       270       280       290       300
Oxy.tri_42188              EMLIRRLHTDILNDARNDPVEQAKKIRQARESQSDQTNSTTQLSVYERMILGSASQQSTD
Oxy.tri_21422              ....................EQIEVKRKKREASSQNR.......................
Cam.sin_17781              ............................................................
Cam.sin_15892              ............................................................
Tet.the_AMT1               ............................................................
Tet.mal_00157670_PEP_      ............................................................
Tet.ell_00199380_PEP_      ............................................................
Tet.vor_00155570_PEP_      ............................................................
Tet.pyr_00179540_PEP_      ............................................................
Tet.can_00119340_PEP_      ............................................................
Tet.bor_00147860_PEP_      ............................................................
Tet.epi_00127660_PEP_      ............................................................
Tet.epi_00126830_PEP_      ............................................................
Tet.par_00146520_PEP_      ............................................................
Par.bur_P08360024          ............................................................
Par.bur_P03800026          ............................................................
Pse.per_1                  ............................................................
Par.bur_P04570033          ............................................................
Crei_1335048048            ............................................................
Crei_159474530             ............................................................
Arob_1183359562            ............................................................
Arep_1183482424            ............................................................

                              310       320       330       340       350       360
Oxy.tri_42188              HQPGEFSNMFRTLEDEQIEINQNFLFDEYDSEDDSIADDKVEIASDDEQMLLQEHKKRGK
Oxy.tri_21422              ............LEDEDEDEDMLEVGQQIERASDDEDDDDFPISTRRS....ARKRTRRQ
Cam.sin_17781              ...........FIFKAPFSSTMPKLKKKIKTEKVEEDNLHYGLRPKKM......TSIFQN
Cam.sin_15892              .........................MKKLKKVKNEEGILPTDIIQTKE......TLLSAT
Tet.the_AMT1               ...........LDQEFLEDN....ENGEQSDEDYDQ...KSLNKAKKP......YKKRQT
Tet.mal_00157670_PEP_      ...........LDQEFLEDN....ENGEQSDEDYDQ...KSLNKAKKP......YKKRQT
Tet.ell_00199380_PEP_      ...........LDQEFLEDN....ENGEQSDEDYDQ...KSLNKSKKP......YKKRQT
Tet.vor_00155570_PEP_      ...........LDQEFLDDP..DADAGEYSEEEYDA...KDLKKSKRP......YKKRQV
Tet.pyr_00179540_PEP_      ...........LDQEFLDEPDAEVDAGELSEEEFES...KSLKKSKRP......YKKRQV
Tet.can_00119340_PEP_      ...........LDEEFLNDP....DAGELSEEEYDGN..KSSKKGKRP......YKKRQV
Tet.bor_00147860_PEP_      ...........LDEEFLNDP....DAGELSEEEYDGN..KSSKKGKRP......YKKRQV
Tet.epi_00127660_PEP_      ...........LTEELYEDP....DAGELSEECQEADDKKQIKKGKRP......YKKRQL
Tet.epi_00126830_PEP_      ...........LDDEFLEDP....DAGELSEEEYDNDDKKAIKKGKRP......YKKRQV
Tet.par_00146520_PEP_      ...........LDDDFLDQE......DQDSDYEKPDNKKKAAIKGSKP......YKKRQT
Par.bur_P08360024          .........................................MISKQKG......NKKQKI
Par.bur_P03800026          .........................................MISKQKG......NKKSKI
Pse.per_1                  ...........NSSENENDGENSDQLEENSEEEEESSDDDNNSNNNSN......NSEQNQ
Par.bur_P04570033          ...........SQLDKKTQPKPTTNGKQHKVSNKSKKIRKNDNNNHHS......HVANGN
Crei_1335048048            ............................MATLPGAAAAAPGANAEVGV........PEPS
Crei_159474530             ............................MATLPGAAAAAPGANAEVGV........PEPS
Arob_1183359562            .......................MDEKEVENSVLDSSNIEKSNATTSN........MDVD
Arep_1183482424            ................MTSDTSAMTADVLNRKRKRSPAMNGDDLSNNS........DEAD
```

```
                              370       380       390       400         410
Oxy.tri_42188                 KYLQDEIVKEEDFDEDDDSDEDIHMDDLENESLSFDRNNRK....SHKPVCKRTREENIL
Oxy.tri_21422                 DVDEDEEAIEVNQVESSDAEVEIPANDIDTESYTEGTNKRKQKLKAKKQVLDKKKNKTEG
Cam.sin_17781                 LNVEETELRDLREGDIN.......FEEDYEEETKSR.KRKRKKRVDILEDDFEYQKEEDD
Cam.sin_15892                 LVNPVEFKDIIDEVEFN.......VESEEEDEEDNGDKRKRKKKVDILDDEVEFDKEDDE
Tet.the_AMT1                  QNGSELVISQQKTKAK........ASANNKKSAKNSQKLDEEEKIVEEEDLSPQKNGAVS
Tet.mal_00157670_PEP_         QNGSELVVSQQKTKAK........ATVNAKKSAKNNQKLSEDEKIVEDEDLSPQKNGVGS
Tet.ell_00199380_PEP_         QNGSELVVFQQKTKAK........PSVAPKKNTKAQKQNDEEKIVEEEEGLSPQKNGVGS
Tet.vor_00155570_PEP_         ENGSEVVIPSSASKTKPAKEVSSSTNGSKKSTKNSQKSSEEEVFVEDGGANSPQKNEAGL
Tet.pyr_00179540_PEP_         ENGSEVVIAQQKSKSK.....EATNGSAKKTTKNSQKSSEEEVFVDDGGASSPKKNEGDS
Tet.can_00119340_PEP_         ENGSEIIVSAQKTKSK.......DTPVIKKNTKNSQKSSEEEEVILEEGV.SPQKHEVGS
Tet.bor_00147860_PEP_         ENGSEIVISAQKTKSK.......DTPVIKKNTKNSQKSSEEEEVILEEGG.SPQKHEVGS
Tet.epi_00127660_PEP_         TNVAEEVVPAEKHKAK.........AGRNSKKSQIINDDEIVEQQEEEDKPQKKNEADLT
Tet.epi_00126830_PEP_         ENVAEEVVSVEKQSKS.........KEVTNGKQHSKESTKISD..EEDESPKINESVYTD
Tet.par_00146520_PEP_         NEEDEEEEVVMASDSG..........EMEISTEKQKRQRKSKYDKVQQKETKDTKIVQE
Par.bur_P08360024             VEDEDQIQKKVKLDDD......QSVSQILEEEDIDSEEIEGLDVSEEEKEPIEDNN....
Par.bur_P03800026             VEDEDQIQKKVKLDDD......QSVSQILEEEDIDSEEIEGLDVSEEEKEPIEDNN....
Pse.per_1                     QRQKRSAVKKIKSYKK.....LKNLQYKDEIDIDEEELNGVDDMNEINGNQNDNSKLIN
Par.bur_P04570033             GKGSSQLQPEAVESIN........FNGNYNLRQKKHKNYLDYGSDQGRDENDLDFGIKQL
Crei_1335048048               LEPQDALQQRIALAEGL.......LALNEADAMQAWQQLP...................
Crei_159474530                LEPQDALQQRIALAEGL.......LALNEADAMQAWQQLP...................
Arob_1183359562               ETSNNETSTAIIKSEDG.......ANSYDDFLKLDFTPE..................EE
Arep_1183482424               NNTTTGTTTSVDSNEND.......YQEQDREPILRLPRLNDAKLLEEVVDDVDYEDQPER

                         420       430       440       450       460
Oxy.tri_42188                 DADLGDEKDDEDTIFIDNLPSDEFSIRRQLQDVKSYIKQFEMLFFEEED...........
Oxy.tri_21422                 DIDKEDAVEEEETVFIDNLPNDEFEIRRMLKEVKKHIKSLEKQFFEEED...........
Cam.sin_17781                 HIDIDED.LYDDNLVLDKLPKKRAELEKLYEKVNQRIAIYKKQFFNEQI...........
Cam.sin_15892                 HIEIDDDDLYDDNLILDKLPKKKHELEELYETVKRKIIIYKKQFFHEQE...........
Tet.the_AMT1                  EDDQQQEASTQEDDYLDRLPKSKKGLQGLLQDIEKRILHYKQLFFKEQN...........
Tet.mal_00157670_PEP_         EDDQQQDASTQEDDYLDRLPKSKKGLQSLLQDIEKRILHYKQLFFKEQN...........
Tet.ell_00199380_PEP_         EDDQQQDASTQEDDYLDRLPKSKKGLQSLLQDIEKRILHYKQLFFKEQN...........
Tet.vor_00155570_PEP_         DDEQQQETAAKEDDYLDRLPKSRKGLSALLESIDKRILHYKQLFFKEQN...........
Tet.pyr_00179540_PEP_         DDEQLQETAAKEDDYLDRLPKSRKGLSALLGDIEKRILHYKQLFFKEQN...........
Tet.can_00119340_PEP_         DDEQQQEMGPKEDDYLDRLPKSRKGLSALLADIDKRIIHYKQLFFKEQN...........
Tet.bor_00147860_PEP_         DDEQQQEMGPKEDDYLDRLPKSRKGLSALLADIDKRILHYKQLFFKEQN...........
Tet.epi_00127660_PEP_         EDEQNLETMAKEDDYLDRLPKSRKGLSAFLQDIEKKILHYKQLFFKEQN...........
Tet.epi_00126830_PEP_         NDELNVETTAKEDDYLDRLPKSRKGLSAFLQDIEKKILHYKQLFFKEQN...........
Tet.par_00146520_PEP_         KEELDEGQQVEEVDYLDRLPKTKKGLQALLQDVERKIFFYKQIFFKEQN...........
Par.bur_P08360024             ...........YIFIDKLPNQETALLKFQEQVRNRIKFYKQRYLQECY...........
Par.bur_P03800026             ...........YIFIDKLPNQETALLKFQEQVRNRIKFYKQRYLQECY...........
Pse.per_1                     SKSYNQEMQADEEEILDKFPKNLRALEKMLEKVNERLDHYKEKYFEEWE...........
Par.bur_P04570033             RDKSESEIDLKETIEYDKLPKNIAALEKLRFKANACVDALKLDWFKEQS...........
Crei_1335048048               ....REALLEQVAKYRGAVRDMASALRSSTLPGGVSEGGLPHADELLVLGRDVAHHAALA
Crei_159474530                ....REALLEQVAKYRGAVRDMASALRSSTLPG...........................
Arob_1183359562               KDEVLKKLIERETELKLKIEKEIEGIKNLELKG......FSALTQKDED...........
Arep_1183482424               YDFDFKKLWLQERGLMERIDGLLKDIARLTDFKGHYRDMVIPSDDEDDL...........

Oxy.tri_42188                 ............................................................
Oxy.tri_21422                 ............................................................
Cam.sin_17781                 ............................................................
Cam.sin_15892                 ............................................................
Tet.the_AMT1                  ............................................................
Tet.mal_00157670_PEP_         ............................................................
Tet.ell_00199380_PEP_         ............................................................
Tet.vor_00155570_PEP_         ............................................................
Tet.pyr_00179540_PEP_         ............................................................
Tet.can_00119340_PEP_         ............................................................
Tet.bor_00147860_PEP_         ............................................................
Tet.epi_00127660_PEP_         ............................................................
Tet.epi_00126830_PEP_         ............................................................
Tet.par_00146520_PEP_         ............................................................
Par.bur_P08360024             ............................................................
Par.bur_P03800026             ............................................................
Pse.per_1                     ............................................................
Par.bur_P04570033             ............................................................
Crei_1335048048               VSAAAAAAAAGGSAGAAGAGPAAADDDAAAAAAADPDAAAAGGEFSHLGEEGVDYTFGNV
Crei_159474530                ............................................................
Arob_1183359562               ...........................................................V
Arep_1183482424               ...........................................................D
```

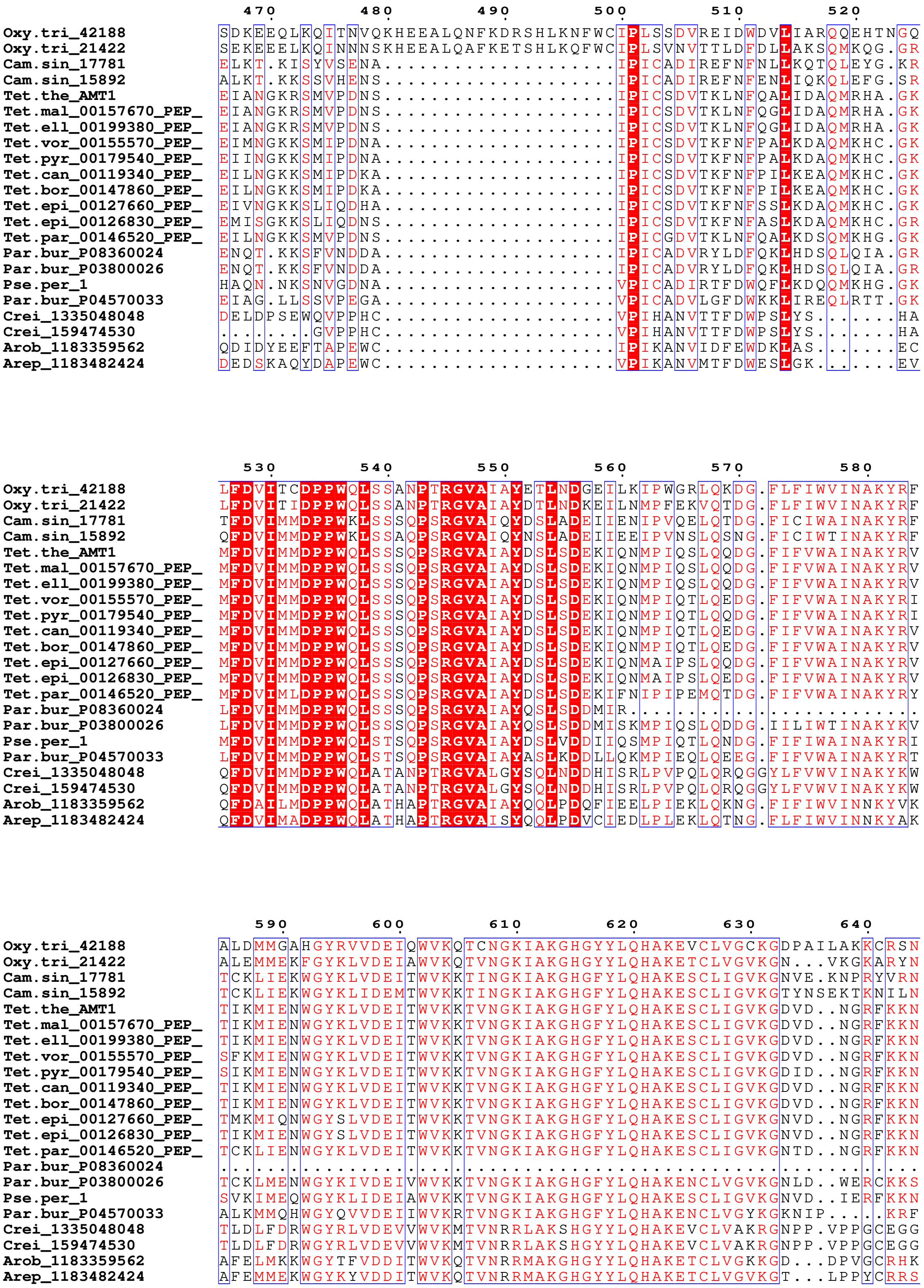
470 480 490 500 510 520
Oxy.tri_42188 SDKEEQLKQITNVQKHEEALQNFKDRSHLKNFWCIPLSSDVREIDWDVLIARQQEHTNGQ
Oxy.tri_21422 SEKEEELKQINNNSKHEEALQAFKETSHLKQFWCIPLSVNVTTLDFDLLAKSQMKQG.GR
Cam.sin_17781 ELKT.KISYVSENA....................IPICADIREFNFNLLKQTQLEYG.KR
Cam.sin_15892 ALKT.KKSSVHENS....................IPICADIREFNFENLIQKQLEFG.SR
Tet.the_AMT1 EIANGKRSMVPDNS....................IPICSDVTKLNFQALIDAQMRHA.GK
Tet.mal_00157670_PEP_ EIANGKRSMVPDNS....................IPICSDVTKLNFQGLIDAQMRHA.GK
Tet.ell_00199380_PEP_ EIANGKRSMVPDNS....................IPICSDVTKLNFQGLIDAQMRHA.GK
Tet.vor_00155570_PEP_ EIMNGKKSMIPDNA....................IPICSDVTKFNFPALKDAQMKHC.GK
Tet.pyr_00179540_PEP_ EIINGKKSMIPDNA....................IPICSDVTKFNFPALKDAQMKHC.GK
Tet.can_00119340_PEP_ EILNGKKSMIPDKA....................IPICSDVTKFNFPILKEAQMKHC.GK
Tet.bor_00147860_PEP_ EILNGKKSMIPDKA....................IPICSDVTKFNFPILKEAQMKHC.GK
Tet.epi_00127660_PEP_ EIVNGKKSLIQDHA....................IPICSDVTKFNFSSLKDAQMKHC.GK
Tet.epi_00126830_PEP_ EMISGKKSLIQDNS....................IPICSDVTKFNFASLKDAQMKHC.GR
Tet.par_00146520_PEP_ EILNGKKSMVPDNS....................IPICGDVTKLNFQALKDSQMKHG.GK
Par.bur_P08360024 ENQT.KKSFVNDDA....................IPICADVRYLDFQKLHDSQLQIA.GR
Par.bur_P03800026 ENQT.KKSFVNDDA....................IPICADVRYLDFQKLHDSQLQIA.GR
Pse.per_1 HAQN.NKSNVGDNA....................VPICADIRTFDWQFLKDQQMKHG.GK
Par.bur_P04570033 EIAG.LLSSVPEGA....................VPICADVLGFDWKKLIREQLRTT.GK
Crei_1335048048 DELDPSEWQVPPHC....................VPIHANVTTFDWPSLYS.......HA
Crei_159474530 ........GVPPHC....................VPIHANVTTFDWPSLYS.......HA
Arob_1183359562 QDIDYEEFTAPEWC....................IPIKANVIDFEWDKLAS.......EC
Arep_1183482424 DEDSKAQYDAPEWC....................VPIKANVMTFDWESLGK.......EV
530 540 550 560 570 580
Oxy.tri_42188 LFDVITCDPPWQLSSANPTRGVAIAYETLNDGEILKIPWGRLQKDG.FLFIWVINAKYRF
Oxy.tri_21422 LFDVITIDPPWQLSSANPTRGVAIAYDTLNDKEILNMPFEKVQTDG.FLFIWVINAKYRF
Cam.sin_17781 TFDVIMMDPPWKLSSSQPSRGVAIQYDSLADEIIENIPVQELQTDG.FICIWAINAKYRF
Cam.sin_15892 QFDVIMMDPPWKLSSAQPSRGVAIQYNSLADEIIEEIPVNSLQSNG.FICIWTINAKYRF
Tet.the_AMT1 MFDVIMMDPPWQLSSSQPSRGVAIAYDSLSDEKIQNMPIQSLQQDG.FIFVWAINAKYRV
Tet.mal_00157670_PEP_ MFDVIMMDPPWQLSSSQPSRGVAIAYDSLSDEKIQNMPIQSLQQDG.FIFVWAINAKYRV
Tet.ell_00199380_PEP_ MFDVIMMDPPWQLSSSQPSRGVAIAYDSLSDEKIQNMPIQSLQQDG.FIFVWAINAKYRV
Tet.vor_00155570_PEP_ MFDVIMMDPPWQLSSSQPSRGVAIAYDSLSDEKIQNMPIQTLQEDG.FIFVWAINAKYRI
Tet.pyr_00179540_PEP_ MFDVIMMDPPWQLSSSQPSRGVAIAYDSLSDEKIQNMPIQTLQQDG.FIFVWAINAKYRI
Tet.can_00119340_PEP_ MFDVIMMDPPWQLSSSQPSRGVAIAYDSLSDEKIQNMPIQTLQEDG.FIFVWAINAKYRV
Tet.bor_00147860_PEP_ MFDVIMMDPPWQLSSSQPSRGVAIAYDSLSDEKIQNMPIQTLQEDG.FIFVWAINAKYRV
Tet.epi_00127660_PEP_ MFDVIMMDPPWQLSSSQPSRGVAIAYDSLSDEKIQNMAIPSLQQDG.FIFVWAINAKYRV
Tet.epi_00126830_PEP_ MFDVIMMDPPWQLSSSQPSRGVAIAYDSLSDEKIQNMAIPSLQEDG.FIFVWAINAKYRV
Tet.par_00146520_PEP_ MFDVIMLDPPWQLSSSQPSRGVAIAYDSLSDEKIFNIPIPEMQTDG.FIFVWAINAKYRY
Par.bur_P08360024 LFDVIMMDPPWQLSSSQPSRGVAIAYQSLSDDMIR.........................
Par.bur_P03800026 LFDVIMMDPPWQLSSSQPSRGVAIAYQSLSDDMISKMPIQSLQDDG.IILIWTINAKYKV
Pse.per_1 MFDVIMMDPPWQLSTSQPSRGVAIAYDSLVDDIIQSMPIQTLQNDG.FIFIWAINAKYRI
Par.bur_P04570033 LFDVIMMDPPWQLSTSQPSRGVAIAYASLKDDLLQKMPIEQLQEEG.FIFVWAINAKYRT
Crei_1335048048 QFDVIMMDPPWQLATANPTRGVALGYSQLNDDHISRLPVPQLQRQGGYLFVWVINAKYKW
Crei_159474530 QFDVIMMDPPWQLATANPTRGVALGYSQLNDDHISRLPVPQLQRQGGYLFVWVINAKYKW
Arob_1183359562 QFDAILMDPPWQLATHAPTRGVAIAYQQLPDQFIEELPIEKLQKNG.FIFIWVINNKYVK
Arep_1183482424 QFDVIMADPPWQLATHAPTRGVAISYQQLPDVCIEDLPLEKLQTNG.FLFIWVINNKYAK
590 600 610 620 630 640
Oxy.tri_42188 ALDMMGAHGYRVVDEIQWVKQTCNGKIAKGHGYYLQHAKEVCLVGCKGDPAILAKKCRSN
Oxy.tri_21422 ALEMMEKFGYKLVDEIAWVKQTVNGKIAKGHGYYLQHAKETCLVGVKGN...VKGKARYN
Cam.sin_17781 TCKLIEKWGYKLVDEITWVKKTINGKIAKGHGFYLQHAKESCLIGVKGNVE.KNPRYVRN
Cam.sin_15892 TCKLIEKWGYKLIDEMTWVKKTINGKIAKGHGFYLQHAKESCLIGVKGTYNSEKTKNILN
Tet.the_AMT1 TIKMIENWGYKLVDEITWVKKTVNGKIAKGHGFYLQHAKESCLIGVKGDVD..NGRFKKN
Tet.mal_00157670_PEP_ TIKMIENWGYKLVDEITWVKKTVNGKIAKGHGFYLQHAKESCLIGVKGDVD..NGRFKKN
Tet.ell_00199380_PEP_ TIKMIENWGYKLVDEITWVKKTVNGKIAKGHGFYLQHAKESCLIGVKGDVD..NGRFKKN
Tet.vor_00155570_PEP_ SFKMIENWGYKLVDEITWVKKTVNGKIAKGHGFYLQHAKESCLIGVKGDVD..NGRFKKN
Tet.pyr_00179540_PEP_ SIKMIENWGYKLVDEITWVKKTVNGKIAKGHGFYLQHAKESCLIGVKGDID..NGRFKKN
Tet.can_00119340_PEP_ TIKMIENWGYKLVDEITWVKKTVNGKIAKGHGFYLQHAKESCLIGVKGDVD..NGRFKKN
Tet.bor_00147860_PEP_ TIKMIENWGYKLVDEITWVKKTVNGKIAKGHGFYLQHAKESCLIGVKGDVD..NGRFKKN
Tet.epi_00127660_PEP_ TMKMIQNWGYSLVDEITWVKKTVNGKIAKGHGFYLQHAKESCLIGVKGNVD..NGRFKKN
Tet.epi_00126830_PEP_ TIKMIENWGYSLVDEITWVKKTVNGKIAKGHGFYLQHAKESCLIGVKGNVD..NGRFKKN
Tet.par_00146520_PEP_ TCKLIENWGYKLVDEITWVKKTVNGKIAKGHGFYLQHAKESCLIGVKGNTD..NGRFKKN
Par.bur_P08360024 ............................................................
Par.bur_P03800026 TCKLMENWGYKIVDEIVWVKKTVNGKIAKGHGFYLQHAKENCLVGVKGNLD..WERCKKS
Pse.per_1 SVKIMEQWGYKLIDEIAWVKKTVNGKIAKGHGFYLQHAKESCLIGVKGNVD..IERFKKN
Par.bur_P04570033 ALKMMQHWGYQVVDEIIWVKRTVNGKIAKGHGFYLQHAKENCLVGYKGKNIP.....KRF
Crei_1335048048 TLDLFDRWGYRLVDEVVWVKMTVNRRLAKSHGYYLQHAKEVCLVAKRGNPP.VPPGCEGG
Crei_159474530 TLDLFDRWGYRLVDEVVWVKMTVNRRLAKSHGYYLQHAKEVCLVAKRGNPP.VPPGCEGG
Arob_1183359562 AFELMKKWGYTFVDDITWVKQTVNRRMAKGHGYYLQHAKETCLVGKKGD...DPVGCRHK
Arep_1183482424 AFEMMEKWGYKYVDDITWVKQTVNRRMAKGHGYYLQHAKETCLVGVKGT...LPPYCRRS

```
                               650       660       670       680       690
Oxy.tri_42188                  IESDVIFSERRGQSQKPEEIYELVEALVPNGYYMEIFGRRNNLHNGWVTVGNEL......
Oxy.tri_21422                  IESDVIFSQRRGQSQKPEEIYEIAEALVPNGYYLEIFGRRNNLHNGWVTIGNEL......
Cam.sin_17781                  VANDVIFSERRGQSQKPTEVYDYLEKLVPHGFYLEIFGRRNNLRKNWVTIGNEI......
Cam.sin_15892                  VASDVIFSERRGQSQKPTEMYEYIEKLVPNGFYLEIFGRRNNLRGNWVTIGNEI......
Tet.the_AMT1                   IASDVIFSERRGQSQKPEEIYQYINQLCPNGNYLEIFARRNNLHDNWVSIGNEL......
Tet.mal_00157670_PEP_          IASDVIFSERRGQSQKPEEIYQYINQLCPNGNYLEIFARRNNLHDNWVSIGNEL......
Tet.ell_00199380_PEP_          IASDVIFSERRGQSQKPEEIYQYINQLCPNGNYLEIFARRNNLHDNWVSIGNEL......
Tet.vor_00155570_PEP_          IASDVIFSERRGQSQKPEEIYQYIQQLCPNGHYLEVFARRNNLHDNWVSIGNEL......
Tet.pyr_00179540_PEP_          IASDVIFSERRGQSQKPEEIYQYIQQLSPNGHFLEVFARRNNLHDNWVSIGNEL......
Tet.can_00119340_PEP_          IASDVIFSERRGQSQKPEEIYQYIQQLCPNGQFLEIFARRNNLHDNWVSIGNEL......
Tet.bor_00147860_PEP_          IASDVIFSERRGQSQKPEEIYQYIQQLCPNGQFLEIFARRNNLHDNWVSIGNEL......
Tet.epi_00127660_PEP_          ISSDVIFSERRGQSQKPEEIYQYIQQLCPNGHFLEVFARRNNLHSNWVSIGNEL......
Tet.epi_00126830_PEP_          IVSDVIFSERRGQSQKPEEIYQYIQQLCPNGHFLEVFARRNNLHSNWVSIGNEL......
Tet.par_00146520_PEP_          IASDVIFSERRGQSQKPEEIYTYINELCPNGHYLEIFARRNNLHDNWVSIGNEL......
Par.bur_P08360024              ............................................................
Par.bur_P03800026              VASDVIFSERRGQSQKPEEIYKYVEELIPNGHYLEIFGRRNNIRKNWVTIGNEL......
Pse.per_1                      IASDIIFSERRGQSQKPEEIYEYAEQLVPNGFYLEVFGRRNNLRSNWITIGNEL......
Par.bur_P04570033              IGSDIIWSERRGQSQKPQEIYDLIEQLVPNGYYLEIFGRRNNLRDKWVTIGNEI......
Crei_1335048048                VGSDIIFSERRGQSQKPEEIYHLIEQLVPNGRYLEIFARKNNLRNYWVSIGNEVTGTGLP
Crei_159474530                 VGSDIIFSERRGQSQKPEEIYHLIEQLVPNGRYLEIFARKNNLRNYWVSIGNEVTGTGLP
Arob_1183359562                ISSDVIYSVRRGQSQKPEELYEMIEELIPNGKYLEIFGRKNNLRDYWVTIGNEL......
Arep_1183482424                VGSDVIYSERRGQSQKPEQIYELIEEMMPGGKYLEIFGRKNNLRDYWITVGNEL......

Oxy.tri_42188                  ............................................................
Oxy.tri_21422                  ............................................................
Cam.sin_17781                  ............................................................
Cam.sin_15892                  ............................................................
Tet.the_AMT1                   ............................................................
Tet.mal_00157670_PEP_          ............................................................
Tet.ell_00199380_PEP_          ............................................................
Tet.vor_00155570_PEP_          ............................................................
Tet.pyr_00179540_PEP_          ............................................................
Tet.can_00119340_PEP_          ............................................................
Tet.bor_00147860_PEP_          ............................................................
Tet.epi_00127660_PEP_          ............................................................
Tet.epi_00126830_PEP_          ............................................................
Tet.par_00146520_PEP_          ............................................................
Par.bur_P08360024              ............................................................
Par.bur_P03800026              ............................................................
Pse.per_1                      ............................................................
Par.bur_P04570033              ............................................................
Crei_1335048048                DEDMQALRDLHHIPGAVYGKNGGGGGGPSNGGAAPPASRGPSGGGAAPPAAARAAAAKPP
Crei_159474530                 DEDMQALRDLHHIPGAVYGKN.......................................
Arob_1183359562                ............................................................
Arep_1183482424                ............................................................

Oxy.tri_42188                  ............................................................
Oxy.tri_21422                  ............................................................
Cam.sin_17781                  ............................................................
Cam.sin_15892                  ............................................................
Tet.the_AMT1                   ............................................................
Tet.mal_00157670_PEP_          ............................................................
Tet.ell_00199380_PEP_          ............................................................
Tet.vor_00155570_PEP_          ............................................................
Tet.pyr_00179540_PEP_          ............................................................
Tet.can_00119340_PEP_          ............................................................
Tet.bor_00147860_PEP_          ............................................................
Tet.epi_00127660_PEP_          ............................................................
Tet.epi_00126830_PEP_          ............................................................
Tet.par_00146520_PEP_          ............................................................
Par.bur_P08360024              ............................................................
Par.bur_P03800026              ............................................................
Pse.per_1                      ............................................................
Par.bur_P04570033              ............................................................
Crei_1335048048                LPLGPTAAAAAAASAAASTLAEAAAAAAAAGADGEVQQQDVDLPMPQAEDAGDALPTPLA
Crei_159474530                 ...APHLVSKLFLYAPNSSREEG.....................................
Arob_1183359562                ............................................................
Arep_1183482424                ............................................................
```

```
Oxy.tri_42188              ............................................................
Oxy.tri_21422              ............................................................
Cam.sin_17781              ............................................................
Cam.sin_15892              ............................................................
Tet.the_AMT1               ............................................................
Tet.mal_00157670_PEP_      ............................................................
Tet.ell_00199380_PEP_      ............................................................
Tet.vor_00155570_PEP_      ............................................................
Tet.pyr_00179540_PEP_      ............................................................
Tet.can_00119340_PEP_      ............................................................
Tet.bor_00147860_PEP_      ............................................................
Tet.epi_00127660_PEP_      ............................................................
Tet.epi_00126830_PEP_      ............................................................
Tet.par_00146520_PEP_      ............................................................
Par.bur_P08360024          ............................................................
Par.bur_P03800026          ............................................................
Pse.per_1                  ............................................................
Par.bur_P04570033          ............................................................
Crei_1335048048            APPPAEVAAAAQRQGAGGAGAGPGAGEPEDVSIGLGSLPSLGMLESALRSGGVGVCGGGG
Crei_159474530             ............................................................
Arob_1183359562            ............................................................
Arep_1183482424            ............................................................
```

```
Oxy.tri_42188              ....................................
Oxy.tri_21422              ....................................
Cam.sin_17781              ....................................
Cam.sin_15892              ....................................
Tet.the_AMT1               ....................................
Tet.mal_00157670_PEP_      ....................................
Tet.ell_00199380_PEP_      ....................................
Tet.vor_00155570_PEP_      ....................................
Tet.pyr_00179540_PEP_      ....................................
Tet.can_00119340_PEP_      ....................................
Tet.bor_00147860_PEP_      ....................................
Tet.epi_00127660_PEP_      ....................................
Tet.epi_00126830_PEP_      ....................................
Tet.par_00146520_PEP_      ....................................
Par.bur_P08360024          ....................................
Par.bur_P03800026          ....................................
Pse.per_1                  ....................................
Par.bur_P04570033          ....................................
Crei_1335048048            GALETCARGGGGGGGGGRPSVGPGLSGLSAGPLRFD
Crei_159474530             ....................................
Arob_1183359562            ....................................
Arep_1183482424            ....................................
```
