## Supplemental file S2 for "Structural basis of DNA N^6^-adenine methylation in eukaryotes"

```
Oxy.tri_794                   ............................................................
Oxy.tri_24932                 ............................................................
Oxy.tri_799                   ............................................................
Cam.sin_964                   ............................................................
Vor.cam_9330                  ............................................................
Par.bur_P04390037             ............................................................
Par.bur_P09660035             ............................................................
Par.tet_001442122.1           ............................................................
Tet.mal_00222350_PEP_         ............................................................
Tet.the_AMT6                  ............................................................
Tet.ell_00078280_PEP_         ............................................................
Tet.vor_00219940_PEP_         ............................................................
Tet.bor_00193820_PEP_         ............................................................
Tet.can_00194390_PEP_         ............................................................
Tet.pyr_00246380_PEP_         ............................................................
Tet.epi_00174830_PEP_         ............................................................
Tet.par_00207750_PEP_         ............................................................
Ich.mul_156080                ............................................................
Par.bur_P01400001             ............................................................
Par.bur_P02340001             ............................................................
Par.tet_GSPATP00024232001     ............................................................
Par.tet_GSPATP00005554001     ............................................................
Vor.cam_9610                  ............................................................
Vor.cam_12567                 ..........................................MGISGLEFQQLNLSPLTI
Vor.cam_9611                  ............................................................
Tet.the_AMT7                  ............................................................
Tet.mal_00069950_PEP_         ............................................................
Tet.ell_00008460_PEP_         ............................................................
Tet.pyr_00085530_PEP_         MVINNIICSDININNIILRHQKNRNRNQRGQVREHHPRSNAVTFAIWHKVQVRSFESHFG
Tet.vor_00057090_PEP_         ............................................................
Tet.bor_00075080_PEP_         ............................................................
Tet.can_00083510_PEP_         ............................................................
Tet.epi_00101720_PEP_         ............................................................
TEPIDO00091130_PEP_           ............................................................
Tet.par_00104560_PEP_         ............................................................
Ich.mul_AMT7                  ............................................................
Cam.sin_20221                 ............................................................
Cam.sin_20222                 ............................................................
Cam.sin_7583                  ............................................................
Oxy.tri_21422                 ...........MESTKNARGNSRDHNNSGASGADAATNSKKKNQPALTQFFKKQPEGAIP
Oxy.tri_42188                 ...........MNQSSQDITTQKSSNGFNPQTQPETLIQVIRKESTFIFKYRKNPYYVPP
Oxy.tri_11745                 ..................MSEKKPARKQKKSETSPQAQPPANSGGNLEANSGDNSNQLKK
Oxy.tri_31967                 ..........MQRNTRSKSPSKHNSDNFEGQLNLKKIVLRKREQRSTTMQGKNYQNFYKR
```

```
                                                         1       10        20        30
Oxy.tri_794                   ...........................MELQQKIQTKSLSKKRFNLRDKKAGKNHKLGDL
Oxy.tri_24932                 ...........................MELQQKIQTKSLSKKRFNLRDKKAGKNHKLGDL
Oxy.tri_799                   ...........................MELQQKIQTKSLSKKRFNLRDKKAGKNHKLGDL
Cam.sin_964                   ............................................................
Vor.cam_9330                  ............................................................
Par.bur_P04390037             ............................................................
Par.bur_P09660035             ............................................................
Par.tet_001442122.1           ............................................................
Tet.mal_00222350_PEP_         ...........................................................M
Tet.the_AMT6                  ............................................................
Tet.ell_00078280_PEP_         ...........................................................M
Tet.vor_00219940_PEP_         ...........................................................M
Tet.bor_00193820_PEP_         ............................................................
Tet.can_00194390_PEP_         ............................................................
Tet.pyr_00246380_PEP_         ............................................................
Tet.epi_00174830_PEP_         .................MKQENIEKTSKTQIKQIDRVKIRKNREKKARKQSGKIIQIPQI
Tet.par_00207750_PEP_         ............................................................
Ich.mul_156080                ............................................................
Par.bur_P01400001             ...............................MQKLKIQKKQVDIRDRLRSRVPGSDSRKV
Par.bur_P02340001             ...............................MQKLKIQKKQVDIRDRLRSRVPGSDSRKV
Par.tet_GSPATP00024232001     ...................................MGKNQRESLRMRLRYNEPGSENKRI
Par.tet_GSPATP00005554001     .............................................MKLRYSEPGSENKRI
Vor.cam_9610                  ............................MKSASKKNRQVVLGESQEQVRRSARERKEPKN
Vor.cam_12567                 NEEELLIATQPQWEDTSIISFFPFFHILEKSASKKNRQVVLGESQEQVRRSARERKEPKN
Vor.cam_9611                  ...................................................MFGYIDFRN
Tet.the_AMT7                  ......................................MAPKKQEQEPIRLSTRTASKKV
Tet.mal_00069950_PEP_         ......................................MAPKKQEQEPIRLSTRTASKKV
Tet.ell_00008460_PEP_         ......................................MAPKKQEQEPIRLSTRTASKKV
Tet.pyr_00085530_PEP_         ENKKGKKQVKKHSQSHCFFLYRYGAIKRNYIVIISKFRIDTTALPETEQNLTSFSSINRV
Tet.vor_00057090_PEP_         ................................................MRLSTRVASKKV
Tet.bor_00075080_PEP_         ......................................MAPRKQEQEPKRLSTRTSSKKV
Tet.can_00083510_PEP_         ......................................MAPRKQEQEPKRLSTRTSSKKV
Tet.epi_00101720_PEP_         ......................................MGLKKQEKVIKRSSIRTSSKKV
TEPIDO00091130_PEP_           ......................................MAPRKKEQEVKRLSIRTSSKKV
Tet.par_00104560_PEP_         .........................................MIYEIYKMYLSTPDEMKVR
Ich.mul_AMT7                  ............................................................
Cam.sin_20221                 ..................................MKTKNQPILTEVPLRRSYRERTLKKY
Cam.sin_20222                 ..................................MKTKNQPILTEVPLRRSYRERTLKKY
Cam.sin_7583                  ...............................................MFCFVNFLNIKRI
Oxy.tri_21422                 KGS.........EANNRTSEQSNSNASANANHHSEHVQTRQSAQLSQQNRDSQNTKVNQI
Oxy.tri_42188                 PISSQTSPNLEVETSNDLNQMSDYEGQIPNNYEINRNSTQFTNNDDQSDNDFYDNNSITT
Oxy.tri_11745                 QKQNLALSNNGNSLDKICSHLKPQAAGLKTSGQGVSQQQQHQSISSSTGAASVKTLTQSS
Oxy.tri_31967                 LDDGLDSGTAQGQRKQSIDSDNIKKPKDRKKRKSKPIKKNKTQLRTLSRKSILQLSKISA
```

```
                                         40        50        60        70        80        90
Oxy.tri_794                  VVEEEDKFENAEESSATLNTESQEGKKPNQQQTNGRVTRNKAPQDYKKMFHPDVFSDDDD
Oxy.tri_24932                VVEEEDKFENAEESSATLNTESQEGKKPNQQQTNGRVTRNKAPQDYKKMFHPDVFSDDDD
Oxy.tri_799                  VVEEEDKFENAEESSASLNTESQEGKKPNQQQTNGRVTRNKAPQDYKKMFHPDVFSDDDD
Cam.sin_964                  .MFEKNPEEEKIEP................................LKRKAEYILDGYKD
Vor.cam_9330                 .MTESSTLTTKANP................................LMNLFLKREAEHPL
Par.bur_P04390037            ............................................................
Par.bur_P09660035            ............................................................
Par.tet_001442122.1          ............................................................
Tet.mal_00222350_PEP_        KSQETLA...ACQSLDK...........................FAHPKKVSP.VQKSQI
Tet.the_AMT6                 MSQETLA...ACQSLDK...........................FAHPKKVSP.VQKSQI
Tet.ell_00078280_PEP_        KSQETLA...ACQSLEK...........................FAHPKKPSP.VQKSQI
Tet.vor_00219940_PEP_        KSQATLA...ACKTLEK...........................FALPKKTSP.VQKN..
Tet.bor_00193820_PEP_        MSEETLA...ACLLLD...........................KFALPKKTSP.IQKHQI
Tet.can_00194390_PEP_        MSEATLA...ACLSLDKQTYFNKNLQKCQQVDFKHLFIFINIKRFALPKKTSP.IQKHQI
Tet.pyr_00246380_PEP_        ......................................................MQKHQI
Tet.epi_00174830_PEP_        MSEATVT...ACQSLEK...........................FALPKKASPSHKQQMI
Tet.par_00207750_PEP_        ............................................................
Ich.mul_156080               ............................................................
Par.bur_P01400001            NYFKMNQGYDNEQDESEYIDDREQQQRIKR......................IRRFHREK
Par.bur_P02340001            NYFKMNQGYDNEQEESEYVDDREQQQRIKR......................IRRFHREK
Par.tet_GSPATP00024232001    NYFKLNQGYDITEDDSEYMDEQDSYKAFKK......................IRKYKTKS
Par.tet_GSPATP00005554001    NYFKLNQGYDVTEDDSEYVDDQDSYKAFKK......................IRKYKTKS
Vor.cam_9610                 DDYFMGSSYIEDIFGPSYNEDEPSPKRGRR......................NYRYDDSD
Vor.cam_12567                DDYFMGSSYIEDIFGPSYNEDEPSPKRGRR......................NYRYDDSD
Vor.cam_9611                 DDYFMGSSYIEDIFGPSYNEDEPSPKRGRR......................NYRYDDSD
Tet.the_AMT7                 DYLQLSNGKLEDFFDDLEEDNKPARNRSRSKKRGRKPLKKADSR.SKTPSRVSNARGRSK
Tet.mal_00069950_PEP_        DYLQLSNGKLEDFFDDLEEDNKPARNRSRSKKRGRKPLKKADSR.SKTPSRASSVRGRSK
Tet.ell_00008460_PEP_        DYLQLSNGKLEDFFDDLEEDNKPARNRSRSKKRGRKPLKKADSR.SKTPSRASCGRGRSK
Tet.pyr_00085530_PEP_        DYLQLSNGKLDGFFDELEEDNKPARSRTRSKKRGRKPKAKVEAR.SKTPSRAAPKRGRSK
Tet.vor_00057090_PEP_        DYLQLSNGKLDGFFDDLEEDNKPARTRSRSKKRGRKPKINIKVARSKTPVRAPAKRGRSK
Tet.bor_00075080_PEP_        DYLQLSNGKLDGFFEELEEDNKPARTRSRSKKRGRKPKRKTDGR.SRTPVRAPARSSRSK
Tet.can_00083510_PEP_        DYLQLSNGKLDGFFEELEEDNKPARTRSRSKKRGRKPKRKTDGR.SRTPVRAPARSSRSK
Tet.epi_00101720_PEP_        DYLQLSNGKLDGYFDELEENNKPARQRSRSKKRGRKPKVSVTER.SRTPSRVTSRKPRSK
TEPIDO00091130_PEP_          DYLQLSNGKLDDFFDELEEDNKPARNRSRSKKRGRKPKAMVIAR.SSTPSSATSRRSRSK
Tet.par_00104560_PEP_        PTLYLNLTAITSKFNRRPCLIRNRETFIENSKIKKRRGLAVDLLQKRLIICNSLTGSLKA
Ich.mul_AMT7                 ............................................................
Cam.sin_20221                DYLTISNGRYSEFEIDLFEDEEYEPKKGRRHRPRKD................SEDFDNSQ
Cam.sin_20222                DYLTISNGRYSEFEIDLFEDEEYEPKKGRRHRPRKD................SEDFDNSQ
Cam.sin_7583                 NFPAIES..IFYLKLTQLKMDDYEPKREKKRKHRKD................SEDFDNSE
Oxy.tri_21422                NQNGATGVNSLLQTPKKKLLETPSKKKTYADE...........PDEIDMREIQSLVDKKV
Oxy.tri_42188                MQIDTSTAKILNNGPLEYNPDLPNKEQKLKDSQVMQNQPPTATSTNSQQRTLQELINIMP
Oxy.tri_11745                GEESLSQSQLGKRKERGEFLGNTNSNNSQNQQQTRPNLRRQANANAIPESNEQRRDSSVR
Oxy.tri_31967                KKNKMNQSDEFKNVKSVKESASNFASDKLLNSNGFAEKQEIVLQSSGPIQLEASAININN
```

```
                                   100       110       120       130
Oxy.tri_794                  FVPKNKGKAKKRDLKRKLKVKKGGRFNNYENIGAQTKG......................
Oxy.tri_24932                FVPKNKGKAKKRDLKRKLKVKKGGRFNNYENIGAQTKG......................
Oxy.tri_799                  FVPKNKGKAKKRDLKRKLKVKKGGRFNNYENIGAQTKG......................
Cam.sin_964                  LIKKVKTTEEPKSSGENNSLAHLLKWRDEIKKINYELE......................
Vor.cam_9330                 PLKKPVKKIKPNDEKNTDPLYQILSWRDTIKMPPLPHP......................
Par.bur_P04390037            .........MKSNKEQLNRLRMVLEYRQNPKYQPQIAE......................
Par.bur_P09660035            .........MKSNKEQLNRLRMVLEYRQNPKYQPQIAE......................
Par.tet_001442122.1          ...MNIKKPTTKSKEDQNRLRMVLDYRWDPKYQPQGKP......................
Tet.mal_00222350_PEP_        IEEPPLQKKIKQSEPGEDQLSLLLKWRSSYIPPQKPTN......................
Tet.the_AMT6                 IEEPPLQKKIKPTEPGEDQLSLLLKWRSSYIPPQKPTN......................
Tet.ell_00078280_PEP_        IEEPPLQKKVKPSEPGEDQLSLLLKWRAQYIPPQKPTN......................
Tet.vor_00219940_PEP_        ............................................................
Tet.bor_00193820_PEP_        IEEPSLQKKLKPNDPQQDQLSLLLKWRNQYIPPAKPTN......................
Tet.can_00194390_PEP_        IEEPSLQKKLKPNDPQQDQLSLLLKWRNQYIPPAKPTN......................
Tet.pyr_00246380_PEP_        IEEPPLKKQQKPNEPGQDQLSLLLKWRNEYIPPITPKN......................
Tet.epi_00174830_PEP_        IDEPSLQKKLKPNDPQQDQLSLLLRWRNEYIPPTKTLN......................
Tet.par_00207750_PEP_        .............................MKVPEKPTS......................
Ich.mul_156080               ............................................................
Par.bur_P01400001            KKS............LNDNLHVLLGWKKDIQLKD..........................
Par.bur_P02340001            KKS............LNDNLQILVGWKKDIQLKD..........................
Par.tet_GSPATP00024232001    KED..........KVTCDNLEVLLQWKGSINCQS..........................
Par.tet_GSPATP00005554001    KKEK........MYIMLANQNGVIIWKCCYNGKNQLSANLM...................
Vor.cam_9610                 DYSEGHKYSKKNKQINTNKVEALLDWKLQLQALPIN.LNYL...................
Vor.cam_12567                DYSEGHKYSKKNKQINTNKVEALLDWKLQLQALPIN.LNYL...................
Vor.cam_9611                 DYSEGHKYSKKNKQINTNKVEALLDWKLQLQALPIN.LNYL...................
Tet.the_AMT7                 SLGPRKTYPRKKNLSPDNQLSLLLKWRNDKIPLKSASE......................
Tet.mal_00069950_PEP_        SLGPRKTYPRKKNLSPDNQLSLLLKWRNDKIPQKSASE......................
Tet.ell_00008460_PEP_        SLGPRKTYPRKKNLSPDNQLSLLLKWRNEKIPLKSASE......................
Tet.pyr_00085530_PEP_        SQTVRRVYPRKKNLSPDNQLSLLLKWRNEKVPLEPASD......................
Tet.vor_00057090_PEP_        TQTVRRVYPRKKNLSPDNQLSLLLKWRNEKVPLQPASD......................
Tet.bor_00075080_PEP_        SQSIRRVYPRKKNLSPDNQLSLLLKWRNEKVPLKPASE......................
Tet.can_00083510_PEP_        SQSIRRVYPRKKNLSPDNQLSLLLKWRNEKVPLKPASE......................
Tet.epi_00101720_PEP_        QQTVRREYPRKKILSPDNQLSLLLKWRKEKVPMKPGVQ......................
TEPIDO00091130_PEP_          SQTVRRVYPRKKILSPDNQLSLLLKWRNEKIPCEPAQQ......................
Tet.par_00104560_PEP_        SLTTP.......KRSPDNQISLLIKWRQEKK..VPSLS......................
Ich.mul_AMT7                 ............................................................
Cam.sin_20221                KKHRKK.......FSSSSRISYLLEWK..SAALLPETP......................
Cam.sin_20222                KKHRKKECIKLLRFSSSSRISYLLEWK..SAALLPETP......................
Cam.sin_7583                 KKYRKK.......MMLTNRISHLLEWK..QNSLLPEPP......................
Oxy.tri_21422                KESAAAQQQLSQSGIEDSAIKRSLRPRKVENYKNMLEG.........DEITLKTIQD...
Oxy.tri_42188                SIEDISQQCKQQQQLKIQAKANSTQSASTANAANGGKGRKRGRTVRFDQPLLGKVRQRNG
Oxy.tri_11745                ESEQPQFYSDQTQLEGQTQITSFFGGGPGATQSTQGGQMMG...................
Oxy.tri_31967                FFEVIKYYNANRTVKCSERIYEEGLDFLEATFLKQNQSLGLKNQKFDLDAVDLPFPEQ..
```

```
Oxy.tri_794                  ............................................................
Oxy.tri_24932                ............................................................
Oxy.tri_799                  ............................................................
Cam.sin_964                  ............................................................
Vor.cam_9330                 ............................................................
Par.bur_P04390037            ............................................................
Par.bur_P09660035            ............................................................
Par.tet_001442122.1          ............................................................
Tet.mal_00222350_PEP_        ............................................................
Tet.the_AMT6                 ............................................................
Tet.ell_00078280_PEP_        ............................................................
Tet.vor_00219940_PEP_        ............................................................
Tet.bor_00193820_PEP_        ............................................................
Tet.can_00194390_PEP_        ............................................................
Tet.pyr_00246380_PEP_        ............................................................
Tet.epi_00174830_PEP_        ............................................................
Tet.par_00207750_PEP_        ............................................................
Ich.mul_156080               ............................................................
Par.bur_P01400001            ............................................................
Par.bur_P02340001            ............................................................
Par.tet_GSPATP00024232001    ............................................................
Par.tet_GSPATP00005554001    ............................................................
Vor.cam_9610                 ............................................................
Vor.cam_12567                ............................................................
Vor.cam_9611                 ............................................................
Tet.the_AMT7                 ............................................................
Tet.mal_00069950_PEP_        ............................................................
Tet.ell_00008460_PEP_        ............................................................
Tet.pyr_00085530_PEP_        ............................................................
Tet.vor_00057090_PEP_        ............................................................
Tet.bor_00075080_PEP_        ............................................................
Tet.can_00083510_PEP_        ............................................................
Tet.epi_00101720_PEP_        ............................................................
TEPIDO00091130_PEP_          ............................................................
Tet.par_00104560_PEP_        ............................................................
Ich.mul_AMT7                 ............................................................
Cam.sin_20221                ............................................................
Cam.sin_20222                ............................................................
Cam.sin_7583                 ............................................................
Oxy.tri_21422                ...............................EQIEVKRKKREASSQNR............
Oxy.tri_42188                DASDDEEPDEIEMLIRRLHTDILNDARNDPVEQAKKIRQARESQSDQTNSTTQLSVYERM
Oxy.tri_11745                ............................................................
Oxy.tri_31967                ............................................................

                                                          140       150       160
Oxy.tri_794                  .......................QKIGERKVLDGVSIFDLIIPWKEKISGQPNQIGSMKK
Oxy.tri_24932                .......................QKIGERKVLDGVSIFDLIIPWKEKISGQPNQIGSMKK
Oxy.tri_799                  .......................QKTGERKVLDGVSIFDLIIPWKEKISGQPNQIGSMKK
Cam.sin_964                  .......................SGSYKEIVVESILEADLNLHAENLQAIYLNIDWKNEN
Vor.cam_9330                 .......................S.SCAAHHLPSITHQNLEGLGTSLQGIFINIDWDLTP
Par.bur_P04390037            .......................LAPEQYKIVKQANPLEAQS..PDIQALFINIKWNHQQ
Par.bur_P09660035            .......................LAPEQYKIVKQANPLEAQS..PDIQALFINIKWNHEQ
Par.tet_001442122.1          .......................AIPESFKIVRESNPIETKLG.NDLQAIFINLKWG..Q
Tet.mal_00222350_PEP_        .......................EDEYKKVICKDISSEKLEQHAGDVSALFINIKWKLSE
Tet.the_AMT6                 .......................EDEYKKIICKDISSEKLEQHAGDVSALFINIKWKLSE
Tet.ell_00078280_PEP_        .......................EDEYKKIVCQNISSEKLEQHAGDISALFINIKWKLSD
Tet.vor_00219940_PEP_        ............................................................
Tet.bor_00193820_PEP_        .......................TEEYKLISTKEIASEQLDKHAGDLSALFINIQWKLAE
Tet.can_00194390_PEP_        .......................TEEYKLISTKEIASEQLDKHAGDLSALFINIQWKLVE
Tet.pyr_00246380_PEP_        .......................ADEFQQITSTDIANEKLEKYGGDLSALFININWKLKE
Tet.epi_00174830_PEP_        .......................PDQFKSIKTKDIASEYLEKHAGDLSALFINISWKLDD
Tet.par_00207750_PEP_        .......................ESEYKLVKSKDILVEDLSKHGNNLQALFINISWKLSE
Ich.mul_156080               ............................................................
Par.bur_P01400001            .....................PETDNSKIFKVKKILRTDLSKYCQGVQGIFINSVFKN..
Par.bur_P02340001            .....................PERDNSKIFNVKKILRTDLSKYCQGVQGIFINQVFRN..
Par.tet_GSPATP00024232001    .....................YETDQKKIVVLKKILGTDLSQYAKGVQGIFIDNLFKK..
Par.tet_GSPATP00005554001    ..............KQIKEGNHQLFKLRIIVLKKILGTDLSQYVKGVQGIFIDNLFRK..
Vor.cam_9610                 ..............QEETQEVLLHEFHTAKLTKNMGSENLAALGQDIQAIFINIDWGKK.
Vor.cam_12567                ..............QEETQEVLLHEFHTAKLTKNMGSENLAALGQDIQAIFINIDWGKK.
Vor.cam_9611                 ..............QEETQEVLLHEFHTAKLTKNMGSENLAALGQDIQAIFINIDWGKK.
Tet.the_AMT7                 ......................TDNKCKVVNVKNIFKSDLSKYGANLQALFINALWKVKS
Tet.mal_00069950_PEP_        ......................TDNTCKVVNVKNIFKSDLSKYGSNLQALFINALWKVKS
Tet.ell_00008460_PEP_        ......................TDNKCKAVNIKNIFKSDLSKYGSNLQALFINALWKVKS
Tet.pyr_00085530_PEP_        ......................TDNTSKVFSSKNIFKSDLSKYGQNLQALFINALWKVKS
Tet.vor_00057090_PEP_        ......................TDNTSKIFNVKNIFKSDLSKYGQNLQALFINALWKVKS
Tet.bor_00075080_PEP_        ......................SDNNSKVVNVKNIFKSDLSKYGTNLQALFINALWKVKS
Tet.can_00083510_PEP_        ......................SDNNSKVVNVKNIFKSDLSKYGTNLQALFINALWKVKS
Tet.epi_00101720_PEP_        ......................NNNINKVVNVKNIFKADLSKYGSNLQALYINALWKVKS
TEPIDO00091130_PEP_          ......................SENINKVVNVKNIFKTDLSKYGSNLQALFINVLWKVKS
Tet.par_00104560_PEP_        ......................SINDTKVITVKSVVKSDISKYAQNIQALFINVNWAIKS
Ich.mul_AMT7                 ...........................RVIQVKNILKQDLSKYSHDLQALFINILWKVDT
Cam.sin_20221                ......................VSKFCNIHNVKSIINTDLSKYGSHIEAIFINIEWENE.
Cam.sin_20222                ......................VSKFCNIHNVKSIINTDLSKYGSHIEAIFINIEWENE.
Cam.sin_7583                 ......................QNKESRMHQVKSILNTDLSKYGTQIEALFINIQWQQD.
Oxy.tri_21422                .......................LEDEDEDEDMLEVGQQIERASDDEDDDDFPISTRRS.
Oxy.tri_42188                ILGSASQQSTDHQPGEFSNMFRTLEDEQIEINQNFLFDEYDSEDDSIADDKVEIASDDEQ
Oxy.tri_11745                .................GVNPGSPQQQLIQSNKKQYMFKAKRHPKSLMKEGVSTLRYVEG
Oxy.tri_31967                ......PKTKVIAEEILQQQELEQLQQKQINGQQVALLDEAAQHSLKSETFINCDLRFFN
```

```
                               170       180       190       200       210       220
Oxy.tri_794                    QPSNASSTTQTSLSSFLSKNKGGMFINYYNSRILEAANKADQVNGINGANSKVKSQVNDL
Oxy.tri_24932                  QPSNASSTTQTSLSSFLSKNKGGMFINYYNSRILEAANKADQVNGINGANSKVKSQVNDL
Oxy.tri_799                    QPSNASSTTQTSLSSFLSKNKG............EAANKADQVNGINGANSKAKSQVNDL
Cam.sin_964                    NPNGL.........................................TTEEFKKLQISDKI
Vor.cam_9330                   ...............................................MECFKKMRLSNEL
Par.bur_P04390037              GATI............................................QEFQQMVIPESM
Par.bur_P09660035              GATI............................................QEFQQMVIPESM
Par.tet_001442122.1            HLKI............................................EQFKQFNLTDSQ
Tet.mal_00222350_PEP_          GQSG.........................................KSIEDLKKLAISDKL
Tet.the_AMT6                   GQSG.........................................KSIEDLKKLAISDKL
Tet.ell_00078280_PEP_          DQSG.........................................KQIEDLKKLAINDKL
Tet.vor_00219940_PEP_          .....................................................LSINEKL
Tet.bor_00193820_PEP_          GEEG.........................................KTIQDLKKLSISDKL
Tet.can_00194390_PEP_          GEEG.........................................KTIQDLKKLSISDKL
Tet.pyr_00246380_PEP_          GQEG.........................................KTIQDLKNLSINEKL
Tet.epi_00174830_PEP_          SSPG.........................................KTIEDLKKLSISDKL
Tet.par_00207750_PEP_          SQNGDS.....................................KLKSIDQFKNINLNEKL
Ich.mul_156080                 ............................................................
Par.bur_P01400001              ................................................DFKYLDLN.TTL
Par.bur_P02340001              ................................................EFKYLDLN.TTL
Par.tet_GSPATP00024232001      ................................................DLKNLDIS.KKL
Par.tet_GSPATP00005554001      ................................................DLKNLDLS.KKL
Vor.cam_9610                   ...........................................GQNMKDFQHLRIESTKL
Vor.cam_12567                  ...........................................GQNMKDFQHLRIESTKL
Vor.cam_9611                   ...........................................GQNMKDFQHLRIESTKL
Tet.the_AMT7                   RKEKE......................................GLNINDLSNLKIPLS.L
Tet.mal_00069950_PEP_          RKEKE......................................GLNINDLSNIKIPLS.L
Tet.ell_00008460_PEP_          RKEKE......................................GLNINDLTNLKIPLS.L
Tet.pyr_00085530_PEP_          RKEKE......................................GQNINDLSNIKIPLS.L
Tet.vor_00057090_PEP_          RKEKD......................................GQNITDLSNIKMPLS.F
Tet.bor_00075080_PEP_          RKEKD......................................GQNINDLNNLKLPLS.L
Tet.can_00083510_PEP_          RKEKD......................................GQNINDLNNLKLPLS.L
Tet.epi_00101720_PEP_          RKEKD......................................GQNIADFASLKLPNS.L
TEPIDO00091130_PEP_            RKEKE......................................GQNISDFPNLKLPNS.L
Tet.par_00104560_PEP_          KKSKD......................................GQDISEFKQIRLPPS.V
Ich.mul_AMT7                   RQGKE......................................GIDIKEFSQFKIDLN.L
Cam.sin_20221                  ..KSE......................................GINLEKFRNLQIDNNKL
Cam.sin_20222                  ..KSE......................................GINLEKFRNLQIDNNKL
Cam.sin_7583                   ..KKE......................................GINIKDFRKIQIENRKL
Oxy.tri_21422                  ...ARKRTRRQDVDEDEEAIEV.....................NQVESSDAEVEIPANDI
Oxy.tri_42188                  MLLQEHKKRGKKYLQDEIVKEE.....................DFDEDDDSDEDIHMDDL
Oxy.tri_11745                  KETIFQIIMPWRQE.............................KEDEEKKKVKKQKNENE
Oxy.tri_31967                  LQYLVDQIGHFEVVQIDPPWRIKG.AQRNNTSFMFSNNKFNLDYSTMSNAEIMNIPVEVL
```

```
                               230       240                                 250       260
Oxy.tri_794                    VAQGLAQIVQSK......................EPIVKTNFDKYAKNVEAILINPCWQT
Oxy.tri_24932                  VAQGLAQIVQSK......................EPIVKTNFDKYAKNVEAILINPCWQT
Oxy.tri_799                    VAQGLAQIVQSK......................EPIVKTNFDKYAKNVEAILINPCWQT
Cam.sin_964                    LNNGMVFVWADK......................ENMNEIMDHLETMGIIYVEIFSFILI
Vor.cam_9330                   LSNGMVFIWGGK......................TNIGELMDYFDQMGYVYAENFTFVLL
Par.bur_P04390037              MSNGILFIWCDK......................DNIMEIIDHLDSQGFNYVENFTMVIL
Par.bur_P09660035              MSNGILFIWCDK......................DNIMEIIDHLDSQGFNYVENFTMVIL
Par.tet_001442122.1            MHNGMLFIWCEK......................DNIMEIIDQLEKLGFNYVENFTTVLL
Tet.mal_00222350_PEP_          INNGIIFIWSEK......................EILGQIIDVLEAKGFNYIENFMINQL
Tet.the_AMT6                   INNGIIFIWSEK......................EILSQIVDVLEAKGFNYIENFMINQL
Tet.ell_00078280_PEP_          INNGIIFIWSEK......................EILGQIIDVLEAKGFSYIENFMINQL
Tet.vor_00219940_PEP_          MNNGIIFIWSEK......................EILGQIIDVLEVKGFQYIENFMINQL
Tet.bor_00193820_PEP_          MNNGIIFIWSEK......................EILGQIIDVLEQKGFQYIENFMINQL
Tet.can_00194390_PEP_          MNNGIIFIWSEK......................EILGQIIDVLEQKGFQYIENFMINQL
Tet.pyr_00246380_PEP_          MNNGIIFIWSEK......................EILSQIFDVLESKGFQYIENFMINQL
Tet.epi_00174830_PEP_          INNGIIFIWSEK......................IILGKIVDVLETKGFQYIENFMVTQL
Tet.par_00207750_PEP_          MNNGIIFVWSEK......................IILSDIITIMEQKGFQYIENFMISQL
Ich.mul_156080                 .......................................................MVAQL
Par.bur_P01400001              MSHGILFVWSDK......................TKLNEMIDCFEEKGFVYIENLVVIQL
Par.bur_P02340001              MNHGILFVWSDK......................TKLNEMIDCFEEKGFVYIENLVVIQL
Par.tet_GSPATP00024232001      ISNGILFIWSDK......................SLINEILETMENKGFTYIENLVVVQL
Par.tet_GSPATP00005554001      ISNGILFIWSDK......................GLINEILEIMESKGFTYIENLVVVQL
Vor.cam_9610                   MAKGLVFVWAPK......................QNLSEMLSIMESKDFTYVENLEIINF
Vor.cam_12567                  MAKGLVFVWAPK......................QNLSEMLSIMESKDFTYVENLEIINF
Vor.cam_9611                   MAKGLVFVWAPK......................QNLSEMLSIMESKDFTYVENLEIINF
Tet.the_AMT7                   MKNGILFIWSEK......................EILGQIVEIMEQKGFTYIENFSIMFL
Tet.mal_00069950_PEP_          MKNGILFIWSEK......................EILGQIVEIMEQKGFTYIENFSIMFL
Tet.ell_00008460_PEP_          MKNGILFIWSEK......................EILGQIVEIMEQKGFTYIENFSIMFL
Tet.pyr_00085530_PEP_          MKNGIVFVWSEK......................EILGQIVDIMEEKGFTYIENFSIMFL
Tet.vor_00057090_PEP_          MKNGIVFIWSEK......................EILSQIVDIMEEKGFTYIENFSIMFL
Tet.bor_00075080_PEP_          MKNGIVFIWSEK......................EILGQIVDIMEEKGFTYIENFSIMFL
Tet.can_00083510_PEP_          MKNGIVFVWSEK......................EILGQIVDIMEEKGFTYIENFSIMFL
Tet.epi_00101720_PEP_          MKNGIIFIWSEK......................EILGQIVEMMEEKGYTYIENFSIMFL
TEPIDO00091130_PEP_            MKNGIIFVWSEK......................EILGQIVEMMEEKGYTYIENFSIMFL
Tet.par_00104560_PEP_          IRNGIIFIWSEK......................EILGEIIDIMEEKGFTYIENFSAIFL
Ich.mul_AMT7                   MKNGIVFIWSEK......................QILGQIIDCMIEKGFQYIENFSIIFL
Cam.sin_20221                  MSKGIVFIWSEK......................EIMCRLMDIMEEKGFVYIENFHIIYL
Cam.sin_20222                  MSKGIVFIWSEK......................EIMCRLMDIMEEKGFVYIENFHIIYL
Cam.sin_7583                   MSKGIVFLWSEK......................ELTGELMDVMEEKGFVYVENFQIIHL
Oxy.tri_21422                  DTESYTEGTNKRKQKLK....AKKQVLDKKKNKTEGDIDKEDAVEEEETVFIDNLPNDEF
Oxy.tri_42188                  ENESLSFDRNNR....K....SHKPVCKRTREENILDADLGDEKDDEDTIFIDNLPSDEF
Oxy.tri_11745                  LKQSSLSNWMKK...............PPAPKKNPKDDDQNKVISKEELKYQEHKFGNEE
Oxy.tri_31967                  SRKGFCFLWILNSQMHIGYECMNKWGYEVVDQLTWIKTKNKKIHISHGFYFLHSSEVCLV
```

```
                              270       280       290       300       310       320
Oxy.tri_794                   QKDKNGKVKGVTMEEFQKLNFSKDLMIDGLVFVWVEKDIIYPVIKFFETQSLFYVENVCW
Oxy.tri_24932                 QKDKNGKVKGVTMEEFQKLNFSKDLMIDGLVFVWVEKDIIYPVIKFFETQSLFYVENVCW
Oxy.tri_799                   QKDKNGKVKGVTMEEFQKLNFSKDLMIDGLVFVWVEKDIIYPVIKFFETQSLFYVENVCW
Cam.sin_964                   STEKIKTFTGSEKKQVKKERSILDFFKKKEKIEENKTEVIKPEKNYFE............
Vor.cam_9330                  SRPKIPQKS...QKQLAGNKSLLNFFSKPNPTQPEKKIETKSLVPKLE............
Par.bur_P04390037             SREKILTMN.......DKTKKITDFFTKKEKKEQKLNDKQE...................
Par.bur_P09660035             SREKILTMN.......DKTKKITDFFTKKEKKEQKLNDKQE...................
Par.tet_001442122.1           SLEKILTIMP....SEPQVRKITEFFSKKNVKEQKEIKKIDGL.................
Tet.mal_00222350_PEP_         SADKALEMQRKNQNQQSKEKKITDFFKRLTPQKNIWSDITPEQCIE..............
Tet.the_AMT6                  SADKALEMQRKNQNQQSKEKKITDFFKRLTPQKNIWSDITPEQCIE..............
Tet.ell_00078280_PEP_         SADKALEMQRKNQNQQSKEKKITDFFKRLTPQKNIWSDITPEQCIE..............
Tet.vor_00219940_PEP_         SVDKALEIQKISQNQQSKEKKITDFFKRITPVKNIWSDITSEQCIE..............
Tet.bor_00193820_PEP_         SADKALEMQRKNLNQQSKEKKITDFFKRLTPQKSIWSDITAEECIE..............
Tet.can_00194390_PEP_         SADKALEMQRKNLNQQSKEKKITDFFKRLTPQKSIWSDITAEECID..............
Tet.pyr_00246380_PEP_         SVDKALEMQRKNMNQQSKEKKITDFFKRITPQKNIWSEITPDSCIE..............
Tet.epi_00174830_PEP_         SADKALEMQRKNQNQQNKEKKITDFFKRITPQKDVWSSITPEEMIE..............
Tet.par_00207750_PEP_         SADKALEIQKSNKNNASKEKKITDFFKRLTPQKDIWSDMTPEKIIE..............
Ich.mul_156080                NSQKSLEIQRKKINQ.NKEKKITDFFKRLQPNKSIWDSLTPQQIVI..............
Par.bur_P01400001             NQKQAHSELIKYKDIP...NENQAVMDN................................
Par.bur_P02340001             NQTQAHSELKKYKDIP...NENQAVMDN................................
Par.tet_GSPATP00024232001     SLEQALEELNKHMKIE...QTEDAVLDN................................
Par.tet_GSPATP00005554001     SLEKALEELNKHMKIE...QTEEAVLDN................................
Vor.cam_9610                  DIEKAKSLVRKGSPQESGEKNEELDWAR................................
Vor.cam_12567                 DIEKAKSLVRKGSTQESGEKNEELDWAR................................
Vor.cam_9611                  DIEKAKSLVRKGSPQESGEKNEELDWAR................................
Tet.the_AMT7                  GLNKCLQSINHKDEDSQNSTASTNNTNNEAITSDLTLKDTSKFSDQIQDNHSEDSDQARK
Tet.mal_00069950_PEP_         GLNKCLSSINHKDEDSQNSTASTNNTNNEAITSDMTLKDSSKFSDQIQDNHSEDSDQARK
Tet.ell_00008460_PEP_         GLNKCLQSINNKDEDSQNSTASTNNTNNEAITSDLTLKESSKFSDQLHDNHSEDSDQARK
Tet.pyr_00085530_PEP_         SLEKCIQAIKHKDEDSQNSTASTNNTNNEVITSDLTLKDSSKFSDPINDNNSEDSEQAKK
Tet.vor_00057090_PEP_         SLEKCVQAINHKDEDSQNSTASTNNTNNEVITSDLTLKDSSKFSDPINDINSEDSEQAKK
Tet.bor_00075080_PEP_         SLEKSVSAINNNNEDSQNSTASTNNTINEAITSDLTLKDSSKFSEQINDNNSDDSDQARK
Tet.can_00083510_PEP_         SLEKSVSAINNNNEDSQNSTASTNNTINEAITSDLTLKDSSKYSEQINDNNSEDSDQARK
Tet.epi_00101720_PEP_         SLDKCVSAIKNKDEDSQNSTASTNNTINDPITSDLTLKDQSKFSDPLIDN.SDESDQTSK
TEPIDO00091130_PEP_           SLDKCVSAIKNKDEDSQNSTASTINTTNDAITSDLTLKDSSKFSDPLIDN.SDESDQARK
Tet.par_00104560_PEP_         SLDKSLNAIKRITDEQKAGHSHSIQHSNESIPSDITLKDSTKYSETTNETSEDSEETRKV
Ich.mul_AMT7                  DQNKCLSEINKDQKQETQQEKDKNQKNLRLIQKKAFQKHSSRWKNYIKNSNLNGRKKISN
Cam.sin_20221                 SAEKGLKELDKNYIKQNSN.KEIKKEVSSSKTISDSIDED....................
Cam.sin_20222                 SAEKGLKELDKNYIKQNSN.KEIKKEVSSSKTISDSIDED....................
Cam.sin_7583                  SLEKGLNELEKSCTKQIVSEKSSNKKELNSKTVSDASVED....................
Oxy.tri_21422                 EIRRMLKEVKKHIKSLEKQFFEEEDSEKEEELKQINNNSKHEEALQAFKETSHLKQFWCI
Oxy.tri_42188                 SIRRQLQDVKSYIKQFEMLFFEEEDSDKEEQLKQITNVQKHEEALQNFKDRSHLKNFWCI
Oxy.tri_11745                 DDEDMSRVSKVIKVKEVQSIRLTDYTDKDAHVQAILINPKWKISQDIYGDGDTEEGSMQE
Oxy.tri_31967                 GYKCPPNEYIEFKSKVSNNLIVAEIRKKSQKPEQIYTLIDILIPGAKKIELFARNNNLRR
```

```
                              330
Oxy.tri_794                   VMLDQT......................................................
Oxy.tri_24932                 VMLDQT......................................................
Oxy.tri_799                   VMLDQT......................................................
Cam.sin_964                   ............................................................
Vor.cam_9330                  ............................................................
Par.bur_P04390037             ............................................................
Par.bur_P09660035             ............................................................
Par.tet_001442122.1           ............................................................
Tet.mal_00222350_PEP_         ............................................................
Tet.the_AMT6                  ............................................................
Tet.ell_00078280_PEP_         ............................................................
Tet.vor_00219940_PEP_         ............................................................
Tet.bor_00193820_PEP_         ............................................................
Tet.can_00194390_PEP_         ............................................................
Tet.pyr_00246380_PEP_         ............................................................
Tet.epi_00174830_PEP_         ............................................................
Tet.par_00207750_PEP_         ............................................................
Ich.mul_156080                ............................................................
Par.bur_P01400001             ............................................................
Par.bur_P02340001             ............................................................
Par.tet_GSPATP00024232001     ............................................................
Par.tet_GSPATP00005554001     ............................................................
Vor.cam_9610                  ............................................................
Vor.cam_12567                 ............................................................
Vor.cam_9611                  ............................................................
Tet.the_AMT7                  QQTPDD.ITQKKNKLLKKSSVPSIQKLFEEDPVQTPSVNKP...................
Tet.mal_00069950_PEP_         QSTPEN.NTQKKSKSLKKSSVPSIQKLFEEDPVQTSSVNKP...................
Tet.ell_00008460_PEP_         QSNLEE.NNLKKSKLNKKNSVPSIQKLFEDDPVQTASVKQP...................
Tet.pyr_00085530_PEP_         PTSSSSDLSQKKSKAGKKSAPINIQKLFEEDPVIPLPAAKQQP.................
Tet.vor_00057090_PEP_         PTSSSSELTQKKGKAAKKSTAISIQKLFEDDLVVVPPVSKQ.P.................
Tet.bor_00075080_PEP_         PSSATN.ITQKKNKTTKKNAPLSIQKLFEEDPVATPIINKP...................
Tet.can_00083510_PEP_         PSSSTN.ITQKKNKTTKKNAPLSIQKLFEEDPVATPIINKP...................
Tet.epi_00101720_PEP_         PTKKTVSQKNKKTTDNKKGTPINIQKLFEEDPLLTSQTKAV...................
TEPIDO00091130_PEP_           PSKAITK..NKKIRADKKGNPINIQKLFEEDPPLPAQTKPV...................
Tet.par_00104560_PEP_         E.........EKGKKKKKKSVVSMNKLFEDLPTHQSNAPSTSAS................
Ich.mul_AMT7                  K...........................................................
Cam.sin_20221                 ............................................................
Cam.sin_20222                 ............................................................
Cam.sin_7583                  ............................................................
Oxy.tri_21422                 PLSVNVTTLDFDLLAKSQMKQG.GRLFDVITIDPPWQLSSAN..................
Oxy.tri_42188                 PLSSDVREIDWDVLIARQQEHTNGQLFDVITCDPPWQLSSAN..................
Oxy.tri_11745                 GGDSSSEEDVKQGKGMQMKSAKKQRANAFQGITMKEFSKLVFPQSLVYDGILFIWVEKEY
Oxy.tri_31967                 GWLSLGNQLGENYVNWKNVVICDDCNKDITTGNRYYKSKLKANYDICEKCFTVKQTNYAN
```

| | |
|---|---|
| Oxy.tri_794 | ............................................................ |
| Oxy.tri_24932 | ............................................................ |
| Oxy.tri_799 | ............................................................ |
| Cam.sin_964 | ............................................................ |
| Vor.cam_9330 | ............................................................ |
| Par.bur_P04390037 | ............................................................ |
| Par.bur_P09660035 | ............................................................ |
| Par.tet_001442122.1 | ............................................................ |
| Tet.mal_00222350_PEP_ | ............................................................ |
| Tet.the_AMT6 | ............................................................ |
| Tet.ell_00078280_PEP_ | ............................................................ |
| Tet.vor_00219940_PEP_ | ............................................................ |
| Tet.bor_00193820_PEP_ | ............................................................ |
| Tet.can_00194390_PEP_ | ............................................................ |
| Tet.pyr_00246380_PEP_ | ............................................................ |
| Tet.epi_00174830_PEP_ | ............................................................ |
| Tet.par_00207750_PEP_ | ............................................................ |
| Ich.mul_156080 | ............................................................ |
| Par.bur_P01400001 | ............................................................ |
| Par.bur_P02340001 | ............................................................ |
| Par.tet_GSPATP00024232001 | ............................................................ |
| Par.tet_GSPATP00005554001 | ............................................................ |
| Vor.cam_9610 | ............................................................ |
| Vor.cam_12567 | ............................................................ |
| Vor.cam_9611 | ............................................................ |
| Tet.the_AMT7 | ............................................................ |
| Tet.mal_00069950_PEP_ | ............................................................ |
| Tet.ell_00008460_PEP_ | ............................................................ |
| Tet.pyr_00085530_PEP_ | ............................................................ |
| Tet.vor_00057090_PEP_ | ............................................................ |
| Tet.bor_00075080_PEP_ | ............................................................ |
| Tet.can_00083510_PEP_ | ............................................................ |
| Tet.epi_00101720_PEP_ | ............................................................ |
| TEPIDO00091130_PEP_ | ............................................................ |
| Tet.par_00104560_PEP_ | ............................................................ |
| Ich.mul_AMT7 | ............................................................ |
| Cam.sin_20221 | ............................................................ |
| Cam.sin_20222 | ............................................................ |
| Cam.sin_7583 | ............................................................ |
| Oxy.tri_21422 | ............................................................ |
| Oxy.tri_42188 | ............................................................ |
| Oxy.tri_11745 | IMDVCKFLED.................................................. |
| Oxy.tri_31967 | KQGFIDESQHGYKASKNQYLERFYLIENATDEDILHEYHQCNICNAEPIWGLRFKCSSCE |

| | |
|---|---|
| Oxy.tri_794 | ............................................................ |
| Oxy.tri_24932 | ............................................................ |
| Oxy.tri_799 | ............................................................ |
| Cam.sin_964 | ............................................................ |
| Vor.cam_9330 | ............................................................ |
| Par.bur_P04390037 | ............................................................ |
| Par.bur_P09660035 | ............................................................ |
| Par.tet_001442122.1 | ............................................................ |
| Tet.mal_00222350_PEP_ | ............................................................ |
| Tet.the_AMT6 | ............................................................ |
| Tet.ell_00078280_PEP_ | ............................................................ |
| Tet.vor_00219940_PEP_ | ............................................................ |
| Tet.bor_00193820_PEP_ | ............................................................ |
| Tet.can_00194390_PEP_ | ............................................................ |
| Tet.pyr_00246380_PEP_ | ............................................................ |
| Tet.epi_00174830_PEP_ | ............................................................ |
| Tet.par_00207750_PEP_ | ............................................................ |
| Ich.mul_156080 | ............................................................ |
| Par.bur_P01400001 | ............................................................ |
| Par.bur_P02340001 | ............................................................ |
| Par.tet_GSPATP00024232001 | ............................................................ |
| Par.tet_GSPATP00005554001 | ............................................................ |
| Vor.cam_9610 | ............................................................ |
| Vor.cam_12567 | ............................................................ |
| Vor.cam_9611 | ............................................................ |
| Tet.the_AMT7 | ............................................................ |
| Tet.mal_00069950_PEP_ | ............................................................ |
| Tet.ell_00008460_PEP_ | ............................................................ |
| Tet.pyr_00085530_PEP_ | ............................................................ |
| Tet.vor_00057090_PEP_ | ............................................................ |
| Tet.bor_00075080_PEP_ | ............................................................ |
| Tet.can_00083510_PEP_ | ............................................................ |
| Tet.epi_00101720_PEP_ | ............................................................ |
| TEPIDO00091130_PEP_ | ............................................................ |
| Tet.par_00104560_PEP_ | ............................................................ |
| Ich.mul_AMT7 | ............................................................ |
| Cam.sin_20221 | ............................................................ |
| Cam.sin_20222 | ............................................................ |
| Cam.sin_7583 | ............................................................ |
| Oxy.tri_21422 | ............................................................ |
| Oxy.tri_42188 | ............................................................ |
| Oxy.tri_11745 | ............................................................ |
| Oxy.tri_31967 | DCDLCEQCFDSRLHKLNLLQEQNQSQNDDQSDSMQIDTYRNGHNKDKEDIIQVKPDDIIC |

```
                                              340       350       360
                                                .         .         .
Oxy.tri_794                 .....................KKQEVEATQSIDVSPAYVRDDYQYVRKSHKTLLMFRR..
Oxy.tri_24932               .....................KKQEVEATQSIDVSPAYVRDDYQYVRKSHKTLLMFRR..
Oxy.tri_799                 .....................KKQEVEATQSIEVSPAYVRDDYQYVRKSHKTLLMFRR..
Cam.sin_964                 .......................MPSYEFPNIDPVELFSNENYPYFKKAKKNLLMLRK..
Vor.cam_9330                .......................NDYEIGKVSDPAEVFLNGASEYFCESQRVLYMFRK..
Par.bur_P04390037           .......................LQLSDLPDLDPSTVFWNQDQQYFRKAKRTLLMFRK..
Par.bur_P09660035           .......................LQLSDLPDLDPSTVFWNQDQQYFRKAKRTLLMFRK..
Par.tet_001442122.1         .......................RNQEQLPDLDPTQVFWNEDYTYFRKTKRTLLMFRK..
Tet.mal_00222350_PEP_       ........................QEKFPPNNYVQDIFVNSEYCYFRKSKKILLMLRK..
Tet.the_AMT6                ........................QEKFPPNNYVQDIFVNSEYSFFRKSKKILLMLRK..
Tet.ell_00078280_PEP_       ........................QEKFPPNNYVQDIFVNSEYSYFRKSKKILLMLRK..
Tet.vor_00219940_PEP_       ........................QEKFPPNNYVQDIFLNSDYSYFRKSKKILLMFRK..
Tet.bor_00193820_PEP_       ........................QDKFPPNNYVQDIFVNSDYSYFRKSKKILLMFRK..
Tet.can_00194390_PEP_       ........................QDKFPPNNYVQDIFVNSDYSYFRKSKKILLMFRK..
Tet.pyr_00246380_PEP_       ........................QDKFPPNNYVQDIFMNSEYDYFTKTKKILLMFRK..
Tet.epi_00174830_PEP_       ........................QDKFPANNYVQDIFVNSDYEYFRKSKKILLMFRK..
Tet.par_00207750_PEP_       ........................QEKFPQINNVKDIFVNGEYPYFRISKKVLLMFRK..
Ich.mul_156080              ........................QEKYPEINNVVEIFLNEDDNYFKRAKKVILMFRK..
Par.bur_P01400001           ........................INMLQEKVSLSQMIAKRPYQIFNRSKQVLVMFR...
Par.bur_P02340001           ........................IHLLQEKVSLNQMIAKRPYQIFNKSKQVLVMFR...
Par.tet_GSPATP00024232001   ........................LDFLQQKVQVKDLILHRPSKVLNQSKQVLIMFR...
Par.tet_GSPATP00005554001   ........................LNFLQQKVQVKDLIVNCPSKVLNQSKQVLIMFR...
Vor.cam_9610                ........................LIKALENVDPCDIIADEHSRFFNSSKRTLLMFK...
Vor.cam_12567               ........................LIKALENVDPCDIIADEHSRFFNSSKRTLLMFK...
Vor.cam_9611                ........................LIKALENVDPCDIIADEHSRFFNSSKRTLLMFK...
Tet.the_AMT7                .......IEKSIEQVTQEKKFVMNNLDILKSTDINNLFLRNNYPYFKKTRHTLLMFR...
Tet.mal_00069950_PEP_       .......TEKNTEQITQEKKYVMNNLDILKSTDINNLFLRNNYPYFKKTRHTLLMFR...
Tet.ell_00008460_PEP_       .......IEKTVEQVTQDKKYVMNNLDILKSTDINNLFLRNNYPYFKKTRHTLLMFR...
Tet.pyr_00085530_PEP_       .......AEKTIDQITEEKNFVLDNFETLKSSDINGLFLRNNYPYFKKTRHTLLMFR...
Tet.vor_00057090_PEP_       .......TEKTADQIAEDKQFVMDNFETLKSSDINGLFLRNNYPYFKKTRHTLLMFR...
Tet.bor_00075080_PEP_       .......IEKTEREITQQKNFVMDNLDILKSSNINDLFLKNNYPYFKKTRHTLLMFR...
Tet.can_00083510_PEP_       .......IEKTEREITQQKNFVMDNLDILKSSNINDLFLKNNYPYFKKTRHTLLMFR...
Tet.epi_00101720_PEP_       ........EKTASQIQEEKNFVMNNLDILKSSDINNLFLRNNYPYFKKTRHTLLMFR...
TEPIDO00091130_PEP_         ........EKTQSQILEEKDFVMNNLDILKSSEINNLFLRNNYPYFKKTRHTLLMFR...
Tet.par_00104560_PEP_       .....SNQYKTEQQIEDDKQYVMKNLDKLKLEPINNLFLKNSYPYFKKTKYTLLMFR...
Ich.mul_AMT7                ........QFTQNKAIQYQQFAFKKLYLLQKTKHTLLMFRR...VIFIIILIFIFIK...
Cam.sin_20221               ...............NEIKEEFIQNMNRLGNIEANNLFLQEDGSYFKKSKKMLLMFR...
Cam.sin_20222               ...............NEIKEEFIQNMNRLGNIEANNLFLQEDGSYFKKSKKMLLMFRKVY
Cam.sin_7583                ...............LERKEEFLGNLHKLGNLEANKILLEKDGQFLKKSKKNLLMFR...
Oxy.tri_21422               ..............PTRGVAIAYDTLNDKEILNMPFEKVQTDGFLFIWVINAKYRFALEM
Oxy.tri_42188               ..............PTRGVAIAYETLNDGEILKIPWGRLQKDGFLFIWVINAKYRFALDM
Oxy.tri_11745               ....QNFFYIENMCWIMLDENMREEVEKTRLQDATPAFVRENYTYFKKSKKTLLMFRR..
Oxy.tri_31967               REHDFMCIELPLIADGLAAHNQYKCVSCYMRPIIGACFVCADCNHFSLCQNCYFTRPFQN

                                 370       380       390       400
                                   .         .         .         .
Oxy.tri_794                 ....MHKKNGNPLELRHQRTCDVCFDFVETDVHNYKPN................EYIYKL
Oxy.tri_24932               ....MHKKNGNPLELRHQRTCDVCFDFVETDVHNYKPN................EYIYKL
Oxy.tri_799                 ....MHKKNGNPLELRHQRTCDVCFDFVETDVHNYKPN................EYIYKL
Cam.sin_964                 ....INKKET..LELRHQRTSDVYFTLYDE.KNHSIDEK.............GFECIYKM
Vor.cam_9330                ....INKKES..LELRHQRTSDVFFTIGDD.K..GLDWK.............GKEIVYRM
Par.bur_P04390037           ....ISKQN...LELRHQRTGDIFFDVVD..DPQGISDI.............GLQYVYQM
Par.bur_P09660035           ....ISKQN...LELRHQRTGDIFFDVVD..DPQGISDI.............GLEYVYKM
Par.tet_001442122.1         ....QSKQQ...LELRHQRTGDIFFDVIEK.DAQNISAI.............GMEYIYKM
Tet.mal_00222350_PEP_       ....FNKDAQ..LELRHQRTSDIFFDIFEQDKPNDVSKK.............GMEFVYKM
Tet.the_AMT6                ....FNKDAQ..LELRHQRTSDIFFDIFEQNKPNDVSKK.............GMEFVYKM
Tet.ell_00078280_PEP_       ....FNKDAQ..LELRHQRTSDIFFDIFEQNKPNDVSKK.............GMEFVYKM
Tet.vor_00219940_PEP_       ....FNKDAQ..LELRHQRTSDIFFDVFAQKNPNDISKR.............GMEFAYRM
Tet.bor_00193820_PEP_       ....FNKDAQ..LELRHQRTSDIFFDIFCQKKPNDISKK.............GMEFVYKM
Tet.can_00194390_PEP_       ....FNKDAQ..LELRHQRTSDIFFDIFCQKKPNNISKK.............GMEFVYKM
Tet.pyr_00246380_PEP_       ....FNKDAQ..LELRHQRTSDIFFDVFEQNKPNNVSKK.............GMEFVYRM
Tet.epi_00174830_PEP_       ....FNKEAQ..LELRHQRTSDIFFDVFQQSKINEVSKK.............GMEFTYRM
Tet.par_00207750_PEP_       ....FNKDQQ..LELRHQRTSDIFFDLFSEKNPNNVSQK.............GMEFVYKM
Ich.mul_156080              ....FNKDAQ..LELRHQRTSDVFFDIFNEQKPDDLSQK.............GMQFIYKM
Par.bur_P01400001           ....RFDEMKTPLELRHQRTSDVVFDVCDE...PFSEK..............LKEYLYKL
Par.bur_P02340001           ....RFDEMKTPLELRHQRTSDVVFDVSDE...PFSEK..............LKEYLYKL
Par.tet_GSPATP00024232001   ....KFDEQKSQLELRHQRTPDVLFDIVNNG..KSCLK..............TKEYIYQT
Par.tet_GSPATP00005554001   ....KFDEQKTQLELRHQRTPDVLFDFVSNG..KKSEK..............SKEYIYQT
Vor.cam_9610                ....KNSEIK..FELRHQRTCDVVFDMPLRLGEKCNSD..............IKAYLYNM
Vor.cam_12567               ....KNSEIK..FELRHQRTCDIVFDMPLRLGEKCNSD..............LKAYLYNM
Vor.cam_9611                ....KNSEIK..FELRHQRTCDVVFDMPLRLGEKCNSD..............IKAYLYNM
Tet.the_AMT7                ....RIGDKNQKLELRHQRTSDVVFEVTDEQDPS.............KVDTMMKEYVYQM
Tet.mal_00069950_PEP_       ....RIGDKNQKLELRHQRTSDVVFEVTDEQDPS.............KVDTMMKEYVYQM
Tet.ell_00008460_PEP_       ....RIGDKNQKLELRHQRTSDVVFEVTDEQDPS.............KVDFMMKEYVYQM
Tet.pyr_00085530_PEP_       ....RIGDKNVKLELRHQRTSDVVFEITDEHDPA.............KIDTMMKEYVYQM
Tet.vor_00057090_PEP_       ....RIGDKNVKLELRHQRTSDVVFEVTDEHDPA.............KIDTMMKEYVYQM
Tet.bor_00075080_PEP_       ....RIGDKNVKLELRHQRTSDVVFEITDENDPA.............KLDTMMKEYVYQM
Tet.can_00083510_PEP_       ....RIGDKNVKLELRHQRTSDVVFEVTDENDPA.............KLDTMMKEYVYQM
Tet.epi_00101720_PEP_       ....RVGDKNVKLELRHQRTSDVVFEVAEDYDPS.............KIDTMMKEYVYQM
TEPIDO00091130_PEP_         ....RVGDKNVKLELRHQRTSDVVFEVTEDQDPS.............KIDTMMKEYVYQM
Tet.par_00104560_PEP_       ....KFADKTQQLELRHQRTSDVVFEVSDESDPS.............KIDNMLKEYVYQM
Ich.mul_AMT7                ....QFGEKNQQLELRHQRTPDVVFELTEQNDPSSLFIILYHFYQVIDIDKQMKEYIYQM
Cam.sin_20221               ....KHDNKKDNLELRHQRTCDVLFDLVTQSNTT..............SDMKTKNYVYKM
Cam.sin_20222               LIETEHDNKKDNLELRHQRTCDVLFDLVTQSNTT..............SDMKTKNYVYKM
Cam.sin_7583                ....KHDNKKDNLELRHQRTCDVLFDLVVPENVG..............CDIKTKNYIYKM
Oxy.tri_21422               MEKFGYKLVDEIAWVKQTVNGKIAKGHGYYLQHAKETCLVG....VKGN...VKGKARYN
Oxy.tri_42188               MGAHGYRVVDEIQWVKQTCNGKIAKGHGYYLQHAKEVCLVG....CKGDPAILAKKCRSN
Oxy.tri_11745               ....LSTDVNNKLELRHQRTGDVVFDWKDTINPHSKPQ...............YYTYKL
Oxy.tri_31967               LKVRGHTPNHHIELIVEPRQTVKKYVKCNGCGEMPIQDVRYKCNNCFDFDFCEKCYQLYC
```

```
                           410       420       430
Oxy.tri_794                IETLLPQASIDEE.KKH...VRLVELWAH...........................DSTP
Oxy.tri_24932              IETLLPQASIDEE.KKH...VRLVELWAH...........................DSTP
Oxy.tri_799                IETLLPQASIDEE.KKH...VRLVELWAH...........................DSTP
Cam.sin_964                IETLLIKANYSK..GES...LKMMELWAKE..........................NKP.
Vor.cam_9330               IETLLISANFKE..KEH...LKLMELWSTK..........................DYT.
Par.bur_P04390037          IETLLPKAKYSN..GEQ...LKMMELFADP..........................KSTG
Par.bur_P09660035          IETLLPKAKYSN..GEQ...LKMMELFADP..........................KSTG
Par.tet_001442122.1        IETLLPKAQYKA..GEG...LKMMELFADP..........................NTFG
Tet.mal_00222350_PEP_      IETLLPKANYSEENKGA...LKMMELYADD..........................KSQP
Tet.the_AMT6               IETLLPKANYSEENKGA...FKMMELYADD..........................KSQP
Tet.ell_00078280_PEP_      IETLLPKANYSEENKGA...LKMMELYADE..........................KSQP
Tet.vor_00219940_PEP_      IETLLPKANYSEENKGA...FKMMELYATD..........................KSQP
Tet.bor_00193820_PEP_      IETLLPKANYCEENKGG...LKMMELFSTD..........................KTQP
Tet.can_00194390_PEP_      IETLLPKANYCEENKGG...LKMMELFSTD..........................KTQP
Tet.pyr_00246380_PEP_      IETLLPKANFSEENKGA...FKMMELYATD..........................KSQP
Tet.epi_00174830_PEP_      IETLLPKANYCEENKGS...FKMMELYANE..........................GTQP
Tet.par_00207750_PEP_      IETLLPKARYSPSSND....FKMMELFADN..........................KTQP
Ich.mul_156080             IETLLPKANYSEDNKQS...LKLMELWANP..........................EILP
Par.bur_P01400001          IETLLP.............ISKFLEIYAD...........................PDRP
Par.bur_P02340001          IETLLP.............ISKFLEIYAD...........................PDRP
Par.tet_GSPATP00024232001  IETLLP.............KSQLMEIFAQ...........................KDQP
Par.tet_GSPATP00005554001  IETLLP.............KSQLMEIFAQ...........................RDQP
Vor.cam_9610               IETLLPDCLTVNQIRKNEFEFKMMELWAD...........................KGDY
Vor.cam_12567              IETLLPDCLTVNQIRKNEFEFKMMELWAD...........................KGDY
Vor.cam_9611               IETLLPDCLTVNQIRKNEFEFKMMELWAD...........................KGDY
Tet.the_AMT7               IETLLPKAQFIPGVDK...HLKMMELQN...........................QLIFL
Tet.mal_00069950_PEP_      IETLLPKAQFIPGVDK...HLKMMELFA...........................NPDNY
Tet.ell_00008460_PEP_      IETLLPKAQFIPGVDK...HFKMMELFA...........................NSENY
Tet.pyr_00085530_PEP_      IETLLPKAIYTPGVDK...HLKMMELFA...........................NADNC
Tet.vor_00057090_PEP_      METLLPKAVYTPGVDK...HFKMMELFA...........................NPENC
Tet.bor_00075080_PEP_      METLLPKALYTPGVDK...NFKMMEIFA...........................NPENA
Tet.can_00083510_PEP_      METLLPKALYTPGVDK...NFKMMEIFA...........................NPENA
Tet.epi_00101720_PEP_      METLLPKAIYTPGVDK...HFKMMELFA...........................NPENT
TEPIDO00091130_PEP_        METLLPKAIYNPSVDK...HFKMMELFA...........................NPENT
Tet.par_00104560_PEP_      IETLLPKAQYIPGVDK...HLKMLELFG...........................NSSRP
Ich.mul_AMT7               IETLLPKALNNQKTDK...FFKMMELQN...........................QI...
Cam.sin_20221              IETLLPRSLENLESNG...HLSMMELELIF....................EYRWGDKNSP
Cam.sin_20222              IETLLPRSLENLESNG...HLSMMELELIF....................EYRWGDKNSP
Cam.sin_7583               IETLLPRSLENLESNA...SLSMMEL...........................WGDKNSH
Oxy.tri_21422              IESDVIFSQRRGQSQKPEEIYEIAEALVPNGY.................YLEIFGRRNNL
Oxy.tri_42188              IESDVIFSERRGQSQKPEEIYELVEALVPNGY.................YMEIFGRRNNL
Oxy.tri_11745              IETLLPKAMVDLKKQPH...LRMIELWAED...........................DTP
Oxy.tri_31967              VEKKELKTVYSTSHKGYHTFTRLMLASSNAVANNTKDQNQGELLEDGETQTKGKSNRNRK

                            440
Oxy.tri_794                RKGWIKFIE............................
Oxy.tri_24932              RKGWIKFIE............................
Oxy.tri_799                RKGWIKFIE............................
Cam.sin_964                RKGWISISEKNM.........................
Vor.cam_9330               RKGWITVSEESIPMQ......................
Par.bur_P04390037          RLGWVQVIKE...........................
Par.bur_P09660035          RLGWVQVIKE...........................
Par.tet_001442122.1        RSGWIQVIQE...........................
Tet.mal_00222350_PEP_      RKGWISVYEQE..........................
Tet.the_AMT6               RKGWISVYEQE..........................
Tet.ell_00078280_PEP_      RKGWISVFEQE..........................
Tet.vor_00219940_PEP_      RKGWISVFEQK..........................
Tet.bor_00193820_PEP_      RKGWISVFEQEQ.........................
Tet.can_00194390_PEP_      RKGWISVFEQEQ.........................
Tet.pyr_00246380_PEP_      RKGWISVFEQN..........................
Tet.epi_00174830_PEP_      RKGWISVYEESE.........................
Tet.par_00207750_PEP_      RQGWISVFEEN..........................
Ich.mul_156080             QKGWIRIIENEQN........................
Par.bur_P01400001          RKGWYSICESK..........................
Par.bur_P02340001          RKGWYSICESK..........................
Par.tet_GSPATP00024232001  RKGWISVCENK..........................
Par.tet_GSPATP00005554001  RKDWISVCESK..........................
Vor.cam_9610               RRGWISVVQE...........................
Vor.cam_12567              RRGWISVVQE...........................
Vor.cam_9611               RRGWISVVQE...........................
Tet.the_AMT7               LFKNLKNYQLKQICQH.....................
Tet.mal_00069950_PEP_      RNGWISVVEK...........................
Tet.ell_00008460_PEP_      RPGWISVVEK...........................
Tet.pyr_00085530_PEP_      RPGWVSVVEK...........................
Tet.vor_00057090_PEP_      RPGWVSVVEK...........................
Tet.bor_00075080_PEP_      RQGWISVNEK...........................
Tet.can_00083510_PEP_      RQGWISVNEK...........................
Tet.epi_00101720_PEP_      RPGWISVIEK...........................
TEPIDO00091130_PEP_        RPGWISVIEK...........................
Tet.par_00104560_PEP_      REGWISVCEK...........................
Ich.mul_AMT7               .....................................
Cam.sin_20221              RKGWIILNEK...........................
Cam.sin_20222              RKGWIILNEK...........................
Cam.sin_7583               RKGWIILTEK...........................
Oxy.tri_21422              HNGWVTIGNEL..........................
Oxy.tri_42188              HNGWVTVGNEL..........................
Oxy.tri_11745              RKGWIKFIGLNQ.........................
Oxy.tri_31967              VQSQIATTQEGVKIKRPPGRPAGKSKSKKQINQDSQL
```
