## Supplemental file S3 for "Structural basis of DNA N^6^-adenine methylation in eukaryotes"

```
             1        10        20        30        40        50        60
Otri-p1-1   MRVYLKFCNRKQIHYTHTMSSSISAAIMAGNQNKKIAESKSLWNYALSPGWTQQEVEILK
Otri-p1-2   MRVYLKFCNRKQIHYTHTMSSSISAAIMAGNQNKKIAESKSLWNYALSPGWTQQEVEILK
Otri-p1-5   ..................MSSSISAAIMAGNQNKKIAESKSLWNYALSPGWTQQEVEILK
Slem-p1     ...................................MADSKSLHNYTLSPGWTREEVDILK
Otri-p1-4   .............................MSVHHKMADSKSLHNYTLSPGWTREEVDILK
Hgra-p1     .........MSGVQQNRAMSKNAGNQQIVVGTVPKIADSKSLHNYTLSPGWTKEEVEILK
Otri-p1-3   ..................MSSSISAAIIAGNQNKKIAESKSLWNYALSPGWTQQEVEILK
Tthe-p1     ..........................MSLKKGKFQHNQSKSLWNYTLSPGWREEEVKILK
Pper-p1     .................................MEQGNSNSLWNYTLSPGWNEQEVQVLK
Ptet-p1     .............................MSKQIVHHQSQSLWNYTLSPGWTMKEVEILN

                     70        80        90        100       110       120
Otri-p1-1   IALMKFGVGRWSAINKSGVLPTKQIQQCYLQTQRLIGQQSLAEFMGLHLDIDRIAADNKQ
Otri-p1-2   IALMKFGVGRWSAINKSGVLPTKQIQQCYLQTQRLIGQQSLAEFMGLHLDIDRIAADNKQ
Otri-p1-5   IALMKFGVGRWSAINKSGVLPTKQIQQCYLQTQRLIGQQSLAEFMGLHLDIDRIAADNKQ
Slem-p1     IALMKFGIGKWKKIQKSGCLPSKTISQMNLQTQRVLGQQSLAEFMGLHG...........
Otri-p1-4   IALMKFGIGKWKKIQKSGCLPSKTISQMNLQTQRLLGQQSLAEFMGLHVYLDRVFRDNSL
Hgra-p1     ICLMKFGIGRWKKIIKCGCLPSKTISQMNLQTQRLLGQQSLAEFMGLHVHLDRVFNDNLK
Otri-p1-3   IALMKFGVGRWKTIEQSQCLPTKTMSQMYLQTQRLVGQQSLAEFMGLHLDLEQIFIKNAE
Tthe-p1     SALQLFGIGKWKKIMESGCLPGKSIGQIYMQTQRLLGQQSLGDFMGLQIDLEAVFNQNMK
Pper-p1     QALQYFGIGKWKQIQEHQCLPDKTISQIYMQTQRLLGQQSLGDLFGLRLHLDKVWENNKK
Ptet-p1     LALMKFGIGKWRKIINSECLPGKSIGQIYMQTQRLLGQQSLGEFMGLQVDLKKVYLHNMQ

                       130       140       150       160
Otri-p1-1   K...RGIRKQGFLVNQGCKLTPEEKDELRKINQEKYGLTAEHVEAIKLPAP.........
Otri-p1-2   K...RGIRKQGFLVNQGCKLTPEEKDELRKINQEKYGLTAEHVEAIKLPAP.........
Otri-p1-5   K...RGIRKQGFLVNQGCKLTPEEKDELRKINQEKYGLSAEHVEAIKLPAP.........
Slem-p1     ...PEIQRKNNFIINTGNNLTQLEKEKRLKLNKQKYGLELTYIKSLRLPKP.......VN
Otri-p1-4   KTGPEIQRKNNFIINTGNNLTQPEKEKRLRLNKQKYGLDLAFIKTLRLPKPES....ATG
Hgra-p1     KVGPQVIRKNNFIINTGDNQTAIERDARIKLNKQKYGLKADEIKALKLPKSLAGNGKAKG
Otri-p1-3   RQGAGVFRKNGCIINTGDNMTKVQIAKLRKKNSKIFGLTQPFVQSLHLPKAKVK......
Tthe-p1     K..QDVLRKNNCIINTGDNPTKEERKRRIEQNRKIYGLSAKQIAEIKLPKVK........
Pper-p1     K..VGVIRKFGCIINTGDNPNKNEKLERQRLYNQQFGLSKEEIKKLKIPYSY........
Ptet-p1     K..KNVFRKNNSIINTGDNMTQEERKKRIAENKKNFGISIADIALIKLPRYH........

             170       180       190       200       210       220
Otri-p1-1   CHLVEIFQIDKIMHPRSTLSTMDKIKHLIKLEDALKSKLEMIREGKRQQKFEQLQQKLKT
Otri-p1-2   CHLVEIFQIDKIMHPRSTLSTMDKIKHLIKLEDALKSKLEMIREGKRQQKFEQLQQKLKT
Otri-p1-5   CHLVEIFQIDKIMHPRSTLSTMDKIKHLIKLEDALKSKLEMIREGKRQQKFEQLQQKLKT
Slem-p1     NKKEVVLSIDQIFNAKSNFSVVEKLKHLEGLKISLEHKLYQIEKKRHIKEMRKLYKPLDQ
Otri-p1-4   GKREAILSMDQIFAQKSHFTVVEKLKHLEALKNALCSKLGKIERRRRNKELSKIYRPLGQ
Hgra-p1     GKDSLILSIDQIFSAKSNFSVIEKIQHLQRLQGALETKLSAIERRRLKKEEQKKYRAIKQ
Otri-p1-3   .EWLKVLTLDQILSAKSNFSTAEKIHYLKILENALERKLKKILR...LQELVSIYRPCNI
Tthe-p1     KHAPQYMTLEDIEN..EKFTNLEILTHLYNLKAEIVRRLAEQGETIAQPSIIKSLNNLNH
Pper-p1     KNLNQVMSLEQIQRNEKKLTNLQIIEEYQNLKNEITQRIKIFKQIKIEQQQQV.......
Ptet-p1     VNS.QTSFLSHDEIMEGNFTTVEKINNLCALKKDILRKLSKIENGELEEAFDDEIEQKRR

             230       240       250       260       270       280
Otri-p1-1   TEASGRGSVTRVQRQMSDLHLGSAHQNRNSDLDEENDQSVMIIDESQQQNLTPKGKAQTM
Otri-p1-2   TEASGRGSVTRVQRQMSDLHLGSAHQNRNSDLDEENDQSVMIIDESQQQNLTPKGKAQTM
Otri-p1-5   TEASGRGSVTRVQRQMSDLHLGSSHQNRNSDLDEENDESVMIIDESQQENLTPKGKAQAM
Slem-p1     AIVLAKIN........................GGQAYEYVETVDL...............
Otri-p1-4   LIVVQKN.........................ADDQYEFVDIIDENE.............
Hgra-p1     SIIVQRQV........................NLEGKEEYEFVGISNIA...........
Otri-p1-3   GIVVQKRLGS....................SIGDEYFEYVDCVKIEEKSVGNLDFALPNR
Tthe-p1     NLEQNQNSNSSTETKVTLEQSGKKKYKVLAIEETELQNGPIATNSQKKSINGKRKNNRKI
Pper-p1     ............................................................
Ptet-p1     YKKVMTDSDSSSA...............................................

             290            300       310       320       330       340
Otri-p1-1   LTN.....QTQTMKKQADDSRDEQHLPLISTSASVSNPSSTSKSSALKLNSMKQSDTAIA
Otri-p1-2   LTN.....QTQTMKKQADDSREEQHLPLNSTSASVSNPSSTSKSSALKLNSMKQSDTAIA
Otri-p1-5   LTHQKYNEVTQTMIKQGDDSRQQQHLPLDSTSASVSNPSSTSKSSTMKSNSMKQSETAIA
Slem-p1     ............................................................
Otri-p1-4   ............................................................
Hgra-p1     ............................................................
Otri-p1-3   NTD........STSLNEDFSFLDSTQKPQKLKAGSGRENKRKKMRDGLKDERAQRQSLME
Tthe-p1     NSDSEGNEEDISLEDIDSQESEINSEEIVEDDEEDEQIEEPSKIKKRKKNPEQESEEDDI
Pper-p1     ............................................................
Ptet-p1     ............................................................
```

```
                 350       360                                     370       380
                   .         .                                       .         .
Otri-p1-1   SMKPSSSGKKTKVDSSFVSKQSNQQ......................STSYSETNVDTQN
Otri-p1-2   SMKPSSSGKKTKVDSSFVSKQSNQQSTGPIQKQAHQQNLDRNRSELGSTFAQQTNVDTQN
Otri-p1-5   SMKPSSIGKKTKVDSSFVTKQSNQQSTAPIQKQAHQQNLDRNRSELGSTFAQQASVDTQN
Slem-p1     ............................................................
Otri-p1-4   ............................................................
Hgra-p1     ............................................................
Otri-p1-3   ALDEQEFDETKFQDSDGEMPDLNM....................................
Tthe-p1     EEDQEEDELVVNEEEIFEDDDDDEDNQDSSEDDDDDED......................
Pper-p1     ............................................................
Ptet-p1     ............................................................


                   390       400       410
                     .         .         .
Otri-p1-1   SNNQGTSTASGNFISQSDDEEALMPKLKRRRVEDSE
Otri-p1-2   SNNQGTSTASGNFISQSDDEEALMPKLKRRRVKDSE
Otri-p1-5   SNNQGTSTASGNFISQSDDEEALMPKLKRRRVEDSE
Slem-p1     ....................................
Otri-p1-4   ....................................
Hgra-p1     ....................................
Otri-p1-3   ....................................
Tthe-p1     ....................................
Pper-p1     ....................................
Ptet-p1     ....................................
```
